# The Environmental Risks of neonicotinoid pesticides: a review of the evidence post-2013

**DOI:** 10.1101/098897

**Authors:** Thomas James Wood, Dave Goulson

**Affiliations:** The University of Sussex

## Abstract

Neonicotinoid pesticides were first introduced in the mid-1990s and since then their use has grown rapidly so that they have become the most widely used class of insecticides in the world, with the majority being used as seed coatings. Neonicotinoids are water-soluble, and so a small quantity applied to a seed will dissolve when in contact with water in the soil and be taken up by the roots of the developing plant. Once inside the plant it becomes systemic and is found in vascular tissues and foliage, providing protection against herbivorous insects. This prophylactic use of neonicotinoids has become extremely widespread on a wide range of arable crops across much of the developed world.

However, only approximately 5% of the neonicotinoid active ingredient is taken up by crop plants and most instead disperses into the wider environment. Since the mid-2000s numerous studies have raised concerns that neonicotinoids may be having a negative effect on non-target organisms. In particular, neonicotinoids were associated with mass poisoning events of honeybees and were shown to have serious negative effects on honeybee and bumblebee fitness when consumed. In response to this growing body of evidence, the European Food Safety Authority (EFSA) was commissioned to produce risk assessments for the use of clothianidin, imidacloprid and thiamethoxam and their impact on bees. These risk assessments, published in January 2013, conclude that the use of these compounds on certain flowering crops poses a high ris
k to bees. On the basis of these findings, the European Union adopted a partial ban on these substances in May 2013 which came into force on 1^st^ December 2013.

The purpose of this review is to collate and summarise scientific evidence published since 2013 that investigates the impact of neonicotinoids on non-target organisms and to bring it into one place to aid informed decision making. Due to international concern over the unintended impacts of neonicotinoids on wildlife, this topic has received a great deal of scientific attention in this three year period. As the restrictions were put in place because of the risk neonicotinoids pose to bees, much of the recent research work has naturally focussed on this group.

**Risks to bees:** Broadly, the EFSA risk assessments addressed risks of exposure to bees from neonicotinoids through various routes and the direct lethal and sublethal impact of neonicotinoid exposure. New scientific evidence is available in all of these areas, and it is possible to comment on the change in the scientific evidence since 2013 compared to the EFSA reports. This process is not meant to be a formal assessment of the risk posed by neonicotinoids in the manner of that conducted by EFSA. Instead it aims to summarise how the new evidence has changed our understanding of the likely risks to bees; is it lower, similar or greater than the risk perceived in 2013. With reference to the EFSA 2013 risk assessments baseline, advances in each considered area and their impact on the original assessment can be summarised thus:

- *Risk of exposure from pollen and nectar of treated flowering crops.* The EFSA reports calculated typical exposure from flowering crops treated with neonicotinoids as seed dressings. Considerably more data are now available in this area, with new studies broadly supporting the calculated exposure values. For bees, flowering crops pose a **Risk Unchanged** to that reported by EFSA 2013a.
- *Risk from non-flowering crops and cropping stages prior to flowering.* Non-flowering crops were considered to pose no risk to bees. No new studies have demonstrated that these non-flowering crops pose a direct risk to bees. They remain a **Risk Unchanged**.
- *Risk of exposure from the drilling of treated seed and subsequent dust drift.* Despite modification in seed drilling technology, available studies suggest that dust drift continues to occur, and that dust drift still represents a source of acute exposure and so is best considered a **Risk Unchanged**.
- *Risk of exposure from guttation fluid.* Based on available evidence this was considered a low-risk exposure path by EFSA 2013a. New data have not changed this position and so it remains a **Risk Unchanged**.
- *Risk of exposure from and uptake of neonicotinoids in non-crop plants.* Uptake of neonicotinoids by non-target plants was considered likely to be negligible, though a data gap was identified. Many studies have since been published demonstrating extensive uptake of neonicotinoids and their presence in the pollen, nectar and foliage of wild plants. Bees collecting pollen from neonicotinoid-treated crops can generally be expected to be exposed to the highest neonicotinoid concentrations, but non-trivial quantities of neonicotinoids are also present in pollen and nectar collected from wild plants, and this source of exposure may be much more prolonged than the flowering period of the crop. Exposure from non-target plants clearly represents a **Greater Risk**.
- *Risk of exposure from succeeding crops.* A data gap was identified for this issue. Few studies have explicitly investigated this, but this area does represent some level of risk as neonicotinoids are now known to have the potential to persist for years in soil, and can be detected in crops multiple years after the last known application. However, as few data exist this is currently considered a **Risk Unchanged**.
- *Direct lethality of neonicotinoids to adult bees.* Additional studies on toxicity to honeybees have supported the values calculated by EFSA. More data have been produced on neonicotinoid toxicity for wild bee species and meta-analyses suggest a broadly similar response. Reference to individual species is important but neonicotinoid lethality should be broadly considered a **Risk Unchanged**.
- *Sublethal effects of neonicotinoids on wild bees.* Consideration of sublethal effects by EFSA was limited as there is no agreed testing methodology for the assessment of such effects. A data gap was identified. Exposure to neonicotinoid-treated flowering crops has been shown to have significant negative effects on free flying wild bees under field conditions and some laboratory studies continue to demonstrate negative effects on bee foraging ability and fitness using field-realistic neonicotinoid concentrations. **Greater Risk**.

Within this context, research produced since 2013 suggest that neonicotinoids pose a similar to greater risk to wild and managed bees, compared to the state of play in 2013. Given that the initial 2013 risk assessment was sufficient to impose a partial ban on the use of neonicotinoids on flowering crops, and given that new evidence either confirms or enhances evidence of risk to bees, it is logical to conclude that the current scientific evidence supports the extension of the moratorium, and that the extension of the partial ban to other uses of neonicotinoids should be considered.

**Broader risks to environmental health:** In addition to work on bees, our scientific understanding has also been improved in the following areas which were not previously considered by EFSA:

- Non-flowering crops treated with neonicotinoids can pose a risk to non-target organisms through increasing mortality in beneficial predator populations.
- Neonicotinoids can persist in agricultural soils for several years, leading to chronic contamination and, in some instances, accumulation over time.
- Neonicotinoids continue to be found in a wide range of different waterways including ditches, puddles, ponds, mountain streams, rivers, temporary wetlands, snowmelt, groundwater and in outflow from water processing plants.
- Reviews of the sensitivity of aquatic organisms to neonicotinoids show that many aquatic insect species are several orders of magnitude more sensitive to these compounds than the traditional model organisms used in regulatory assessments for pesticide use.
- Neonicotinoids have been shown to be present in the pollen, nectar and foliage of non-crop plants adjacent to agricultural fields. This ranges from herbaceous annual weeds to perennial woody vegetation. We would thus expect non-target herbivorous insects and non-bee pollinators inhabiting field margins and hedgerows to be exposed to neonicotinoids. Of particular concern, this includes some plants sown adjacent to agricultural fields specifically for the purposes of pollinator conservation.
- Correlational studies have suggested a negative link between neonicotinoid usage in agricultural areas and population metrics for butterflies, bees and insectivorous birds in three different countries.

Overall, this recent work on neonicotinoids continues to improve our understanding of how these compounds move through and persist in the wider environment. These water soluble compounds are not restricted to agricultural crops, instead permeating most parts of the agricultural environments in which they are used and in some cases reaching further afield via waterways and runoff water. Field-realistic laboratory experiments and field trials continue to demonstrate that traces of residual neonicotinoids can have a mixture of lethal and sublethal effects on a wide range of taxa. Susceptibility varies tremendously between different taxa across many orders of magnitude, with some showing a negative response at parts per billion with others show no such effects at many thousands of parts per billion. Relative to the risk assessments produced in 2013 for clothianidin, imidacloprid and thiamethoxam which focussed on their effects on bees, new research strengthens arguments for the imposition of a moratorium, in particular because it has become evident that they pose significant risks to many non-target organisms, not just bees. Given the improvement in scientific knowledge of how neonicotinoids move into the wider environment from all crop types, a discussion of the risks posed by their use on non-flowering crops and in non-agricultural areas is urgently needed.

## 1. INTRODUCTION AND STATE OF PLAY

Neonicotinoid pesticides were first introduced in the 1990s and since then their use has grown rapidly to become the most widely used class of insecticide in the world. This increase in popularity has largely occurred from the early 2000s onwards (Figure 1). This use has largely been driven by the adoption of seed treatments. Neonicotinoids are water-soluble, and so a small quantity applied to a seed will dissolve when in contact with water and be taken up by the roots of the developing plant. Once inside the plant it becomes systemic and is found in vascular tissues and foliage, providing protection against herbivorous insects. This prophylactic use of neonicotinoids has become extremely widespread – for example, between 79–100% of maize hectares in the United States in 2011 were treated with a neonicotinoid seed dressing (Douglas and Tooker 2015).

**Figure 1.**
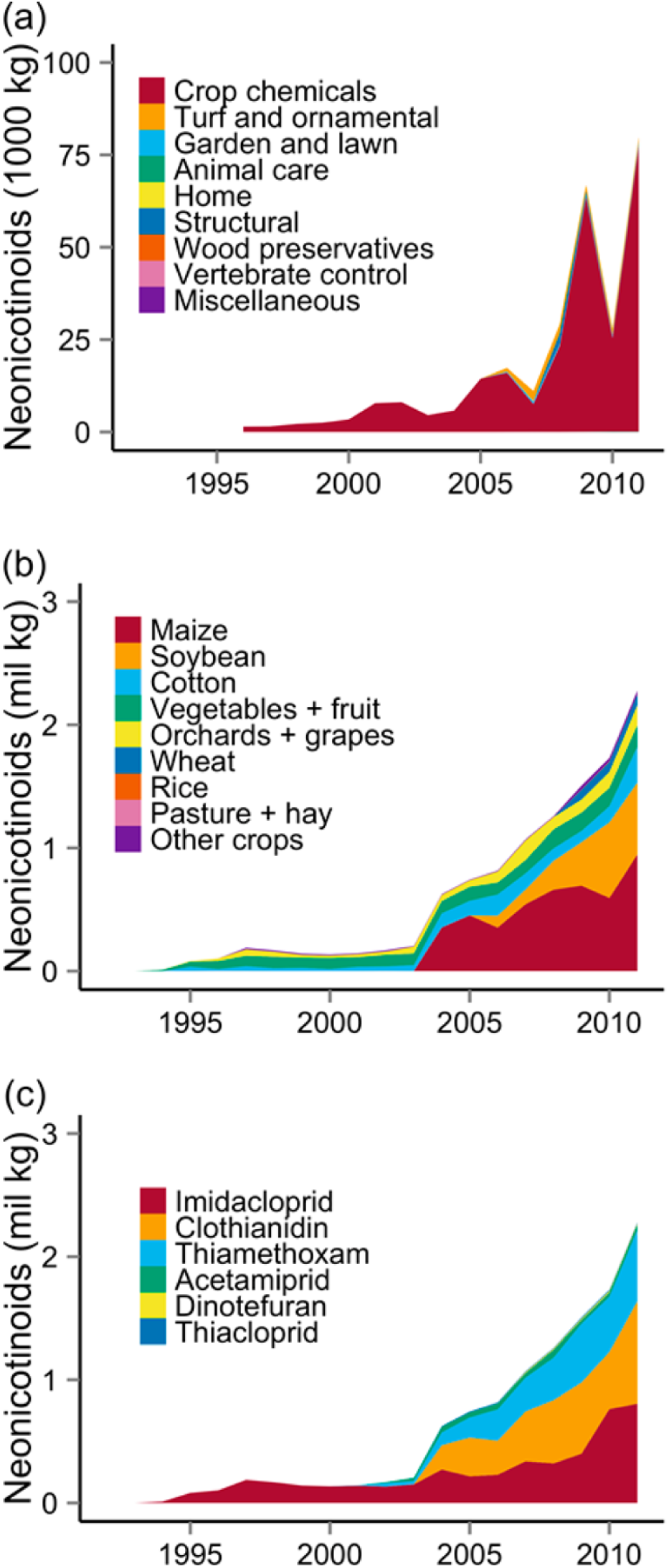
Neonicotinoid sales by (a) product type, (b) use by crop and (c) active ingredient, from 1992 to 2011. Data on use (a) is based on sales data from Minnesota. Data on crops and active ingredients are for the entire U.S., from United States Geological Survey. y-axes represent mass of neonicotinoid active ingredient in thousands or millions of kg. Reproduced from Douglas and Tooker (2015)

However, only approximately 5% of the neonicotinoid active ingredient is taken up by crop plants and most instead disperses into the wider environment. In recent years numerous authors have raised concerns about the impact neonicotinoids may have on non-target organisms. Neonicotinoids released in dust abraded by seed drilling machinery were implicated in mass poisonings of honeybees in Germany and Italy (Pistorius et al. 2009; Bortolotti et al. 2009), neonicotinoids were found in agricultural soils (Bonmatin *et al.* 2005) and also in the pollen and nectar of treated crops (Bonmatin *et al.* 2007). In 2012, two high profile studies were published that showed exposure to neonicotinoids in pollen and nectar could have serious effects on honeybee navigation and mortality (Henry *et al.* 2012) and bumblebee colony development and queen production (Whitehorn *et al.* 2012) . In response to the growing body of work the European Food Safety Authority (EFSA), the body with regulatory oversight for agricultural chemicals, was commissioned to produce a risk assessment on the three most widely used agricultural neonicotinoids (clothianidin, imidacloprid and thiamethoxam) and the risk that they posed to bees (EFSA 2013a; 2013b; 2013c). On the basis of the available evidence EFSA recommended a moratorium on the use of neonicotinoids on treated crops which was accepted and implemented by the European Commission at the end of 2013.

This moratorium is due to conclude shortly. One of the specified objectives was to allow further research on the impact of neonicotinoids on bees in order to inform subsequent regulatory decisions. Since 2013, a great number of studies have been published that consider the impact of neonicotinoids on bees and also a wide range of other non-target taxa. Many large reviews of neonicotinoids impacts on non-target organisms have also been published, for example Nuyttens et al. (2013) on neonicotinoid contaminated dust, Godfray *et al.* 2014; 2015) on the risks neonicotinoids pose to pollinators, Bonmatin *et al.* (2015) on environmental fate of and exposure to neonicotinoids, Pisa *et al.* (2015) and Gibbons *et al.* (2015) on the impacts of neonicotinoids on non-target terrestris organisms and Morrissey *et al.* (2015) on contamination of aquatic ecosystems with neonicotinoids and their impact on aquatic organisms, to name a few.

The purpose of this review is to consider the scientific evidence published since 2013 that covers the impact of neonicotinoids on wild non-target organisms (therefore excluding the domesticated honeybee) and to bring it together into one place to aid informed decision making. It is not a formal risk assessment, though comparisons will be made with the knowledge base used in the EFSA risk assessments specifically and that which was known in 2013 more generally. The findings will be of interest to those considering the wider impact of neonicotinoid pesticide use when assessing their future use in agricultural environments.

## 2. EVIDENCE FOR EXPOSURE TO NEONICOTINOID PESTICIDES

### 2.1 Risk of exposure for non-target organisms from neonicotinoids applied directly to crops

Due to their systemic nature, neonicotinoids applied to crops by any application method (e.g. seed dressing, foliar spray, soil drench) will be taken up by crop tissues and can subsequently be found in all parts of the treated plant (Simon-Delso *et al.* 2015). The EFSA 2013a; 2013b; 2013c) reports identify and discuss a number of exposure pathways through which bees can be exposed to neonicotinoids, where the risk of exposure is dependent on application rate, application type and crop type. However, knowledge about the extent and significance of these pathways was poor. Since then, a large number of studies have been published further documenting neonicotinoid exposure from treated crops. Important reviews include Nuyttens et al. (2013), Godfray *et al.* (2014), Long and Krupke (2015) and Bonmatin *et al.* (2015).

#### 2.1.1 Risk of exposure from pollen and nectar of treated flowering crops

Using data from 30 (clothianidin), 16 (thiamethoxam) and 29 (imidacloprid) outdoor studies and known authorised application rates, EFSA 2013a; 2013b; 2013c) calculated expected residue rates in pollen and nectar of the studied crops (Table 1). Levels are variable but all are within one order of magnitude. Levels in pollen are consistently higher than levels in nectar. Godfray *et al.* (2014) reviewed 20 published studies to calculate an arithmetic mean maximum level of 1.9 ppb for nectar and 6.1 ppb for pollen in treated crops, in line with the EFSA findings.

**Table 1.** Summary of expected residues in pollen and nectar of various neonicotinoid-treated flowering crops calculated by EFSA from the review of outdoor field trials. No nectar values are available for maize as this plant does not produce nectar. Blanks are where no minimum values were stated

| Crop | Pesticide | Residues in pollen (ng/g) |  | Residues in nectar (ng/g) |  |
| --- | --- | --- | --- | --- | --- |
|  |  | Minimum | Maximum | Minimum | Maximum |
| Oilseed rape | Clothianidin | 5.95 | 19.04 | 5 | 16 |
| Sunflower | Clothianidin |  | 3.29 |  | 0.324 |
| Maize | Clothianidin | 7.38 | 36.88 | *n/a* | *n/a* |
| Oilseed rape | Imidacloprid | 1.56 | 8.19 | 1.59 | 8.35 |
| Sunflower | Imidacloprid |  | 3.9 |  | 1.9 |
| Maize | Imidacloprid | 3.02 | 15.01 | *n/a* | *n/a* |
| Cotton | Imidacloprid | 3.45 | 4.6 | 3.45 | 4.6 |
| Oilseed rape | Thiamethoxam | 4.592 | 19.29 | 0.648 | 2.72 |
| Sunflower | Thiamethoxam | 2.378 | 3.02 | 0.59 | 0.75 |
| Maize | Thiamethoxam | 13.419 | 21.513 | *n/a* | *n/a* |

Since 2014 a number of studies have been published which report neonicotinoid concentrations in the pollen and nectar of neonicotinoid-treated flowering crops. These results have been approximately in line with the concentrations reported by EFSA and Godfray *et al.* In oilseed rape treated with thiamethoxam, Botías *et al.* (2015) found average concentrations of 3.26 ng/g of thiamethoxam, 2.27 ng/g of clothianidin and 1.68 ng/g of thiacloprid in the pollen. Oilseed rape nectar contained similar average concentrations of 3.20 ng/g of thiamethoxam, 2.18 ng/g of clothianidin and 0.26 ng/g of thiacloprid. Xu *et al.* (2016) found average levels of clothianidin in oilseed rape of 0.6 ng/g. No pollen samples were taken. In maize pollen, Stewart *et al.* (2014) found average thiamethoxam and clothianidin levels between the limit of detection (LOD) of 1 ng/g to 5.9 ng/g across a range of seed treatments. Xu et al. (2016) found average clothianidin concentration of 1.8 ng/g in maize pollen. Additionally, Stewart et al. (2014) found no neonicotinoid residues in soybean flowers or cotton nectar.

Several studies published since 2013 have used free flying bees to experimentally demonstrate that proximity to treated flowering crops increases their exposure to neonicotinoids (Table 2). Using honeybees, neonicotinoid concentrations in pollen taken from foragers returning to nests placed next to untreated flowering crops ranged from 0–0.24 ng/g compared to pollen from nests next to treated flowering crops which ranged from 0.84–13.9 ng/g. There have been fewer studies of bumblebees and hence the sample size is much smaller, with concentrations of neonicotinoids in pollen from untreated areas ranging from <0.1-<0.3 ng/g compared to 0.4–0.88 ng/g for nests placed next to treated areas. The only available study looking at solitary bee collected pollen found *Osmia bicornis* collecting <0.3 ng/g in untreated areas and 0.88 ng/g in treated areas. Similar trends are found in the nectar results, though fewer studies are available. Rolke *et al.* (2016) found neonicotinoid concentrations of 0.68–0.77 ng/ml in honeybee collected nectar samples from apiaries adjacent to neonicotinoid-treated oilseed rape, compared to <0.3 ng/ml from apiaries adjacent to untreated oilseed rape. However, Rundlöf *et al.* (2015) found concentrations of 5.4 ng/ml in bumblebee collected nectar and 10.3 ng/ml in honeybee collected nectar taken from bees originating from nests placed adjacent to treated oilseed rape compared to 0–0.1 ng/ml from bees from nests adjacent to untreated oilseed rape.

**Table 2.** Summary of studies published since 2013 that document neonicotinoid residues in pollen and nectar collected by free flying bees at sites adjacent to treated and untreated flowering crops. Results for samples collected at treated sites are highlighted in bold. SS = spring-sown, WS = winter-sown, US = unclear sowing date

| Species | Sample type | Samples collected | Nest location | Mean total neonicotinoid concentration (ng/ml or ng/g) | Reference |
| --- | --- | --- | --- | --- | --- |
| *Apis mellifera* | Nectar | 2005-2009 (dates unknown) | Adjacent to untreated US OSR fields | <1 (*limit of quantification*) | Pilling *et al.* (2013) |
| *Apis mellifera* | Nectar | 2005-2009 (dates unknown) | Adjacent to treated US OSR fields | **0.7-2.4** (range of reported median values) | Pilling *et al.* (2013) |
| *Apis mellifera* | Nectar | 6<sup>th</sup> May 2014 | Adjacent to untreated WS OSR fields | <0.3 (*limit of detection*) | Rolke *et al.* (2016) |
| *Apis mellifera* | Nectar | 6<sup>th</sup> May 2014 | Adjacent to treated WS OSR fields | **0.68** | Rolke *et al.* (2016) |
| *Apis mellifera* | Nectar | 10<sup>th</sup>-14<sup>th</sup> May 2014 | Adjacent to untreated WS OSR fields | <0.3 (*limit of detection*) | Rolke *et al.* (2016) |
| *Apis mellifera* | Nectar | 10<sup>th</sup>-14<sup>th</sup> May 2014 | Adjacent to treated WS OSR fields | **0.77** | Rolke *et al.* (2016) |
| *Apis mellifera* | Nectar | June 2013 (peak OSR flowering) | Adjacent to untreated SS OSR fields | 0.1 | Rundlöf *et al.* (2015) |
| *Apis mellifera* | Nectar | June 2013 (peak OSR flowering) | Adjacent to treated SS OSR fields | **10.3** | Rundlöf *et al.* (2015) |
| *Bombus terrestris* | Nectar | June 2013 (peak OSR flowering) | Adjacent to untreated SS OSR fields | 0 | Rundlöf *et al.* (2015) |
| *Bombus terrestris* | Nectar | June 2013 (peak OSR flowering) | Adjacent to treated SS OSR fields | **5.4** | Rundlöf *et al.* (2015) |
| *Apis mellifera* | Pollen | 2005-2009 (dates unknown) | Adjacent to untreated maize fields | <1 (*limit of quantification*) | Pilling *et al.* (2013) |
| *Apis mellifera* | Pollen | 2005-2009 (dates unknown) | Adjacent to treated maize fields | **1-7** (range of reported median values) | Pilling *et al.* (2013) |
| *Apis mellifera* | Pollen | 2005-2009 (dates unknown) | Adjacent to untreated US OSR fields | <1 (*limit of quantification*) | Pilling *et al.* (2013) |
| *Apis mellifera* | Pollen | 2005-2009 (dates unknown) | Adjacent to treated US OSR fields | **<1-3.5** (range of reported median values) | Pilling *et al.* (2013) |
| *Apis mellifera* | Pollen | First two weeks of July 2012 | Located in untreated SS OSR fields | 0.24 | Cutler *et al.* (2014) |
| *Apis mellifera* | Pollen | First two weeks of July 2012 | Located in treated SS OSR fields | **0.84** | Cutler *et al.* (2014) |
| *Apis mellifera* | Pollen | June 2013 (peak OSR flowering) | Adjacent to untreated WS OSR fields | <0.5 (*limit of detection*) | Rundlöf *et al.* (2015) |
| *Apis mellifera* | Pollen | June 2013 (peak OSR flowering) | Adjacent to treated WS OSR fields | **13.9** | Rundlöf *et al.* (2015) |
| *Apis mellifera* | Pollen | May to September 2011 | Non-agricultural area | 0.047 | Long and Krupke (2016) |
| *Apis mellifera* | Pollen | May to September 2011 | Adjacent to untreated maize fields | 0.078 | Long and Krupke (2016) |
| *Apis mellifera* | Pollen | May to September 2011 | Adjacent to treated maize fields | **0.176** | Long and Krupke (2016) |
| *Apis mellifera* | Pollen | 6<sup>th</sup> May 2014 | Adjacent to untreated WS OSR fields | <0.3 (*limit of detection*) | Rolke *et al.* (2016) |
| *Apis mellifera* | Pollen | 6<sup>th</sup> May 2014 | Adjacent to treated WS OSR fields | **0.50** | Rolke *et al.* (2016) |
| *Apis mellifera* | Pollen | 10<sup>th</sup>-14<sup>th</sup> May 2014 | Adjacent to untreated WS OSR fields | <0.3 (*limit of detection*) | Rolke *et al.* (2016) |
| *Apis mellifera* | Pollen | 10<sup>th</sup>-14<sup>th</sup> May 2014 | Adjacent to treated WS OSR fields | **0.97** | Rolke *et al.* (2016) |
| *Bombus terrestris* | Pollen | 10<sup>th</sup> May 2014 | Adjacent to untreated WS OSR fields | <0.3 (*limit of detection*) | Rolke *et al.* (2016) |
| *Bombus terrestris* | Pollen | 10<sup>th</sup> May 2014 | Adjacent to treated WS OSR fields | **0.88** | Rolke *et al.* (2016) |
| *Bombus impatiens* | Pollen | July to August 2013 | Adjacent to untreated maize fields | <0.1 (*limit of detection*) | Cutler and Scott-Dupree (2014) |
| *Bombus impatiens* | Pollen | July to August 2013 | Adjacent to treated maize fields | **0.4** | Cutler and Scott-Dupree (2014) |
| *Osmia bicornis* | Pollen | 14<sup>th</sup> May 2014 | Adjacent to untreated WS OSR fields | <0.3 (*limit of detection*) | Rolke *et al.* (2016) |
| *Osmia bicornis* | Pollen | 14<sup>th</sup> May 2014 | Adjacent to treated WS OSR fields | **0.88** | Rolke *et al.* (2016) |

This level of variation of up to one order of magnitude in neonicotinoid concentrations found in bee collected pollen and nectar in different studies is substantial. The detected levels in pollen and nectar presumably depend significantly on the dose and mode of treatment, the studied crop, the season, the location, the soil type, the weather, time of day samples are collected, and so on. Even different crop varieties can result in significant variation in the residue content of pollen and nectar (Bonmatin *et al.* 2015). Because pollen samples taken from a series of bees will be from a mixture of different plants, most of which will not be crop plants, the neonicotinoid residues in crop pollen will be diluted by untreated, non-crop pollen. However, for the reported studies, the higher neonicotinoid concentrations are within an order of magnitude of the 6.1 ng/g in pollen and 1.9 ng/ml in nectar values calculated by Godfray *et al.* (2014). Additionally, in all cases, the concentrations of neonicotinoids in pollen and nectar were higher at sites adjacent to neonicotinoid-treated flowering crops than at sites adjacent to untreated crops. The available evidence shows that proximity to treated flowering crops increases the exposure of bees to neonicotinoid pesticides. The recent evidence for concentrations found in flowering crops is approximately in line with the levels reported by EFSA 2013a; 2013b; 2013c).

#### 2.1.2 Risk from non-flowering crops and cropping stages prior to flowering

The EFSA studies state that some of the crops on which clothianidin is authorised as a seed-dressing do not flower, are harvested before flowering, or do not produce nectar or pollen, and therefore these crops will not pose any risk to bees via this route of exposure. Whilst non-flowering crops are clearly not a source of exposure through produced pollen and nectar, they do represent a source of neonicotinoids that can dissipate into the wider environment (discussed in Section 2.2). Additionally, treated crops of any type represent additional pathways of neonicotinoid exposure to other organisms.

Depending on crop species and consequent seed size, neonicotinoid-treated seeds contain between 0.2–1 mg of active ingredient per seed (Goulson 2013). For a granivorous grey partridge weighing 390 g Goulson calculated that it would need to consume around five maize seeds, six sugar beet seeds or 32 oilseed rape seeds to receive a nominal LD_50_. Based on US Environmental Protection Agency estimates that around 1% of sown seed is accessible to foraging vertebrates at recommended sowing densities, Goulson calculated that sufficient accessible treated seed would be present to deliver a LD_50_ to ~100 partridges per hectare sown with maize or oilseed rape. Given that grey partridges typically consume around 25 g of seed a day there is the clear potential for ingestion of neonicotinoids by granivorous animals, specifically birds and mammals. However, whilst some experimental studies have been conducted to investigate mortality and sublethal effects of treated seeds on birds (see Section 3.5), no studies are available that demonstrate consumption of treated seed by farmland birds under field conditions or quantify relative consumption of treated versus untreated seed to better understand total exposure via this route.

In addition to insect herbivores, developing seedlings treated with neonicotinoids are predated by molluscan herbivores. Because neonicotinoids have relatively low efficacy against molluscs, Douglas *et al.* (2015) investigated neonicotinoid residues in the slug *Deroceras reticulatum,* a major agricultural pest, using neonicotinoid seed-treated soybean in both laboratory and field studies. Total neonicotinoid concentrations from samples of field collected slugs feeding on treated soybean were as high as 500 ng/g with average levels over 100 ng/g after 12 days of feeding. No neonicotinoids were detected in slugs feeding on untreated control plants. After 169 days, no neonicotinoids were detected in either control or treated slugs. In the laboratory, slugs consuming soybean seedlings incurred low mortality of between 6–15% depending on the strength of the seed treatment. In laboratory experiments, slugs were exposed to the ground beetle *Chlaenius tricolor* after feeding on soybean. *C. tricolor* is a typical predatory beetle found in agro-ecosystems and is known to be an important predator of slugs. For beetles that consumed slugs, 61.5% (n=16/26) of those from the neonicotinoid treatment subsequently showed signs of impairment compared to none of those in the control treatment (n=0/28). Of the 16 that showed impairment, seven subsequently died. This study is also discussed in Section 3.3. A similar result was found by Szczepaniec *et al.* (2011) who found that the application of imidacloprid to elm trees caused an outbreak of spider mites *Tetranychus schoenei*. This increase was as a result of a reduction in the density of their predators which incurred increased mortality after ingesting imidacloprid-containing prey items. Many beneficial predatory invertebrates feed on pests of crops known to be treated with neonicotinoids, but to date no other studies have assessed whether neonicotinoids are transmitted to these predators through direct consumption of crop pests in agro-ecosystems.

Additionally, flowering crops in a non-flowering stage can also pose a potential threat to natural enemy populations. The soybean aphid parasitoid wasp *Aphelinus certus* is an important parasite of the soybean aphid *Aphis glycines*. Frewin *et al.* (2014) gave *A. certus* access to laboratory populations of aphids feeding on control and neonicotinoid-treated soybean plants. *A. certus* parasitised a significantly smaller proportion of aphids on treated plants than on untreated plants. Frewin *et al.* hypothesise two potential reasons for this effect – firstly that exposure to neonicotinoid residues within aphid hosts may have increased mortality of the immature parasitoid or the parasitism combined with residues may have increased aphid mortality. Secondly, *A. certus* may avoid parasitising pesticide-poisoned aphids. *Aphelinus* species are known to use internal cues to determine host suitability, and it is possible that they may use stress- or immune-related aphid hormones to judge host suitability. Given that a key part of biological control of insect pests using parasitic wasps is to increase the parasitoid abundance early in the season, the reduction in the parasitism rate caused by neonicotinoid seed-treatment could potentially impair the ability of *A. certus* to control soybean aphid.

Non-flowering neonicotinoid crops present possible exposure routes through direct consumption of treated seed or consumption of seedling plants that may result in the transmission of neonicotinoids to higher trophic levels, including beneficial insects that offer a level of pest control through predatory behaviour. As the EFSA reports did not consider the impact of neonicotinoids on non-bees, no comparison can be made here.

#### 2.1.3 Risk of exposure from the drilling of treated seed and subsequent dust drift

Numerous studies (12 listed by Godfray *et al.* 2014) prior to 2013 identified that neonicotinoids present in seed dressings can be mechanically abraded during the drilling process and can subsequently be emitted as dust. This dust can contain very high levels of neonicotinoids, up to 240,000 ng/g under certain conditions (see the review by Nuyttens *et al.* 2013). Acute contact with this dust can in certain cases result in the mass poisoning of honeybees (e.g. Pistorius *et al.* 2009; Bortolotti *et al.* 2009). Concentrations of neonicotinoids in dust created during sowing and the total volume released into the air depend on application rate, seed type, seed treatment quality (including additions such as talcum powder), seed drilling technology and environmental conditions. Girolami *et al.* (2013) demonstrated that the dust cloud created by seed drills is an ellipsoidal shape approximately 20 m in diameter. Using cage experiments, a single pass of a drilling machine was sufficient to kill all honeybees present. The use of tubes designed to direct exhaust air towards the ground did not substantially increase bee survival rate. Neonicotinoid concentrations of up to 4000 ng/g were detected in honeybees with an average concentration of 300 ng/g. Similar concentrations were detected in bees exposed to both unmodified and modified drills.

On the basis of the available evidence, the EFSA reports (2013a; 2013b; 2013c) concluded that maize produces the highest dust drift deposition, while for sugar beet, oilseed rape and barley seeds the dust drift deposition was very limited. No information was available for other crops, and given that seed type is an important factor determining neonicotinoid release, extrapolation to other crops is highly uncertain. A high acute risk was not excluded for bees foraging or flying in adjacent crops during the sowing of maize, oilseed rape, and cereals. In practice, this assessment indicates that forager honeybees or other pollinators flying adjacent to the crop are at high risk (e.g. via direct contact to dust) and may be able to carry considerable residues back to the hive (for social bees). Bees present further away or foraging upwind during the sowing will be considerably less exposed. The reports conclude that the aforementioned assessments do not assess potential risk to honeybees from sublethal effects of dust exposure. No information on neonicotinoid residues in nectar in the adjacent vegetation following dust drift was available.

In recent years, various types of improved seed drills have been adopted that direct air from the drills towards the soil, reducing the dust drift effect by up to 95% (see Manzone *et al.* 2015). Air deflectors have become mandatory for certain products in the Netherlands, France, Belgium and Germany (Godfray *et al.* 2014). Bonmatin *et al.* (2015) and Long and Krupke (2015) reviewed existing literature on the exposure of pollinators and other non-target organisms to contaminated dust from seed drilling machines, predominantly covering pre-April 2013 literature. The authors conclude that despite attention by regulators they consider dust drift to be a likely cause of environmental neonicotinoid contamination, in particular when best practice is not followed.

Recent studies continue to detect neonicotinoids in the tissues of wildflowers surrounding agricultural fields immediately after planting. Stewart *et al.* (2014) detected average neonicotinoid concentrations of 9.6 ng/g in whole wildflowers collected from field margins adjacent to fields planted with maize (n=18), cotton (n=18) and soybean (n=13). The samples were collected a few days after sowing (typically within three days), with the highest concentration of 257 ng/g collected adjacent to a maize field sown the previous day with thiamethoxam-treated seed. Detailed data on concentrations adjacent to each crop type are not available. No samples were taken from vegetation adjacent to crops sown without a neonicotinoid seed dressing. Rundlöf *et al.* (2015) collected flowers and leaves from wild plants growing adjacent to treated and untreated oilseed rape fields two days after sowing. Adjacent to the treated fields neonicotinoid concentrations were lower than in the previous study at 1.2 ng/g, but this was higher than the control fields where no neonicotinoids were detected. This is in line with previous findings that suggest a lower contamination risk from dust originating from oilseed rape seeds than for maize seeds.

#### 2.1.4 Risk of exposure from guttation fluid

Some plants secrete small volumes of liquid (xylem sap) at the tips of leaves or other marginal areas, often referred to as guttation droplets. Six published studies and an EFSA review found extremely high neonicotinoid concentrations in guttation droplets of up to 4–5 orders of magnitude greater than those found in nectar, particularly when plants are young (see Godfray *et al.* 2014). Using a clothianidin concentration of 717,000 ng/g and an acute oral toxicity of 3.8 ng/bee for clothianidin (see Section 3.1.1), EFSA (2013a) calculated that a honeybee would only need to consume 0.005 μl to receive an LD_50_. Given that honeybee workers can carry between 1.4–2.7 ml of water a day, there is the clear potential for lethal exposure via this route. The risk assessments for thiamethoxam and imidacloprid were similar (EFSA 2013b; 2013c). However, on the basis of experimental trials, the EFSA reports conclude that whilst guttation droplets were frequently produced, honeybees were rarely seen collecting water from them and therefore the risk should be considered low.

Few studies have looked at neonicotinoid exposure via guttation droplets since 2013. In the one available study, Reetz *et al.* (2015) assessed thiamethoxam concentrations in oilseed rape guttation droplets and measured residues in individual honeybee honey-sacs. The authors note that targeted observations of water-foraging honeybees in the field are nearly impossible, and so returning honeybees from apiaries placed out adjacent to treated oilseed rape crops were instead collected in the autumns of 2010 and 2011 when seedling oilseed rape crops were producing guttation droplets. Oilseed rape produced guttation droplets containing between 70–130 ng/ml clothianidin at the cotyledon stage. Out of 436 honey-sacs, neonicotinoids were only detected in 62 samples at concentrations between 0.1–0.95 ng/ml. However, because there was no behavioural observation it is not possible to state the providence of this contamination with certainty; neonicotinoids are also present in waterbodies and the nectar of wild flowers (see Section 2.2). As such, there is still little evidence documenting the extent to which honeybees or other insects collect or are otherwise exposed to neonicotinoids through contact with guttation droplets.

### 2.2 Risk of exposure for non-target organisms from neonicotinoids persisting in the wider environment

In identifying routes of exposure for honeybees the EFSA reports discussed the possibility of neonicotinoid residues in flowering arable weeds growing in fields with treated crops. This route of exposure was considered to be negligible as weeds would not be present in the field when the crop is sown and considerable uptake via weed plant roots was considered to be unlikely as the substance is concentrated around the treated seed. However, the reports note that potential uptake into flowering weeds cannot be ruled out for granular neonicotinoid applications, highlighting a data gap for this issue.

The persistence of neonicotinoids in soil, water and in wild plants is of potentially serious concern. If these pesticides are able to move into habitats surrounding agricultural fields the range of organisms that they could affect is much greater than simply crop-visiting invertebrates. If these pesticides last for extended periods in the wider environment then neonicotinoid exposure may be chronic, rather than an acute exposure associated with the sowing of treated seeds.

Since April 2013 much empirical data has been produced documenting the fate of residual neonicotinoids in the wider environment after application. Key review publications are Goulson (2013) , Bonmatin *et al.* (2015) and Morrissey *et al.* (2015).

#### 2.2.1 Persistence of neonicotinoids in soil

Although neonicotinoids applied through a seed dressing are designed to be taken up into the target crop plant, only 1.6–20% of the active ingredient is absorbed, with the majority remaining in the soil. A small proportion is dispersed through dust created whilst drilling (see Section 2.1.2).

Neonicotinoids can bind to soil with the strength of the binding dependent on various factors. Neonicotinoids are water soluble (see section 2.2.2) and may leach from soils if water is present. Leaching is lower and sorption is higher in soils with a high content of organic material (Selim *et al.* 2010) . In a recent comparison of soil types, Mörtl *et al.* 2016, Figure 2) found that clothianidin and thiamethoxam leached readily from sandy soils. Clay soils showed higher retention of neonicotinoids but the greatest retention was seen for loam soils. Correspondingly, the highest residual neonicotinoid concentrations were found in loam soils.

**Figure 2.**
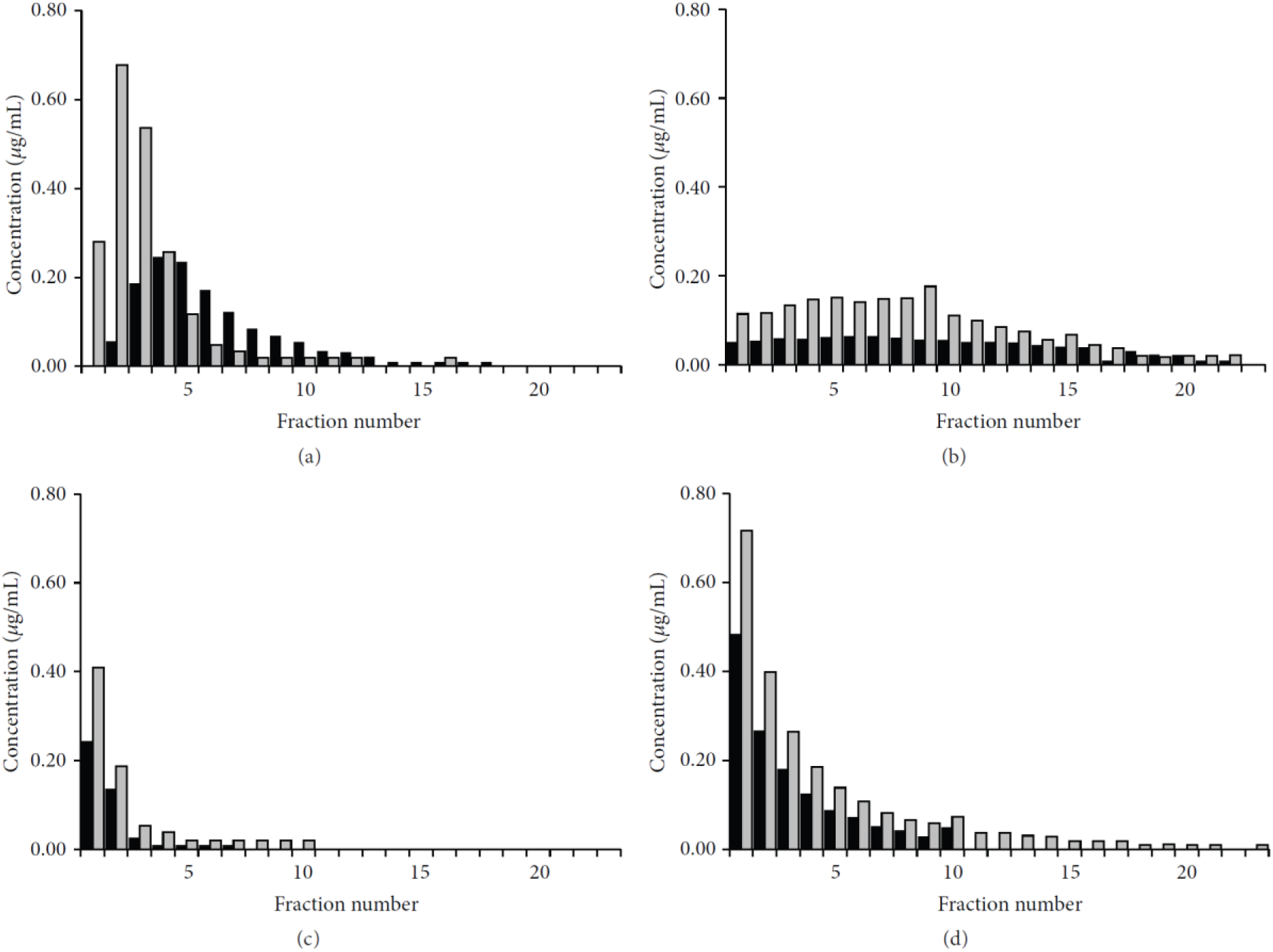
Elution profiles of clothianidin and thiamethoxam upon absorption on soils. Concentrations of clothianidin (black columns) and thiamethoxam (grey columns) measured in aqueous eluates from soil columns of (as) sand, (b) clay and (c) loam soils. Eluates from (d) pumice columns are shown as a control. Concentrations in 10 mL fractions of the eluate are shown in *μ*g/mL, as a function of the fraction number. Reproduced from Mörtl *et al.* (2016)

Whilst several studies have assessed dissipation half-life times (DT_50_) of neonicotinoids in soil, much of this work was conducted before the recent interest in the potentially deleterious effect of neonicotinoids on wider biodiversity. A review of available DT_50_ times from field and laboratory studies conducted between 1999 and 2013 were reviewed by Goulson (2013). Reported DT_50_s are highly variable and typically range from 200 to in excess of 1000 days for imidacloprid, 7-353 days for thiamethoxam and 148–6931 days for clothianidin. DT_50_s appear to be shorter for the nitro-substituted neonicotinoids, at 3–74 days for thiacloprid and 31–450 days for acetamiprid. DT_50_ values of over one year would suggest the likelihood of neonicotinoid bioaccumulation in the soil, assuming continuous input. However, these reported values are highly variable. At the time the EFSA reports were written only one field study was available that assessed neonicotinoid accumulation in the soil over multiple years with continued neonicotinoid input. Bonmatin *et al.* 2005 screened 74 samples of farmland soil from France for imidacloprid. Imidacloprid concentrations were higher in soils which had been treated in two consecutive years than those soils which had only received one treatment, suggesting the possibility of imidacloprid accumulation in the soil. However, as the study only looked at soils treated for a maximum of two years it is not clear whether residues would continue to increase. Two studies had been completed by 2013 but were not widely disseminated. These studies were carried out by Bayer and assessed levels of imidacloprid in soil over six years for seed-treated? barley in the UK (Placke 1998a) and spray application to orchard soils in Germany (Placke 1998b). Goulson (2013) reviewed this data and argued that the studies show accumulation of neonicotinoids in soils over time (Figure 3), with some indication that concentrations may begin to plateau after about five years. However, since the trials were terminated after six years it is not clear whether levels would have continued to increase.

**Figure 3.**
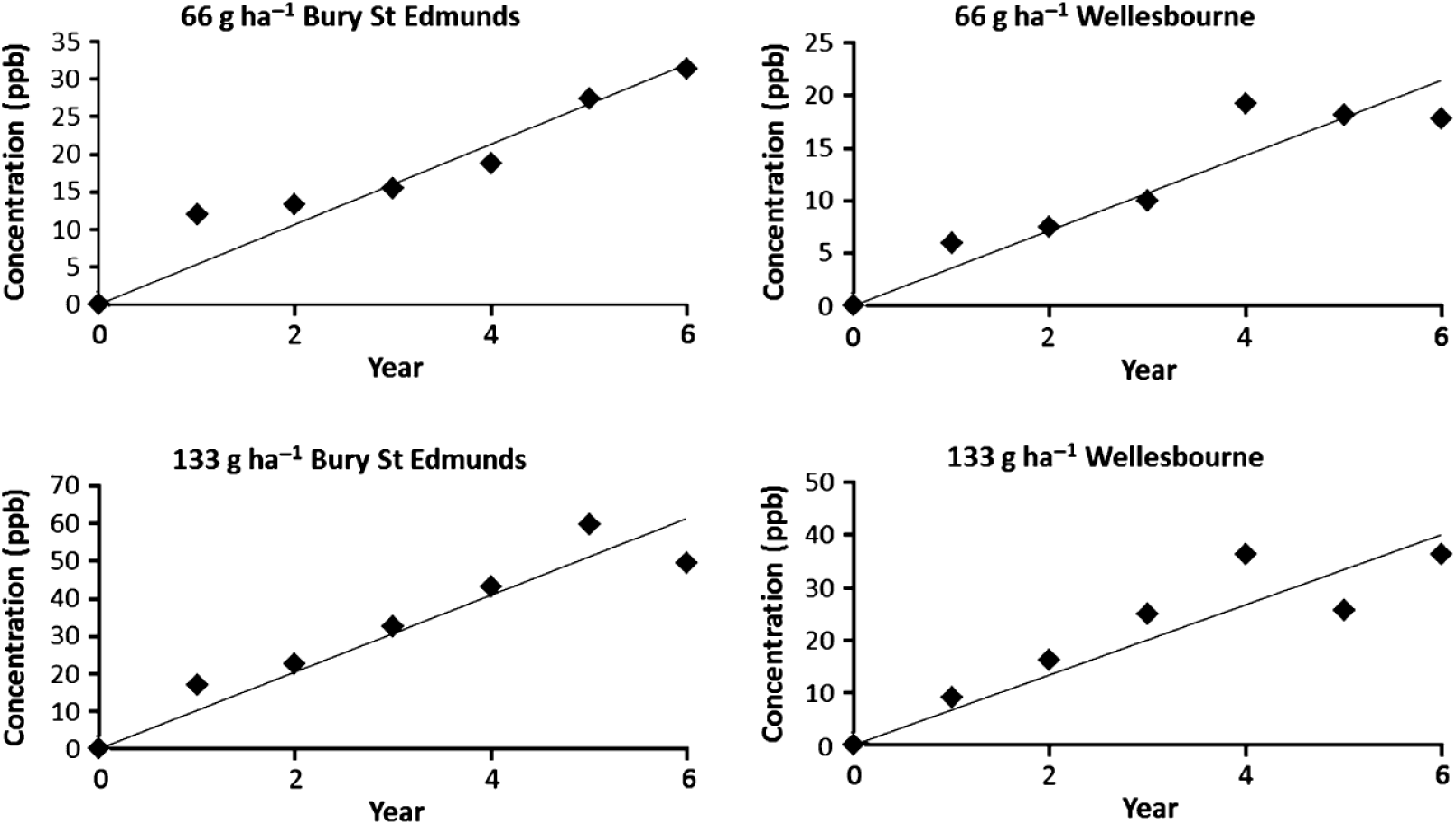
Levels of imidacloprid detected in soil into which treated winter wheat seeds were sown each autumn (1991–1996). Both study sites are in the east of England. Treatment rates were 66 and 133 g active ingredient ha^−1^ except in the first year, when it was 56 and 112 g, respectively. Data from Placke (1998a). Reproduced from Goulson (2013)

Since 2013 a number of studies have been published which have measured neonicotinoid levels in agricultural soils, have calculated DT_50_s of neonicotinoids in real world soils and have measured accumulation in the soil using extensive field trials and field sampling. Data on field-realistic neonicotinoid samples are summarised in Table 3. Jones *et al.* (2014) measured neonicotinoid concentrations in centre and edge soil samples from 18 fields across 6 English counties. Samples were collected in the spring of 2013, prior to crop planting. Imidacloprid (range <0.09–10.7 ng/g), clothianidin (range <0.02–13.6 ng/g) and thiamethoxam (range <0.02–1.5 ng/g) were detected. Residues from the centre of the fields were higher than for the edge of the fields (average imidacloprid 1.62 against 0.76 ng/g, average clothianidin 4.89 against 0.84 ng/g and average thiamethoxam 0.40 against 0.05 ng/g). Neonicotinoids not previously applied in the previous three years (predominantly imidacloprid) were detected in 14 of the 18 fields. Limay-Rios *et al.* (2015) analysed soil samples collected in the springs of 2013 and 2014 from 25 agricultural fields in Ontario, Canada before crops were sown and found average concentrations of 3.45 ng/g of clothianidin and 0.91 ng/g thiamethoxam, with total average neonicotinoid concentration of 4.36 ng/g, similar to the findings of Jones *et al.* (2014).

**Table 3.**
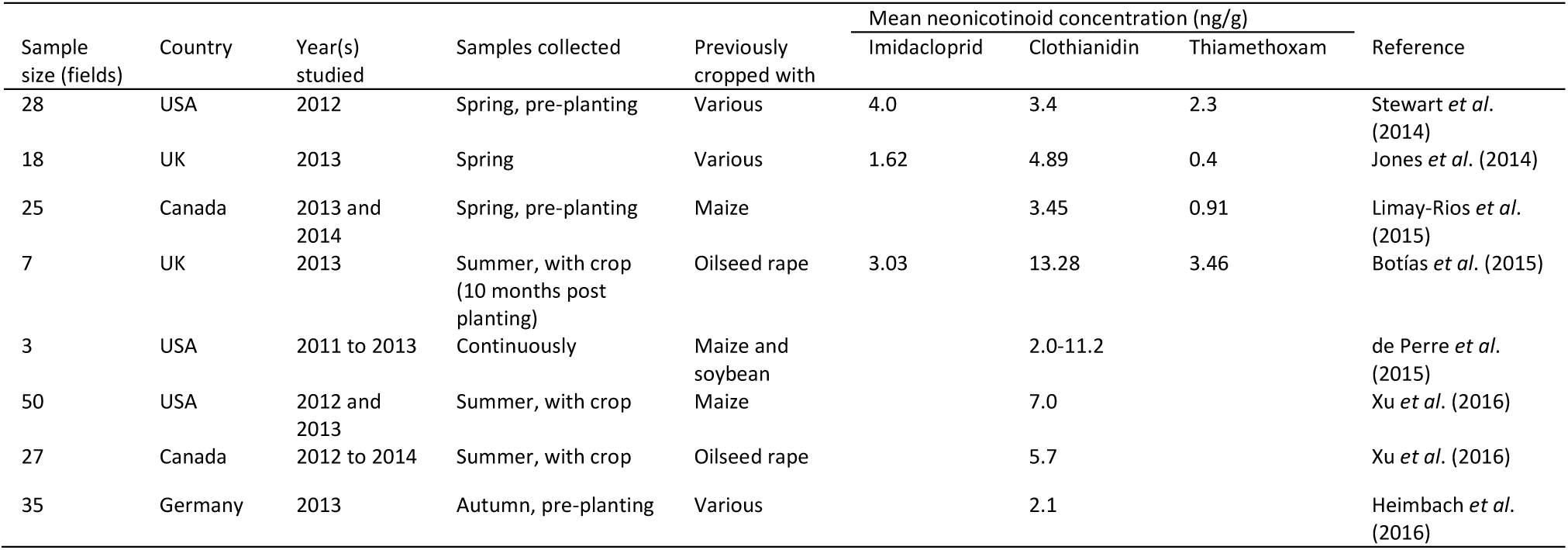
Summary of studies published since 2013 that document neonicotinoid concentrations in agricultural soils.

Botías *et al.* (2015) analysed soil samples from seven winter-sown oilseed rape and five winter-sown wheat fields collected in summer 2013, 10 months after the crops were sown. Samples were collected from field centres (oilseed rape only) and field margins (oilseed rape and winter wheat).

Imidacloprid (range ≤0.07–7.90 ng/g), clothianidin (range 0.41–28.6 ng/g), thiamethoxam (range ≤0.04–9.75 ng/g) and thiacloprid (range ≤0.01–0.22 ng/g) were detected. Residues from the centre of the oilseed rape fields were higher than for the edge of the oilseed rape fields (average imidacloprid 3.03 against 1.92 ng/g, average clothianidin 13.28 against 6.57 ng/g, average thiamethoxam 3.46 against 0.72 ng/g and average thiacloprid 0.04 against ≤0.01 ng/g). Whilst these values are higher than those measured by Jones *et al.* (2014) and Limay-Rios *et al.* (2015) they are within an order of magnitude at their greatest difference.

Hilton *et al.* (2015) presented previously private data from 18 industry trials conducted between 1995 and 1998 for thiamethoxam applied to bare soils, grass and a range of crops (potatoes, peas, spring barley, winter barley, soybean, winter wheat and maize). Thiamethoxam DT_50_s ranged between 7.1 and 92.3 days, with a geometric mean of 31.2 days (arithmetic mean 37.2 days). Across different application methods and environmental conditions, thiamethoxam declined to <10% of its initial concentration within one year. de Perre *et al.* (2015) measured soil clothianidin concentrations over 2011 to 2013, with clothianidin-treated maize sown in the springs of 2011 and 2013. Maize seeds were sown with seed dressings of 0.25 mg/seed and 0.50 mg/seed (Figure 4). At the lower concentration seed dressing, clothianidin residues in the soil ranged from approximately 2 ng/g before planting to 6 ng/g shortly after planting. At the higher seed dressing, clothianidin average residues ranged from 2 ng/g before planting to 11.2 ng/g shortly after planting. For the seed treatment of 0.5 mg/seed, de Perre *et al.* (2015) calculated a DT_50_ for clothianidin of 164 days. For the lower treatment of 0.25 mg/seed a DT_50_ of 955 days was calculated, though this model explained a much lower proportion of the data than the model for the 0.5 mg/seed data.

**Figure 4.**
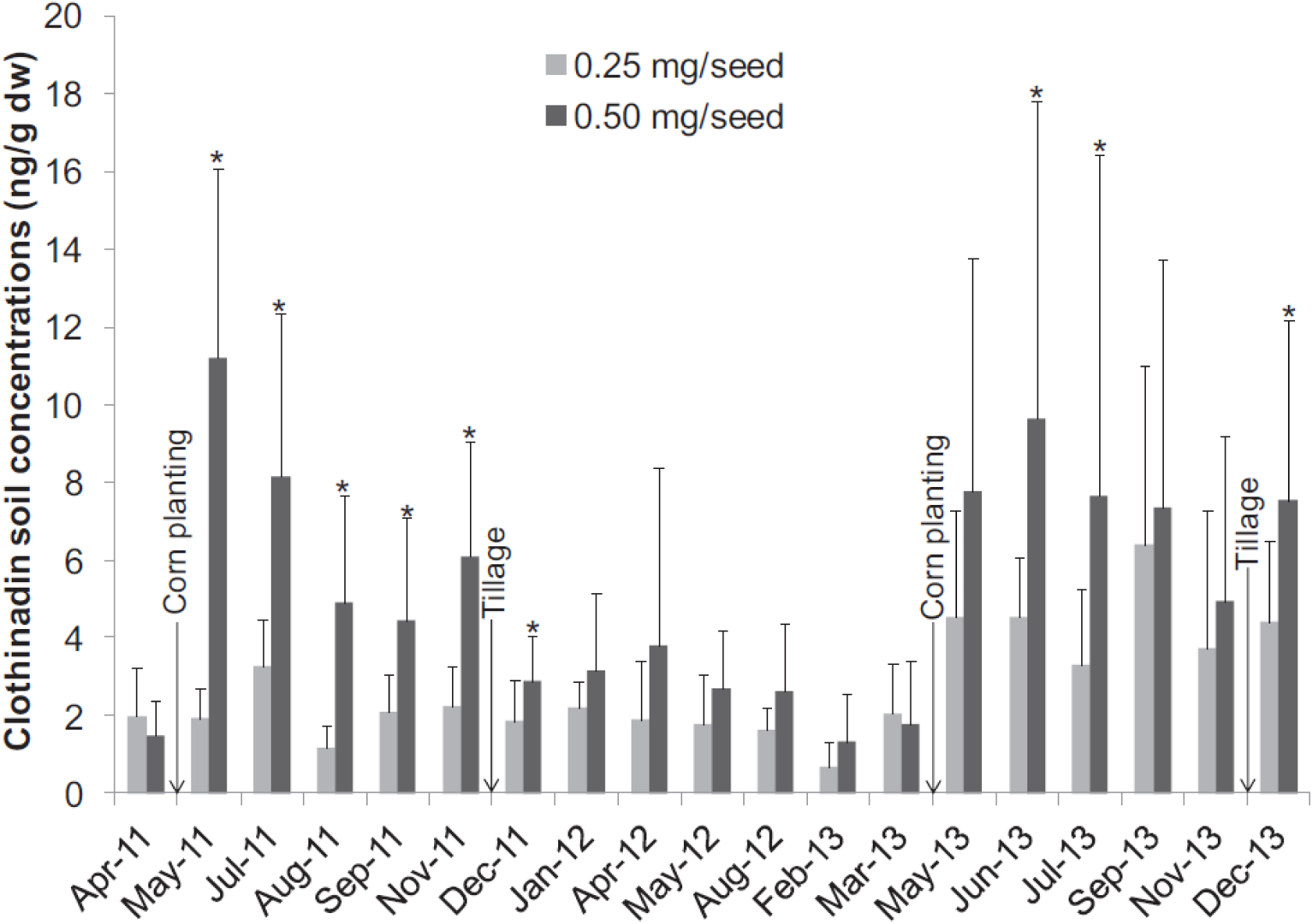
Mean clothianidin soil concentrations from 2011–2013 for each maize seed-coating rate (0.25 mg vs 0.50 mg of clothianidin/seed). Maize planting is presented because it represents the introduction of clothianidin in the field, and tillage events are also presented. Asterisks represent significantly different concentrations between seed-coating treatments for one sampling event (t test, *p* ≤0.05, n=13 and n=17 for 0.25 mg/seed and 0.50 mg/seed, respectively, from April 2011 to March 2013; n=15 for both seed treatment rates since May 2013). Reproduced from de Perre *et al.* (2015). Note – untreated soybeans were sown in 2012.

Schaafsma *et al.* (2016) calculated clothianidin DT_50_s in maize fields in Ontario, Canada in 2013 and 2014, including data published in Schaafsma *et al.* (2015). Soil samples were collected from 18 fields in the spring before crop planting. Average neonicotinoid concentrations (clothianidin and thiamethoxam aggregated) were 4.0 ng/g in 2013 and 5.6 ng/g in 2014. Using the observed residues and the recharge rate applied at planting via treated maize seeds, fields studied in 2013 had an estimated DT_50_ of 0.64 years (234 days) and fields studied in 2014 had an estimated DT_50_ of 0.57 years (208 days). For fields studied in both years the DT_50_ was calculated at 0.41 years (150 days). Schaafsma *et al.* conclude that, at current rates of neonicotinoid application in Canadian maize cultivation, soil residues of neonicotinoids will plateau at under 6 ng/g.

Using the same method, Schaafsma *et al.* also calculated imidacloprid DT_50_ using the data from Placke (1998a; 1998b; Table 4), producing a very similar DT_50_ of 0.57 years (208 days). Schaafsma *et al.* argue the Placke studies show neonicotinoid concentrations plateauing after repeated use of neonicotinoid seed treatments. However, observed levels were high, so even if plateauing occurred after six years the average concentration of neonicotinoids in the soil would be around 30 ng/g (Table 4).

**Table 4.** Observed concentrations of imidacloprid and estimated dissipation rates (half-life) in orchard soil in Germany and in winter barley fields in the United Kingdom. Data taken from Placke (1998a; 1998b). Half-life calculated iteratively by varying the half-life incrementally until the predicted and measured values are equal. Reproduced from Schaafsma *et al.* (2016)

| Field | Observed imidacloprid concentration (ng/g) | Half-life (years) |
| --- | --- | --- |
| Barley_66_1 | 31.4 | 0.74 |
| Barley_133_1 | 49.4 | 0.63 |
| Barley_66_2 | 17.8 | 0.53 |
| Barley_133_2 | 36.3 | 0.54 |
| Orchard_1 | 23.3 | 0.48 |
| Orchard_2 | 34.5 | 0.59 |
| Orchard_3 | 23.1 | 0.47 |
| Mean ± Standard Error | 30.8 | 0.57 ± 0.04 |

Xu *et al.* (2016) analysed soil samples from 50 maize producing sites in the Midwestern USA across 2012 and 2013 and soil samples from 27 oilseed rape producing sites in western Canada across 2012, 2013 and 2014. Samples were collected after planting, but it is not clear exactly how long after. Average clothianidin soil concentration at Midwestern maize producing sites with a range of 2–11 years of planting clothianidin-treated seeds was 7.0 ng/g with a 90^th^ percentile concentration of 13.5 ng/g. Xu *et al.* argue that this average is similar to the theoretical soil concentrations (6.3 ng/g) expected from a single application of 0.25 mg clothianidin-treated maize seed. Clothianidin levels in soil appear to plateau after 4 years (Figure 5a), but the sample size for sites with a history of more than four years is much smaller than the number of sites with a history of under four years of use. At the oilseed rape producing sites, average clothianidin concentrations were 5.7 ng/g with the 90^th^ percentile concentration of 10.2 ng/g. This is also similar to the theoretical soil concentration (6.7 ng/g) from a single application of oilseed rape seed treated at 4 g clothianidin per kg of seed (Figure 5b). The oilseed rape sites do not have the same history of clothianidin use but levels appear to be fairly stable over the four years of applications. For reference, 10 g clothianidin per kg of oilseed rape seed is the most common dosage rate in recent field trials (the Elado seed dressing, Section 3.1.2.1).

**Figure 5.**
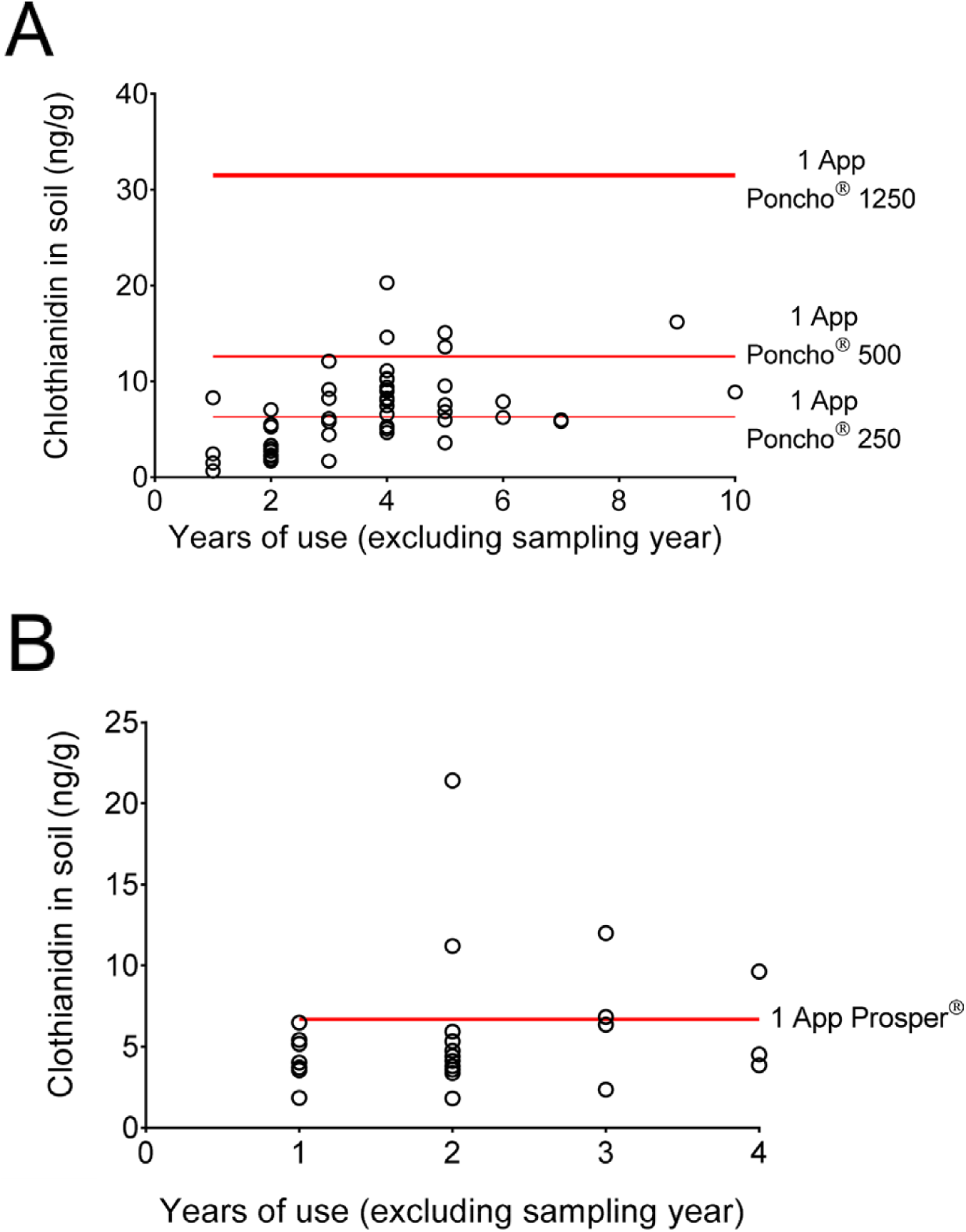
(a) Comparison of clothianidin concentrations in soil with years of clothianidin use for maize producing sites. Red lines indicate theoretical concentrations from a single application of clothianidin-treated seeds for three formulations. (b) Comparison of clothianidin concentrations in soil with years of clothianidin use for oilseed rape producing sites. Red lines indicate theoretical concentrations from a single application of clothianidin-treated seeds. Reproduced from Xu *et al.* (2016)

The current body of evidence shows that detectable levels of neonicotinoids are found in agricultural soils over a year after treated seeds were planted, clearly demonstrating a level of neonicotinoid persistence greater than the annual agricultural cycle. Moreover, neonicotinoids known not to have been recently used can still be present in soils several years after the last application date. The available data suggest that, whilst a proportion of the total neonicotinoids applied can and do persist in the soil from year to year, there appears to be sufficient degradation that means they do not continue to accumulate indefinitely (bioaccumulation) but instead plateau after 2–6 years of repeated application. However, these studies also show that overall, the annual sowing of neonicotinoid-treated seed results in chronic levels of neonicotinoid soil contamination in the range of 3.5–13.3 ng/g for clothianidin and 0.4–4.0 ng/g for thiamethoxam which will act as a constant source of exposure for soil dwelling organisms, and for neonicotinoid transport into the wider environment.

#### 2.2.2 Persistence of neonicotinoids in water and transport mechanisms for contamination of aquatic systems

Neonicotinoids are soluble in water, a property that is necessary for them to function effectively as systemic pesticides which can be taken up by crops. The solubility of neonicotinoids depends on local conditions such as ambient temperature, water pH and the form that the neonicotinoids are applied in, such as granules, as a seed dressing or as dust drift from seed drilling (Bonmatin *et al.* 2015). Under standard conditions (20°C, pH 7), neonicotinoid solubility varies between 184 (moderate) to 590,000 (high) mg/L for thiacloprid and nitenpyram respectively (PPDB 2012). The values for clothianidin, imidacloprid and thiamethoxam are 340 (moderate), 610 (high) and 4,100 (high) mg/L respectively. In contrast, Fipronil has a solubility 2–3 orders of magnitude lower at 3.78 mg/L under the same conditions.

Because of the high solubility of neonicotinoids in water, concerns were raised that neonicotinoids might be passing into water bodies in the wider environment and that this may pose a risk for aquatic organisms. Available evidence to 2015 was reviewed by Bonmatin *et al.* 2015 and Morrissey *et al.* 2015. In general, under simulated environmental conditions, neonicotinoids readily leach into water (Gupta *et al.* 2008; Tisler *et al.* 2009). Neonicotinoids have been identified passing into waterways through several different routes. These include direct leaching into ground water and subsequent discharge into surface water, decay of treated plant material in waterways and direct contact from dust from the drilling of treated seed, treated seeds or spray drift into water bodies (Krupke *et al.* 2012; Nuyttens *et al.* 2013). The majority of this contamination is thought to occur from run-off after acute rainfall (Hladik *et al.* 2014; Sánchez-Bayo and Hyne 2014; Main *et al.* 2016). Run-off will be particularly severe where soil organic content is low and on steep slopes (Goulson 2013).

Whilst rainfall during or shortly after the planting season appears to be the main mechanism for neonicotinoid transport into waterbodies, detectable levels of neonicotinoids can be found in prairie wetlands in Canada during early spring before the planting season (Main *et al.* 2014). Main *et al.* (2016) analysed snow, spring meltwater, particulate matter and wetland water from 16 wetland sites adjacent to agricultural fields that had been used to grow either oilseed rape (canola, treated with neonicotinoids) or oats (not treated). They found that all meltwater samples were contaminated with clothianidin and thiamethoxam in the range of 0.014–0.633 μg/L (1 μg/l = 1 ppb). Levels of contamination in meltwater were higher adjacent to fields planted with neonicotinoid-treated oilseed rape in the previous year (mean 0.267 μg/L). However, fields planted with non-neonicotinoid-treated oats in the previous year still showed similar levels of contamination (mean 0.181 μg/L). Treated oilseed rape and untreated oats are frequently rotated from year to year (Main *et al.* 2014), and the small difference in neonicotinoid concentration in meltwater from fields previously planted with treated and untreated crops suggests the persistence of neonicotinoids in the soil over multiple years (see Section 2.2.2). The findings of this study suggest that neonicotinoid active ingredients previously bound to soil particles are eroded during spring freeze-thaw cycles. The demonstration of this route of transport in addition to general rainfall suggests a more chronic transport of neonicotinoids into water bodies outside the main period of crop planting.

The effect of neonicotinoids on aquatic habitats will depend on their persistence therein. Field and laboratory studies investigating the breakdown of imidacloprid, thiamethoxam and clothianidin in water report half-lives of minutes to several weeks depending on the conditions, several of which are not field-realistic (see Anderson *et al.* 2015; Lu *et al.* 2015). There has been no formal review of the degradation of neonicotinoids in water and existing literature consists of published peer review studies and grey literature government studies, all using different methodologies. However, a number of studies have attempted to measure neonicotinoid degradation under field-realistic conditions. Peña *et al.* (2011) measured degradation of thiamethoxam in wastewaters and sewage in Spain finding maximum absorption at 250–255 nm, suggesting high susceptibility to direct photolysis from natural light. In control waters thiamethoxam half-life was found to be 18.7 hours (Peña *et al.* 2011). Under natural light in rice paddies in Japan, imidacloprid had a half-life of 24.2 hours (Thuyet *et al.* 2011). Under natural light in Switzerland von Gunten *et al.* (2012) reported a half-life of 2 hours for imidacloprid and 254 hours for acetamiprid. Under laboratory conditions, Lu *et al.* (2015) measured half-lives for five neonicotinoids under differing conditions to mimic the seasonal change found in Canada (Table 5). They found 7-8-fold variation in the rate of neonicotinoid photolysis due to the variation in light levels across the season. The results are broadly similar to previously published studies with nitro-substituted neonicotinoid half-lives in the region of <1–3 days depending on light levels.

**Table 5.**
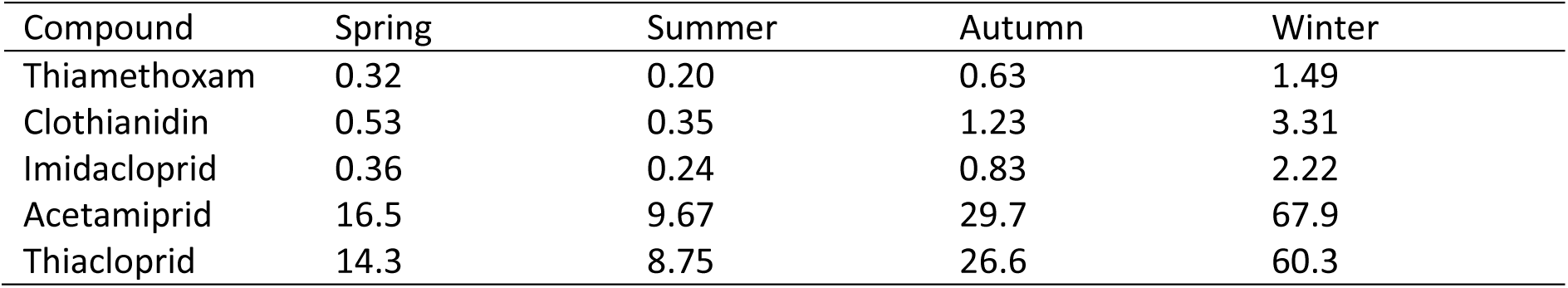
Estimated photolysis and half-lives (t_1/2E_) (days) for neonicotinoid pesticides in surface water at 50°N latitude for spring, summer, autumn and winter by sunlight on clear days. Reproduced from Lu *et al.* (2015)

In addition to these peer reviewed studies, Lu *et al.* drew comparison with European Commission regulatory studies on neonicotinoid compounds (EC 2004a; EC 2004b; EC 2005; EC 2006). The European Commission studies found half-lives in water of 3.3 hours for clothianidin, 2.3–3.1 days for thiamethoxam, 34 days for acetamiprid and 80 days for thiacloprid. The exact methodology used in these studies is unclear and inconsistent (see Lu *et al.* 2015 discussion). Nevertheless, the overall trend is consistent with the cyano-substituted neonicotinoids (acetamiprid and thiacloprid) taking 1–2 orders of magnitude longer to degrade than the nitro-substituted neonicotinoids (thiamethoxam, clothianidin and imidacloprid). The short half-lives of these three, most widely used neonicotinoids suggests that, under field conditions, free neonicotinoids in surface waters should be broken down by natural light in a matter of hours or days. However, local environmental conditions can affect this, with increasing turbidity increasing neonicotinoid persistence. Moreover, in mesocosm experiments, photolysis of thiamethoxam was found to be negligible at depths of greater than 8 cm (Lu *et al.* 2015). This significant light attenuation through the water column suggests that neonicotinoids may be shielded from photolysis even in shallow waterbodies. In waterbodies such as groundwater that are not exposed to light there will be no photolysis. In these circumstances clothianidin is persistent and has the potential to accumulate over time (Anderson *et al.* 2015), though empirical data demonstrating this is lacking.

#### 2.2.3 Levels of neonicotinoid contamination found in waterbodies

The most comprehensive review of levels of neonicotinoid contamination in global surface waters was conducted by Morrissey *et al.* (2015), though see also Anderson *et al.* (2015). Morrissey reviewed reported average and peak levels of neonicotinoid contamination from 29 studies from 9 countries between 1998 and 2013. The water bodies studied included streams, rivers, drainage, ditches, groundwater, wetlands, ponds, lakes, puddled surface waters and runoff waters. Study systems were adjacent to or receiving run-off water from agricultural land. From this dataset (Figure 6), the geometric mean for average surface water neonicotinoid concentration was 0.13 μg/L (=0.13 ppb, n=19 studies) and the geometric mean for peak surface water concentration was 0.63 μg/L (=0.63 ppb, n=27 studies). Because most monitoring schemes use spot sampling, they are likely to underreport the true maximum concentrations that occur immediately after maximum periods of neonicotinoid influx (Xing *et al.* 2013). As peak concentrations are often found after acute events such as heavy rainfall, this limits our understanding of the true average and maximum concentrations that are found in waterbodies.

**Figure 6.**
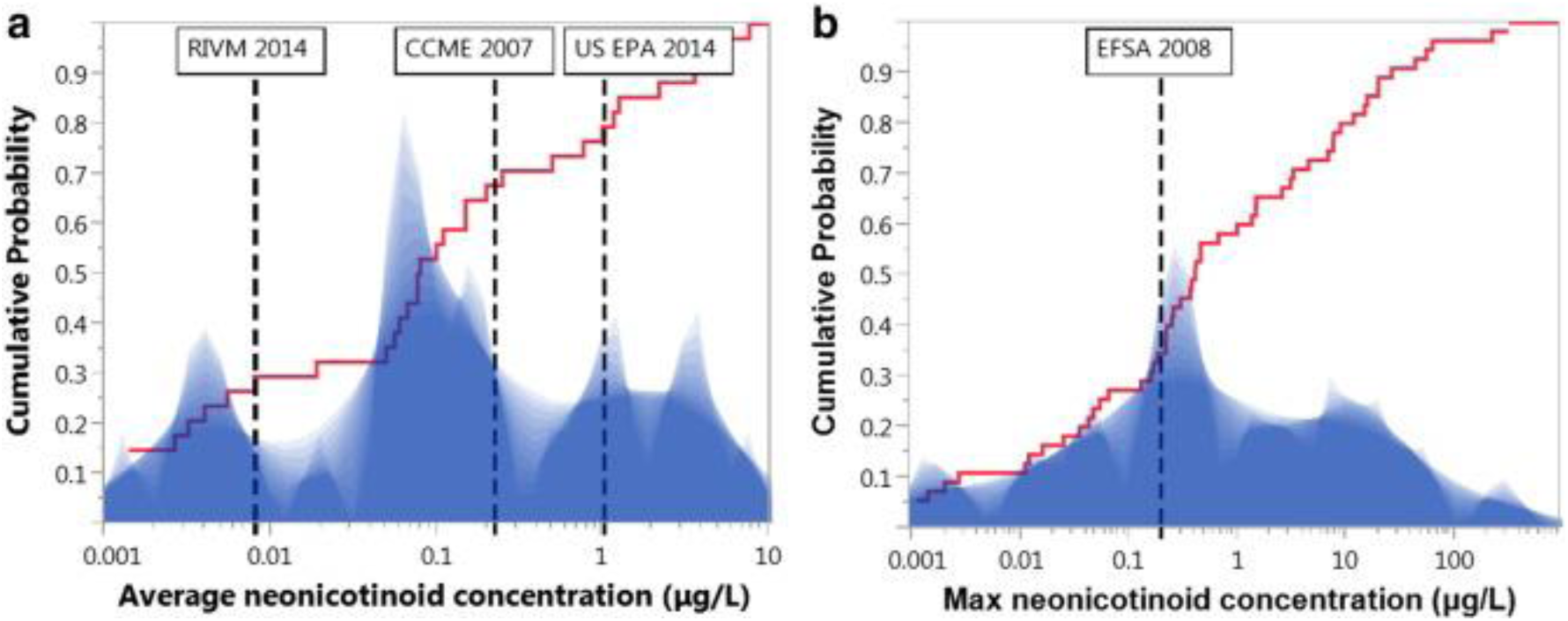
Shadow histogram of a) average and b) maximum individual neonicotinoid concentrations (log scale, μg/L) reported from water monitoring studies. Overlaid is the cumulative distribution probability (red ascending line) using all available surface water monitoring data showing proportion of data below any given neonicotinoid concentration. Vertical dashed lines illustrate multiple ecological quality reference values set for average imidacloprid water concentrations (RIVM 2014: 0.0083 μg/L, CCME 2007: 0.23 μg/L and US EPA 2014: 1.05 μg/L) or for maximum imidacloprid water concentrations (EFSA, 2008: 0.2 μg/L). Reproduced from Morrissey *et al.* 2015.

Since Morrissey *et al.* (2015) was published, a number of studies have become available documenting broadly similar neonicotinoid contamination levels in a wide range of aquatic environments. At a small scale in agricultural regions, Schaafsma *et al.* (2015) measured concentrations in surface water (puddles and ditches) in and around 18 maize fields in Ontario, Canada. They found arithmetic mean residues of 0.002 μg/L of clothianidin (maximum = 0.043 μg/L) and 0.001 μg/L of thiamethoxam (maximum = 0.017 μg/L). In Iowa, USA, Smalling *et al.* (2015) assessed six wetlands surrounded by agricultural land and found arithmetic mean neonicotinoid concentrations of 0.007 μg/L (maximum 0.070 μg/L). Away from agricultural land, Benton *et al.* (2016) measured concentrations in mountain streams in the southern Appalachians, USA, where eastern hemlock forests are treated with imidacloprid to control pests. Average concentrations of 0.067 μg/L of imidacloprid (maximum = 0.379 μg/L) were found in seven of the 10 streams investigated. de Perre *et al.* (2015) measured concentrations of clothianidin in groundwater below fields of treated maize. Data on average concentrations are not available but concentrations peaked at 0.060 μg/L shortly after crop planting.

At a wider scale, Qi *et al.* (2015) and Sadaria *et al.* (2016) measured concentrations in wastewater treatment plants. Qi *et al.* (2015) recorded imidacloprid at concentrations between 0.045–0.100 μg/L in influent and 0.045–0.106 μg/L in effluent at five waste water treatment plants in Beijing, China with no data available on arithmetic mean concentrations. Sadaria *et al.* (2016) assessed influent and effluent wastewater at 13 conventional waste water treatment plants around the USA. For influent, imidacloprid was found at arithmetic mean concentrations of 0.061 μg/L, acetamiprid at 0.003 μg/L and clothianidin at 0.149 μg/L. For effluent, imidacloprid was found at concentrations of 0.059 μg/L, acetamiprid at 0.002 μg/L and clothianidin at 0.070 μg/L.

Two nationwide surveys for neonicotinoids were also published. Hladik and Kolpin (2016) measured neonicotinoid concentrations in 38 streams from 24 US states plus Puerto Rico. Five neonicotinoids (acetamiprid, clothianidin, dinotefuran, imidacloprid, thiamethoxam) were recorded with at least one compound found in 53% of sampled streams, with an arithmetic mean contamination of 0.030 μg/L and median contamination of 0.031 μg/L. Thiacloprid was not recorded. Székács et al. (2015) conducted a nationwide survey of Hungarian watercourses, finding clothianidin at concentrations of 0.017–0.040 μg/L and thiamethoxam at concentrations of 0.004–0.030 μg/L.

Across all studies, the highest levels of neonicotinoid contamination were found in agricultural areas. In the most comprehensive nationwide survey of streams across the USA conducted between 2012 and 2014, levels of clothianidin and thiamethoxam contamination (the now dominant agricultural neonicotinoids) were significantly positively correlated with the proportion of the surrounding landscape used for crop cultivation (Hladik and Kolpin 2016). The most acute levels of neonicotinoid contamination in agricultural areas are reported from surface water in the immediate vicinity of cultivated crops. Puddles adjacent to fields planted with neonicotinoid-treated maize seeds were found to contain maximum concentrations of 55.7 μg/L clothianidin and 63.4 μg/L thiamethoxam in Quebec, Canada (Samson-Robert *et al.* 2014). Surface water in the Netherlands had imidacloprid concentrations up to 320 μg/L (van Dijk *et al.* 2013) and transient wetlands found in intensively farmed areas of Texas had thiamethoxam and acetamiprid concentrations of up to 225 μg/L (Anderson *et al.* 2013). In Hungary, the highest neonicotinoid concentrations of 10–41 μg/L were found in temporary shallow waterbodies after rain events in early summer (Székács et al. 2015). More generally, watercourses draining agricultural fields had high levels of neonicotinoids after rainfall in Canada, the USA and Australia (Hladik *et al.* 2014, Sánchez-Bayo and Hyne 2014). Where repeated sampling of the same site has been carried out, the highest neonicotinoid concentrations have been found in early summer and are associated with rainfall during the planting season (Main *et al.* 2014; Hladik *et al.* 2014). Hladik and Kolpin (2016) measured neonicotinoid concentrations in three agriculturally affected streams in Maryland and Pennsylvania and found peak levels after rain events during the crop planting season in May, though this could not be formally statistically analysed due to low sample size (Figure 7).

**Figure 7.**
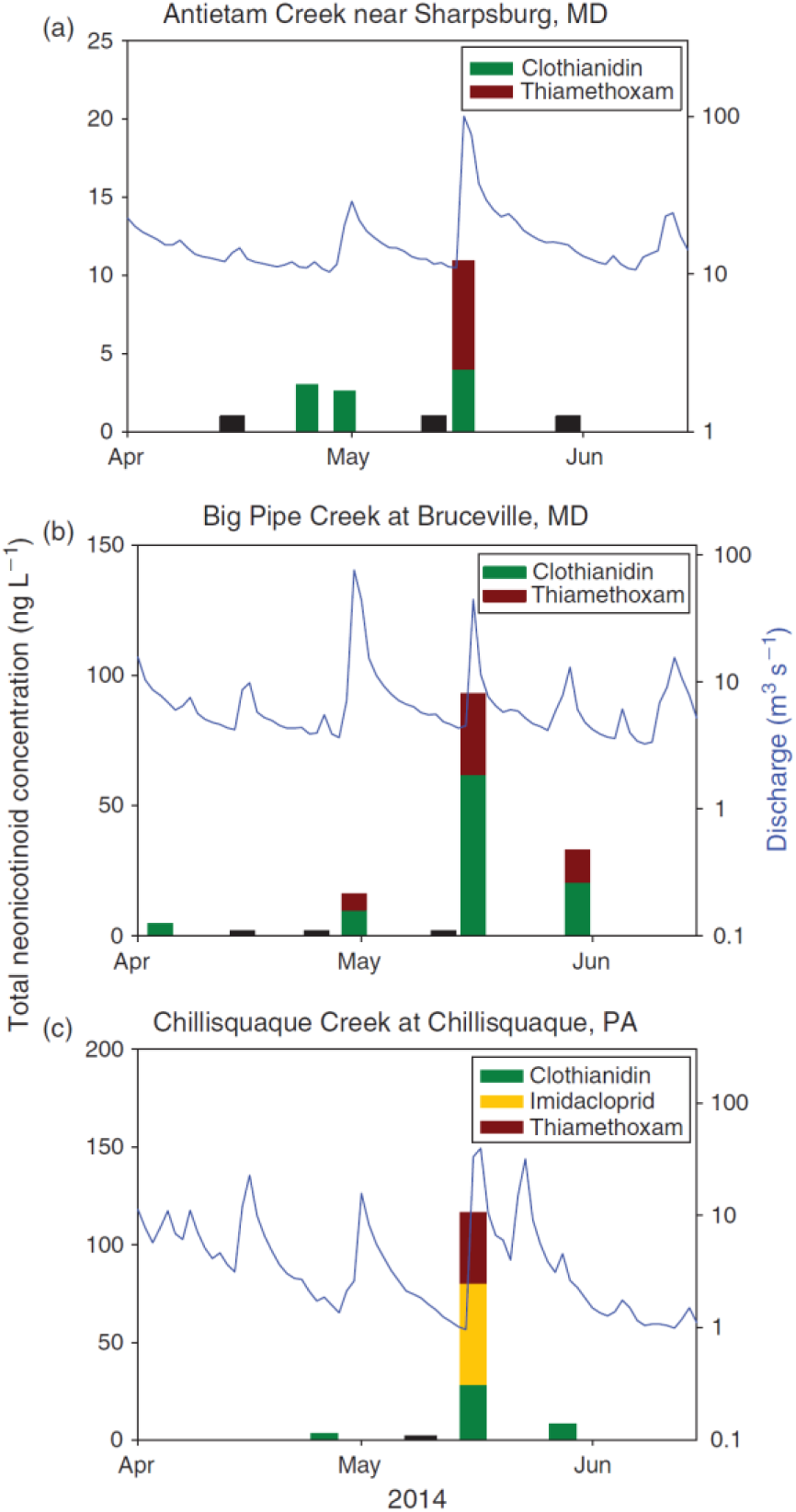
Concentrations of clothianidin, imidacloprid and thiamethoxam and the corresponding stream discharge at three sites in the Chesapeake Bay area sampled in 2014. Black bars represent samples where no neonicotinoids were detected. Reproduced from Hladik and Kolpin (2016)

In addition to agricultural run-off, urban areas also contribute towards neonicotinoid contamination of waterbodies. Whilst the use of imidacloprid as an agricultural pesticide has declined it is still found in a wide range of domestic products and veterinary treatments for pets (Goulson *et al.* 2013). Hladik and Kolpin (2016) continuously monitored neonicotinoid levels in Slope Creek, a stream surrounded by a largely urban catchment (39% urban) and the Chattahoochee river which includes the drainage of Slope Creek and overall has a lower proportion of urbanisation (9%). Imidacloprid was the dominant neonicotinoid found, present in 87% of the 67 collected samples (Figure 8). Dinotefuran and acetamiprid were less frequently encountered. Unlike in the studied watercourses draining agricultural land, no significant relationship was seen with stream flow in either Slope Creek or the Chattahoochee river. Hladik and Kolpin suggest that this may be because, unlike for the planting period of arable crops, there is no distinct period of use for domestic imidacloprid in an urbanised catchment. No clothianidin or thiamethoxam were detected, probably because neither catchment contained cultivated crops.

**Figure 8.**
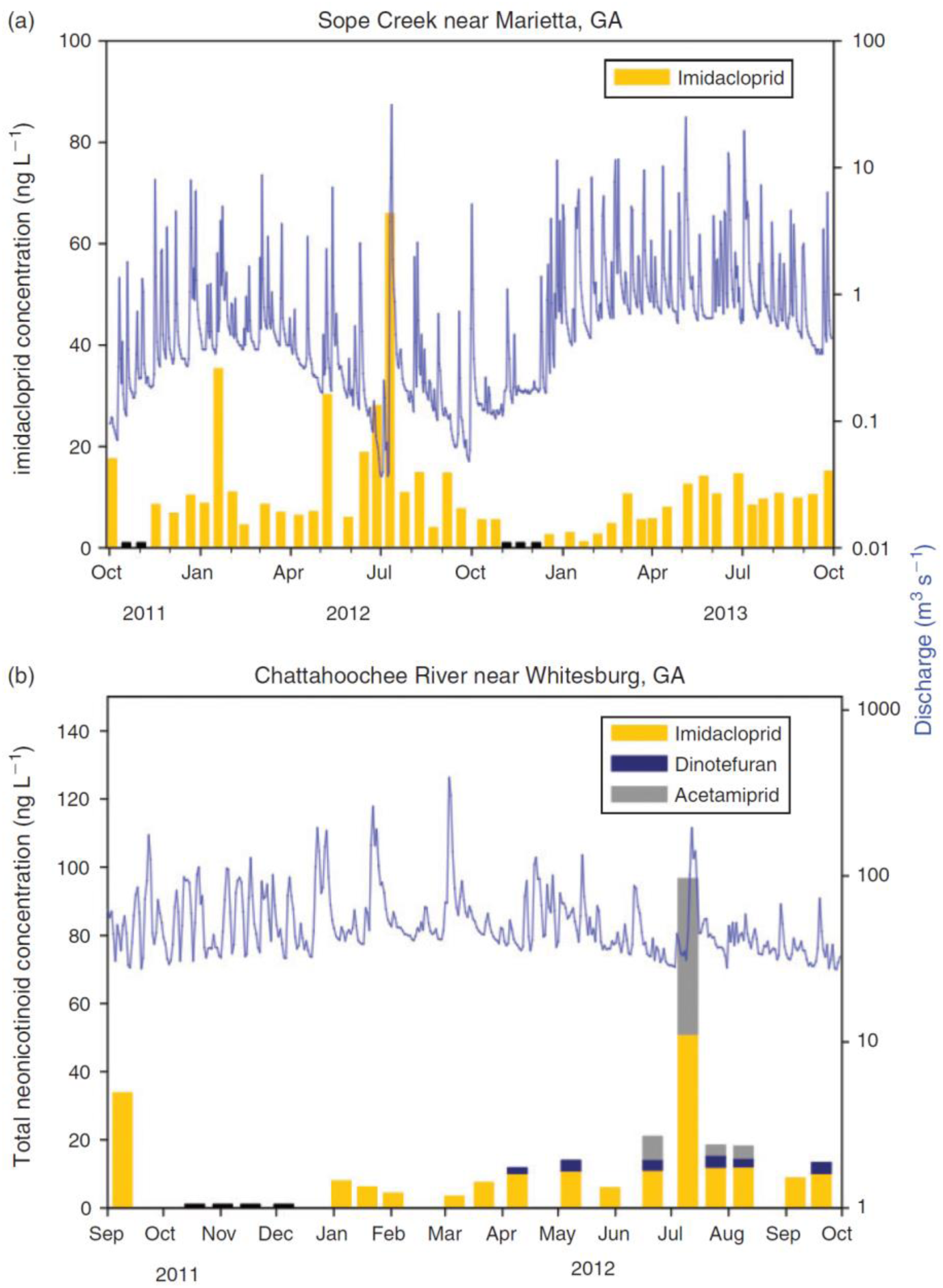
(a) Concentrations of imidacloprid and the corresponding stream discharge from October 2011 to October 2013 for Sope Creek (a largely urban catchment) and (b) Concentrations of imidacloprid, dinotefuran and acetamiprid along with the corresponding stream discharge from September 2011 to September 2012 for Chattahoochee River. Black bars represent samples where no neonicotinoids were detected. Reproduced from Hladik and Kolpin (2016)

#### 2.2.4 Risk of exposure from and uptake of neonicotinoids in non-crop plants

Since neonicotinoids are water soluble and can persist in soils and waterbodies there is the possibility that they may be taken up by any wild plants present nearby. In April 2013 little empirical data was available documenting neonicotinoid contamination of wild plants. The EFSA reports considered that uptake of neonicotinoids by wild weed plants and subsequent exposure would be negligible, as weeds will not be present in the field when the crop is sown and considerable uptake via the roots would be unlikely as the substance is concentrated around the treated seed. No comment was made on the potential uptake of neonicotinoids by other wild plants in the agricultural environments. In the single study available in 2013, Krupke *et al.* (2012) found that dandelions *Taraxacum agg*. growing near to fields planted with neonicotinoid-treated maize contained between 1.1 to 9.4 ng/g clothianidin and <1.0 (LOD) to 2.9 ng/g thiamethoxam. They did not assess whether the pesticides were found in the pollen or nectar. It was not clear whether the contamination came from neonicotinoid dust settling on the external surface of the plants or if the neonicotinoids had been directly taken up through the roots, in which case we would expect them to be present inside all plant tissues, pollen and nectar. Since April 2013, a number of studies have been published which demonstrate that neonicotinoids are frequently taken up in wild plants surrounding agricultural fields (Table 6).

**Table 6.** Summary of studies published since 2013 that document mean neonicotinoid residues in wild plant tissues, pollen and nectar in plants growing close to neonicotinoid-treated agricultural crops. The results of Krupke *et al.* (2012) are included for reference

| Sample size | Vegetation adjacent to | Samples collected | Sample type | Mean neonicotinoid concentration (ng/g) |  |  |  | Reference |
| --- | --- | --- | --- | --- | --- | --- | --- | --- |
|  |  |  |  | Thiamethoxam | Clothianidin | Imidacloprid | Thiacloprid |  |
| 43 | Oilseed rape | May-June 2013 | Pollen | 14.81 |  | 0.56 | <0.04 | Botías *et al.* (2015) |
| 55 | Wheat | May-June 2013 | Pollen | 0.14 |  | <0.16 | <0.04 | Botías *et al.* (2015) |
| 24 | Oilseed rape | May-June 2013 | Nectar | 0.10 |  |  |  | Botías *et al.* (2015) |
| 8 | Wheat | May-June 2013 | Nectar | <0.10 |  |  |  | Botías *et al.* (2015) |
| 33 | Maize | Summer 2014 and 2015 | Nectar * |  | 0.2-1.5 |  |  | Mogren and Lundgren (2016) |
| 40 | Maize | June 2014 | Foliage |  | 0.4 |  |  | Pecenka and Lundgren (2015) |
| 50 | Maize | July 2014 (1 month after planting) | Foliage |  | 0.69 |  |  | Pecenka and Lundgren (2015) |
| 100 | Oilseed rape | May-June 2013 | Foliage | 8.71 | 0.51 | 1.19 |  | Botías *et al.* (2016) |
| 375 | Maize | Summer 2014 and 2015 | Foliage |  | 0.5-13.5** |  |  | Mogren and Lundgren (2016) |
| 6 | Maize | Summer 2011 | Complete flower | 1.15 | 3.75 |  |  | Krupke *et al.* (2012) |
| 78 | Various | Summer 2012 | Complete flower | 7.2 | 1.4 | 1.1 |  | Stewart *et al.* (2014) |
| 7 | Oilseed rape | April-May 2013 (2 days after sowing) | Complete flowers and foliage |  | 1.2 |  |  | Rundlöf *et al.* (2015) |
| 8 | Oilseed rape | April-June 2013 (2 weeks after sowing) | Complete flowers and foliage |  | 1.0 |  |  | Rundlöf *et al.* (2015) |

\* Mogren and Lundgren (2016) sampled honeybees foraging on wild plants and directly extracted nectar from their crop. See main body of text for further discussion
\*\* Range of concentrations, data on mean concentrations not available

Botías *et al.* (2015) collected pollen and nectar from wildflowers growing in field margins adjacent to agricultural fields planted with neonicotinoid-treated oilseed rape and wheat. Pollen samples from 54 wild flower species were collected. Thiamethoxam, imidacloprid and thiacloprid were all detected. Thiamethoxam was the most frequently encountered neonicotinoid and levels were highly variable with the highest concentrations found in *Heracleum sphondylium* at 86 ng/g and *Papaver rhoeas* at 64 ng/g. There was substantial variation in the levels of contamination in the same wildflower species found in different field margins. Average levels of total neonicotinoid contamination in wildflower pollen were significantly higher in margins adjacent to treated oilseed rape (c. 15 ng/g) than for margins adjacent to treated wheat (c. 0.3 ng/g). Levels of neonicotinoids were much lower in wild plant nectar. Only thiamethoxam was detected at average levels of 0.1 ng/g in wild flowers adjacent to oilseed rape fields and <0.1 ng/g adjacent to wheat fields.

Botías *et al.* (2015) is the only available study which has specifically measured neonicotinoid concentrations in pollen and nectar directly taken from wild plants growing in close proximity to neonicotinoid-treated crops. Mogren and Lundgren (2016) assessed neonicotinoid concentrations in the nectar of five wild flower species sown as part of pollinator conservation measures which were located adjacent to neonicotinoid-treated maize. This was achieved by collecting honeybees seen to visit these flowers for nectar and extracting the contents of their crop for neonicotinoid residue analysis. Honeybees generally have a very high fidelity to visiting the same flower species on a single forage flight so the authors assumed that the nectar was representative of that particular species. Average clothianidin concentrations found in this nectar ranged between 0.2 and 1.5 ng/g, with significant differences found between wild plant species. Mogren and Lundgren (2016) also tested the foliage of seven wildflower species for neonicotinoid residues directly. There was high variability in clothianidin uptake between and within plant species (Figure 9). Sunflowers *Helianthus annuus* accumulated the highest levels with concentrations of 0–81 ng/g, with buckwheat *Fagopyrum esculentum* and phacelia *Phacelia tanacetifolia* accumulating lower levels at 0–52 ng/g and 0–33 ng/g respectively. Similarly high levels of variation were found by Botías *et al.* (2016) who sampled the foliage of 45 species of wild plant in field margins adjacent to treated oilseed rape crops. Average total neonicotinoid contamination was 10 ng/g, with the highest levels seen in creeping thistle *Cirsium arvense* of 106 ng/g of thiamethoxam. Pecenka and Lundgren (2015) looked specifically at clothianidin concentrations in milkweed *Asclepias syriaca* in field margins adjacent to clothianidin-treated maize. Levels were lower than the previous two studies, with mean levels of 0.58 ng/g with a maximum concentration of 4.02 ng/g.

**Figure 9.**
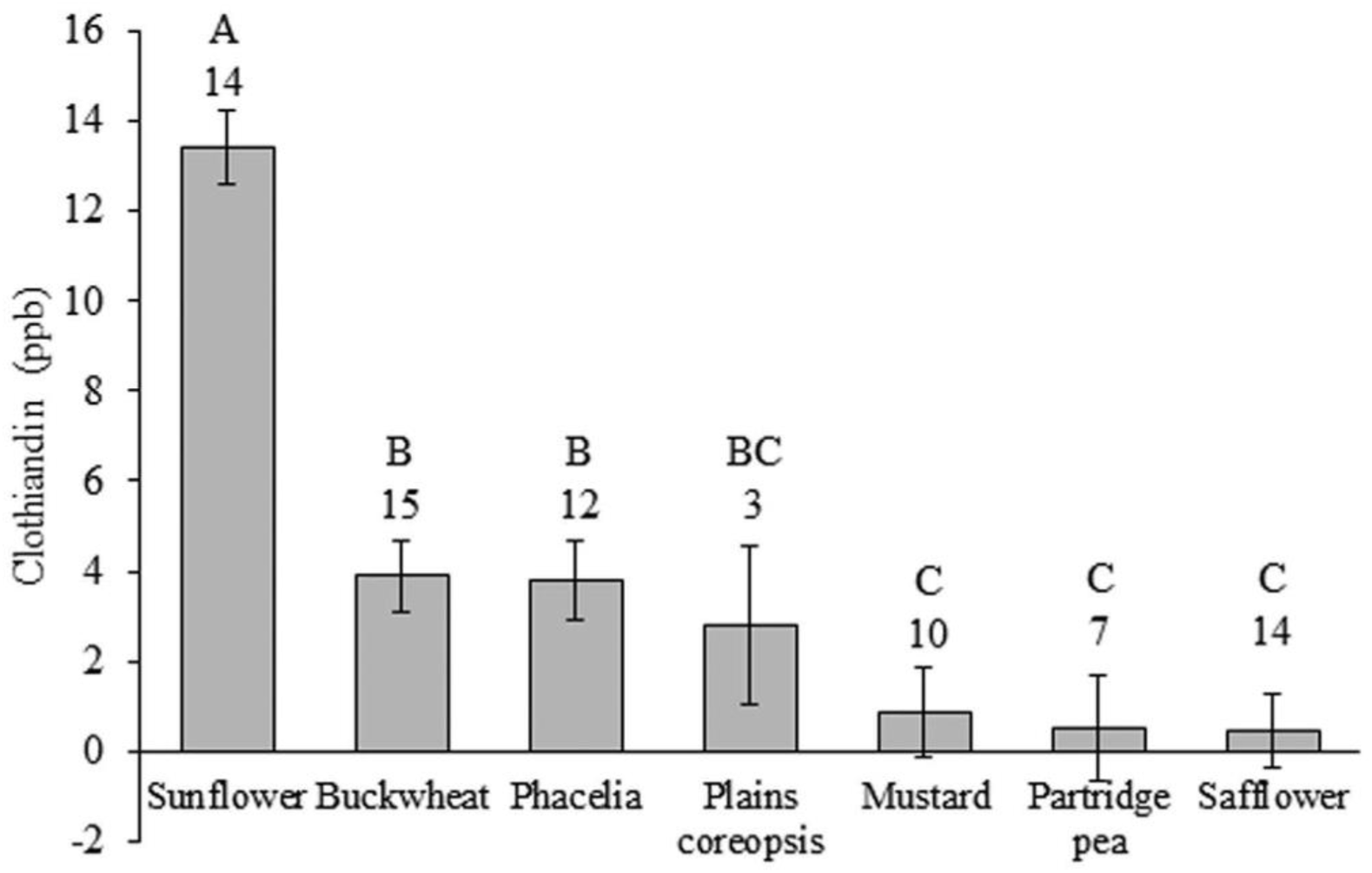
Concentrations of clothianidin in leaf tissues (mean±SE). Letters above bars show significant differences between plant species and numbers represent the number of site-years in which a particular species was analysed. Reproduced from Mogren and Lundgren (2016).

Whilst not looking at specific concentrations in pollen, nectar or foliage, Stewart *et al.* (2014) and Rundlöf *et al.* (2015) found total mean neonicotinoid concentrations of 10 ng/g and 1ng/g respectively in whole wild flower samples collected around neonicotinoid-treated fields. As discussed in Section 2.1.3, these levels may have been a direct result of neonicotinoid-contaminated dust drift onto surrounding vegetation and do not in and of themselves demonstrate uptake of neonicotinoids from contaminated soil and/or water.

Across all studies published since 2013, average levels of neonicotinoids in wild plants range from 1.0–7.2 ng/g in whole flower samples, 0.4–13.5 ng/g in foliage samples, <0.1–1.5 ng/g in nectar samples and <0.04 to 14.8 ng/g in pollen samples. Due to the limited number of studies available, it is difficult to make a comparison with levels in directly treated crop plants. However, they are broadly comparable to the levels found in the treated crop itself (see Section 2.1.1)

In 2013 it was known that honeybees collected neonicotinoid contaminated pollen from crop plants, but the extent to which this was diluted by uncontaminated pollen from wild plants was unknown. Krupke *et al.* (2012) found levels of clothianidin and thiamethoxam in honeybee-collected pollen that ranged between 0 and 88 ng/g, with the proportion of pollen collected from maize (the main treated crop in their study area) also varying substantially between 2.6 and 82.7%. There was no correlation between the proportion of maize pollen collected and the total neonicotinoid concentration. Given the uncertainty over the contamination of wild plants it was not clear what long term chronic neonicotinoid exposure was from pollen or nectar over a whole season. A number of studies have attempted to quantify the levels of neonicotinoids in bee-collected pollen and, through microscopic identification of the constituent pollen grains, to identify the major source of neonicotinoid contamination throughout the season. Most of these studies have used honeybee-collected pollen as the model, as pollen traps are easy to fit to apiaries that can be moved into targeted locations.

Studies are summarised in Table 7. Most of these studies used honeybees, placing apiaries out next to neonicotinoid-treated and untreated crops. As summarised in Section 2.1.1, bees placed near to treated crops collected pollen with higher concentrations of neonicotinoids (Cutler *et al.* 2014; Rundlöf *et al.* 2015; Long and Krupke 2016; Rolke *et al.* 2016). The highest levels of acute contamination are found when a large proportion of crop pollen is collected. Pohorecka *et al.* (2013) found average clothianidin concentrations of 27.0 ng/g in pollen samples (73.7% wildflower pollen) collected from apiaries adjacent to treated maize fields. Rundlöf *et al.* (2015) found average clothianidin concentrations of 13.9 ng/g in pollen samples (37.9% wildflower pollen) collected from apiaries adjacent to treated oilseed rape fields. Apiaries adjacent to untreated oilseed rape fields collected pollen consisting of 47.4% wildflower pollen with no detectable levels of neonicotinoids (<0.5 ng/g).

**Table 7.**
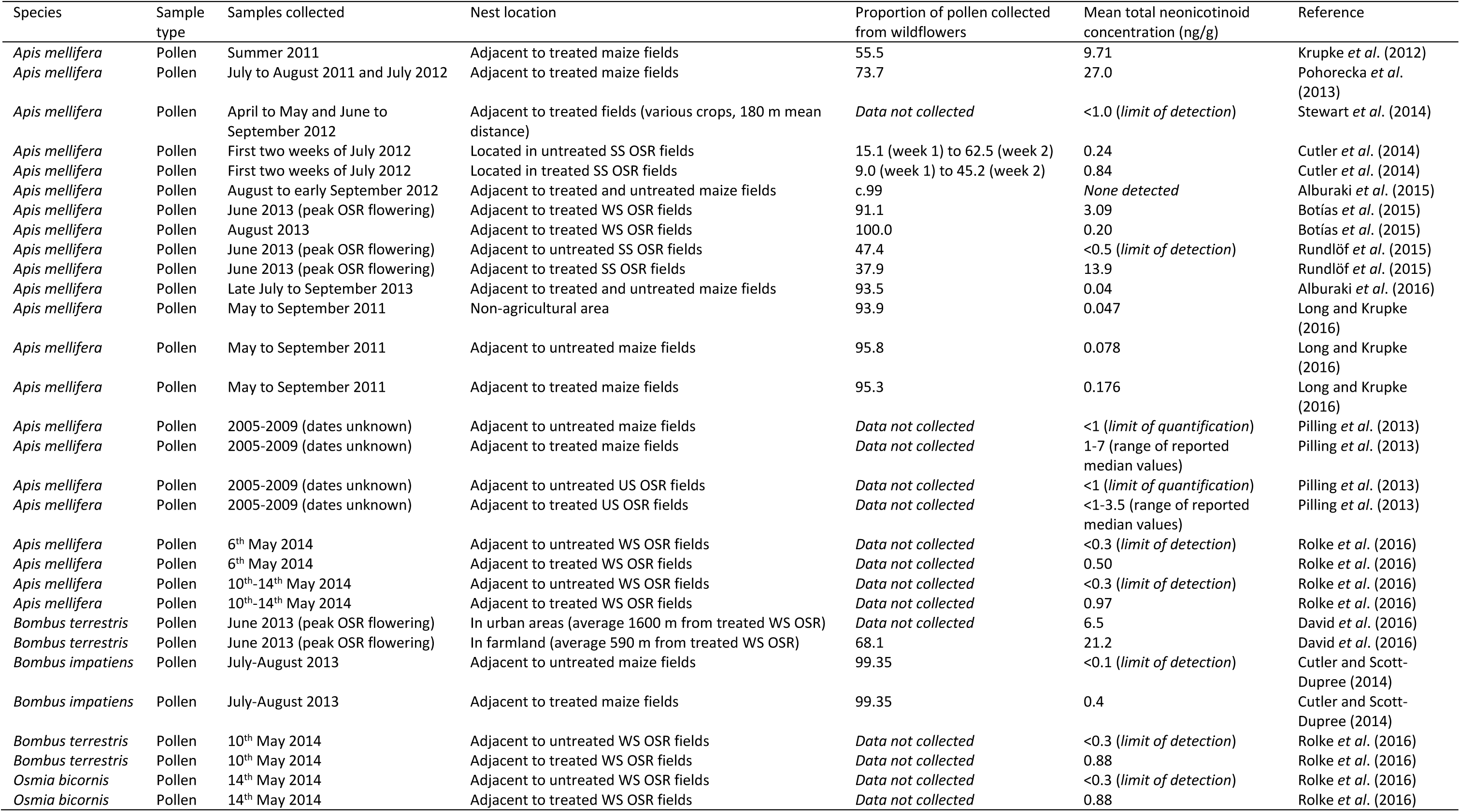
Summary of studies published since 2013 that document mean neonicotinoid residues in pollen collected by free-flying bees. The results of Krupke *et al.* (2012) and studies described in Section 2.1.1 are included for reference. SS = spring-sown, WS = winter-sown, US = unclear sowing date

| Species | Sample type | Samples collected | Nest location | Proportion of pollen collected from wildflowers | Mean total neonicotinoid concentration (ng/g) | Reference |
| --- | --- | --- | --- | --- | --- | --- |
| *Apis mellifera* | Pollen | Summer 2011 | Adjacent to treated maize fields | 55.5 | 9.71 | Krupke *et al.* (2012) |
| *Apis mellifera* | Pollen | July to August 2011 and July 2012 | Adjacent to treated maize fields | 73.7 | 27.0 | Pohorecka *et al.* (2013) |
| *Apis mellifera* | Pollen | April to May and June to September 2012 | Adjacent to treated fields (various crops, 180 m mean distance) | *Data not collected* | <1.0 (*limit of detection*) | Stewart *et al.* (2014) |
| *Apis mellifera* | Pollen | First two weeks of July 2012 | Located in untreated SS OSR fields | 15.1 (week 1) to 62.5 (week 2) | 0.24 | Cutler *et al.* (2014) |
| *Apis mellifera* | Pollen | First two weeks of July 2012 | Located in treated SS OSR fields | 9.0 (week 1) to 45.2 (week 2) | 0.84 | Cutler *et al.* (2014) |
| *Apis mellifera* | Pollen | August to early September 2012 | Adjacent to treated and untreated maize fields | c.99 | *None detected* | Alburaki *et al.* (2015) |
| *Apis mellifera* | Pollen | June 2013 (peak OSR flowering) | Adjacent to treated WS OSR fields | 91.1 | 3.09 | Botías *et al.* (2015) |
| *Apis mellifera* | Pollen | August 2013 | Adjacent to treated WS OSR fields | 100.0 | 0.20 | Botías *et al.* (2015) |
| *Apis mellifera* | Pollen | June 2013 (peak OSR flowering) | Adjacent to untreated SS OSR fields | 47.4 | <0.5 (*limit of detection*) | Rundlöf *et al.* (2015) |
| *Apis mellifera* | Pollen | June 2013 (peak OSR flowering) | Adjacent to treated SS OSR fields | 37.9 | 13.9 | Rundlöf *et al.* (2015) |
| *Apis mellifera* | Pollen | Late July to September 2013 | Adjacent to treated and untreated maize fields | 93.5 | 0.04 | Alburaki *et al.* (2016) |
| *Apis mellifera* | Pollen | May to September 2011 | Non-agricultural area | 93.9 | 0.047 | Long and Krupke (2016) |
| *Apis mellifera* | Pollen | May to September 2011 | Adjacent to untreated maize fields | 95.8 | 0.078 | Long and Krupke (2016) |
| *Apis mellifera* | Pollen | May to September 2011 | Adjacent to treated maize fields | 95.3 | 0.176 | Long and Krupke (2016) |
| *Apis mellifera* | Pollen | 2005-2009 (dates unknown) | Adjacent to untreated maize fields | *Data not collected* | <1 (*limit of quantification*) | Pilling *et al.* (2013) |
| *Apis mellifera* | Pollen | 2005-2009 (dates unknown) | Adjacent to treated maize fields | *Data not collected* | 1-7 (range of reported median values) | Pilling *et al.* (2013) |
| *Apis mellifera* | Pollen | 2005-2009 (dates unknown) | Adjacent to untreated US OSR fields | *Data not collected* | <1 (*limit of quantification*) | Pilling *et al.* (2013) |
| *Apis mellifera* | Pollen | 2005-2009 (dates unknown) | Adjacent to treated US OSR fields | *Data not collected* | <1-3.5 (range of reported median values) | Pilling *et al.* (2013) |
| *Apis mellifera* | Pollen | 6<sup>th</sup> May 2014 | Adjacent to untreated WS OSR fields | *Data not collected* | <0.3 (*limit of detection*) | Rolke *et al.* (2016) |
| *Apis mellifera* | Pollen | 6<sup>th</sup> May 2014 | Adjacent to treated WS OSR fields | *Data not collected* | 0.50 | Rolke *et al.* (2016) |
| *Apis mellifera* | Pollen | 10<sup>th</sup>-14<sup>th</sup> May 2014 | Adjacent to untreated WS OSR fields | *Data not collected* | <0.3 (*limit of detection*) | Rolke *et al.* (2016) |
| *Apis mellifera* | Pollen | 10<sup>th</sup>-14<sup>th</sup> May 2014 | Adjacent to treated WS OSR fields | *Data not collected* | 0.97 | Rolke *et al.* (2016) |
| *Bombus terrestris* | Pollen | June 2013 (peak OSR flowering) | In urban areas (average 1600 m from treated WS OSR) | *Data not collected* | 6.5 | David *et al.* (2016) |
| *Bombus terrestris* | Pollen | June 2013 (peak OSR flowering) | In farmland (average 590 m from treated WS OSR) | 68.1 | 21.2 | David *et al.* (2016) |
| *Bombus impatiens* | Pollen | July-August 2013 | Adjacent to untreated maize fields | 99.35 | <0.1 (*limit of detection*) | Cutler and Scott-Dupree (2014) |
| *Bombus impatiens* | Pollen | July-August 2013 | Adjacent to treated maize fields | 99.35 | 0.4 | Cutler and Scott-Dupree (2014) |
| *Bombus terrestris* | Pollen | 10<sup>th</sup> May 2014 | Adjacent to untreated WS OSR fields | *Data not collected* | <0.3 (*limit of detection*) | Rolke *et al.* (2016) |
| *Bombus terrestris* | Pollen | 10<sup>th</sup> May 2014 | Adjacent to treated WS OSR fields | *Data not collected* | 0.88 | Rolke *et al.* (2016) |
| *Osmia bicornis* | Pollen | 14<sup>th</sup> May 2014 | Adjacent to untreated WS OSR fields | *Data not collected* | <0.3 (*limit of detection*) | Rolke *et al.* (2016) |
| *Osmia bicornis* | Pollen | 14<sup>th</sup> May 2014 | Adjacent to treated WS OSR fields | *Data not collected* | 0.88 | Rolke *et al.* (2016) |

Where bees collect a greater proportion of wildflower pollen, neonicotinoid concentrations are lower. Botías *et al.* (2015) measured neonicotinoid concentrations in pollen during the peak flowering period of oilseed rape and two months after this period. During peak flowering, honeybees collected 91.1% of their pollen from wildflowers and 8.9% from oilseed rape, with a total neonicotinoid concentration of 3.09 ng/g. In the later period, 100% of their pollen was collected from wildflowers, with a total neonicotinoid concentration of 0.20 ng/g. Cutler *et al.* (2014) also sampled honeybee pollen from apiaries adjacent to treated and untreated oilseed rape for a two week period in July during peak flowering. Honeybees collected low levels of crop pollen and higher levels of neonicotinoid contamination were found adjacent to treated fields (9.0% wildflower pollen week 1 to 45.2% week 2, 0.84 ng/g) than untreated fields (15.1% wildflower pollen week 1 to 62.5% week 2, 0.24 ng/g). Long and Krupke (2016) collected data over a longer period of time, from May to September, covering the flowering period of maize, the flowering crop at their study sites. At all sites a high proportion of pollen was collected from wildflowers. Average neonicotinoid concentrations were lowest at non-agricultural sites (93.9% wildflower pollen, 0.047 ng/g), higher at untreated agricultural sites (95.8% wildflower pollen, 0.078 ng/g) and highest at treated agricultural sites (95.3% wildflower pollen, 0.176 ng/g). Alburaki *et al.* 2015 and 2016) found low levels of neonicotinoids when honeybees collected predominantly wildflower pollen, with none detected in loads of 99% wildflower pollen and average neonicotinoid concentrations of 0.04 ng/g in loads of 93.5% wildflower pollen.

Only two studies are available which measured neonicotinoid concentrations in bumblebee collected pollen and quantified the proportion of pollen collected from wildflowers. Cutler and Scott-Dupree (2014) placed out *Bombus impatiens* nests next to neonicotinoid-treated and untreated maize fields. Bumblebees collected a very low proportion of their pollen from maize, less than 1%, in contrast to honeybees which can collect large quantities of maize pollen during its flowering period (Krupke *et al.* 2012; Pohorecka *et al.* 2013, though see Alburaki *et al.* 2015; 2016; Long and Krupke 2016). Levels of neonicotinoid residues were low, at <0.1 ng/g by untreated fields and 0.4 ng/g by treated fields. In contrast, David *et al.* (2016) placed out five *B. terrestris* nests adjacent to treated oilseed rape fields, a crop with pollen attractive to bumblebees. Pollen was sampled from nest stores at the end of June. Bumblebees collected an average of 68.1% wildflower pollen and 31.9% oilseed rape pollen.

Thiamethoxam was found in this pollen at an average concentration of 18 ng/g and thiacloprid at an average concentration of 2.9 ng/g. These levels are much higher than the levels found in honeybee collected pollen from the same study area in the same year of 3.09 ng/g total neonicotinoids, though a much higher proportion (91.9%) of pollen was collected from wildflowers (Botías *et al.* 2015). Comparisons are difficult because few other studies have assessed neonicotinoid concentrations in bumblebee collected pollen with reference to pollen origin. Rolke *et al.* (2016) placed *B. terrestris* colonies out next to treated oilseed rape fields and found much lower concentrations of 0.88 ng/g of clothianidin in pollen taken directly from returning bumblebees, but the origin of this pollen is unknown. The concentrations found by David *et al.* are however lower than the levels reported by Pohorecka *et al.* (2013) and within a factor of two of the levels reported by Rundlöf *et al.* (2015) who found neonicotinoid concentrations of 27.0 ng/g and 13.9 ng/g in honeybee-collected pollen respectively, samples which also contained a high proportion of crop pollen.

Overall, these studies show that the highest acute exposure (0.84–27.0 ng/g) comes during the flowering period of insect-attractive neonicotinoid-treated flowering crops in situations where over a quarter of total pollen intake comes from crop plants. Reported values vary by up to two orders of magnitude depending on crop type, date of sample collection, initial strength of neonicotinoid seed coating and the proportion of wildflower pollen collected. Because only one study has explicitly measured neonicotinoid concentrations in wildflower pollen it is difficult to judge whether wildflower pollen consistently contains higher or lower concentrations of neonicotinoids than crop pollen. However, when looking at honeybee pollen diets in neonicotinoid-treated agricultural areas outside of the main flowering period of attractive crops, or where flowering crops are unattractive to a specific bee species, neonicotinoid concentrations are generally low, in the region of 0.04–0.40 ng/g from pollen diets comprised of 95.3–100% wildflower pollen (Cutler and Scott-Dupree 2014; Botías *et al.* 2015; Long and Krupke 2016; Alburaki et al. 2016). Whilst the highest levels of acute exposure come from pollen diets containing a proportion of crop pollen, because honeybees collect pollen over the whole season, total exposure to neonicotinoids may primarily be determined by concentrations in wildflowers. Botías *et al.* (2015) calculated, based on pollen collected in June and August, that 97% of the total neonicotinoids present in pollen were of wildflower origin. Non-crop plants surrounding agricultural areas represent an additional and chronic source of neonicotinoid exposure.

#### 2.2.5 Risk of exposure from succeeding crops

The risk of neonicotinoid exposure from succeeding crops was identified as a key knowledge gap by the EFSA reports. The available studies suggested that residues in succeeding crops are below LOQ, but the data set was limited. Since 2013, few studies have explicitly looked at neonicotinoid levels in untreated crops grown in soil that had previously been used to grow neonicotinoid-treated crops, as most crops will be sown with a new dose of neonicotinoids each year. However, where specific neonicotinoid formulations are changed this analysis is possible. Botías *et al.* 2015; 2016) analysed neonicotinoid concentrations in oilseed rape treated with thiamethoxam. The fields had been used to grow clothianidin treated cereals over at least the previous two years. Imidacloprid had not been used for the previous three years. Oilseed rape pollen and foliage was found to contain 3.15 ng/g and 1.04 ng/g of thiamethoxam, 1.90 ng/g and 2.91 ng/g of clothianidin and 0 ng/g and 0.23 ng/g of imidacloprid, respectively. As clothianidin can be produced as a metabolite of thiamethoxam it is not possible to comment on the origin of these detected residues. Imidacloprid was absent from the pollen samples, reflecting the time since the last known agricultural use. Given that these compounds can persist in soil for multiple years, the level of exposure from succeeding crops will broadly depend on the date since the last application, as well as the other factors determining neonicotinoid persistence in soil (Section 2.2.1). However, as demonstrated by the presence of imidacloprid in foliage samples, succeeding crops can take up residues of neonicotinoids remaining from applications made at least two years previously. Given the presence of neonicotinoids in annual, perennial and woody vegetation surrounding agricultural land (Section 2.2.4), and the medium-term persistence of neonicotinoids in soil and water (Sections 2.2.2 and 2.2.3), the risk of exposure from succeeding crops is likely to be in line with levels reported from general vegetation in agricultural environments. However, more explicit investigation in this area is required.

## 3. EVIDENCE FOR IMPACT OF NEONICOTINOIDS ON ANIMAL HEALTH

### 3.1 Sensitivity of bumblebees and solitary bees to neonicotinoids

#### 3.1.1 Direct lethality of neonicotinoids to adult wild bees

Almost all of the studies conducted on the toxicity of neonicotinoids to bees have been conducted on honeybees, *Apis mellifera*. Fourteen studies conducted up to 2010 were reviewed in a meta-analysis by Cresswell (2011) who concluded that for acute oral toxicity imidacloprid has a 48-h LD_50_=4.5 ng/bee. The EFSA studies (2013a; 2013b; 2013c) reviewed existing studies for acute oral toxicity up to 2013, including both peer reviewed studies and also private studies that are not in the public domain (summarised in Godfray *et al.* 2014). These analyses produced LD_50_s of 3.7 ng/bee for imidacloprid, 3.8 ng/bee for clothianidin and 5.0 ng/bee for thiamethoxam. Equivalent LD_50_s for acute contact have also been calculated by EFSA (2013a; 2013b; 2013c) for honeybees to be 81 ng/bee for imidacloprid, 44 ng/bee for clothianidin and 24 ng/bee for thiamethoxam.

However, the EFSA reports highlighted a knowledge gap for the effects of neonicotinoids on bees other than honeybees. Arena and Sgolastra (2014) conducted a meta-analysis comparing the sensitivity of bees to pesticides relative to the sensitivity of honeybees. This analysis combined data from 47 studies covering 53 pesticides from six chemical families with a total of 150 case studies covering 18 bee species (plus *A. mellifera*). Arena and Sgolastra calculated a sensitivity ratio R between the lethal dose for species *a (A. mellifera)* and for species *s* (other than *A. mellifera*), R = LD_50_*a*/LD_50_*s*. A ratio of over 1 indicates that the other bee species is more sensitive to the selected pesticides than *A. mellifera* and vice versa. There was high variability in relative sensitivity ranging from 0.001 to 2085.7, but across all pesticides a median sensitivity of 0.57 was calculated, suggesting that *A. mellifera* was generally more sensitive to pesticides than other bee species. In the vast majority of cases (95%) the sensitivity ratio was below 10.

Combining data for all neonicotinoids (acetamiprid, imidacloprid, thiacloprid and thiamethoxam) and for both acute contact and acute oral toxicity, nine studies covering nine bee species (plus *A. mellifera*) were found. These studies showed a median sensitivity ratio of 1.045 which is the highest median value of all the analysed pesticide chemical families. The most relatively toxic neonicotinoids to other bees were the cyano-substituted neonicotinoids acetamiprid and thiacloprid as these exhibit lower toxicity to honeybees than the nitro-substituted neonicotinoids imidacloprid and thiamethoxam.

Selecting pesticides covered by the moratorium (excluding acetamiprid and thiacloprid and including fipronil) and including both acute contact and acute oral toxicity, 12 studies covering 10 bee species (plus *A. mellifera*) were found. These studies showed a median sensitivity ratio of 0.957 which is close to the calculated sensitivity ratio for all neonicotinoids. The greatest discrepancy between honeybees and other bees was found for stingless bees (Apidae: Meliponini). The effect of acute contact of fipronil on *Scaptotrigona postica* (24-fold greater), of acute contact of fipronil on *Melipona scutellaris* (14-fold greater) and of acute contact of Thiacloprid on *Nannotrigona perilampoides* (2086-fold) were the only three cases with a sensitivity ratio of over 10. Stingless bees are predominantly equatorial with the greatest diversity found in the neotropics. No species are found in Europe (Nieto *et al.* 2014). In contrast, studies on *B. terrestris* consistently report a lower sensitivity ratio between 0.005 and 0.914, median 0.264. *B. terrestris* is widespread in Europe and is the most commonly used *non-Apis* model system for assessing the effects of neonicotinoids on wild bees (see Section 3.1.2). Differences in bee body weight have been proposed to explain these differences, with sensitivity to pesticides inversely correlated with body size (Devilliers *et al.* 2003). However, this has not been consistently demonstrated and other mechanisms have been suggested such as species level adaptation to feeding on alkaloid-rich nectar (Cresswell *et al.* 2012) and differential abilities to clear neonicotinoid residues from their bodies (Cresswell *et al.* 2014). With the limited data available Arena and Sgolastra could not comment on the strength of these claims.

Spurgeon *et al.* (2016) calculated various toxicity measures of clothianidin on honeybees, the bumblebee species *B. terrestris* and the solitary bee species *O. bicornis.* Acute oral toxicity 48-h, 96-h and 240-h LD_50_s for honeybees were 14.6 ng/bee, 15.4 ng/bee and 11.7 ng/bee respectively. For *B. terrestris*, the corresponding values were 26.6 ng/bee, 35 ng/bee and 57.4 ng/bee respectively. For *O. bicornis*, the corresponding values were 8.4 ng/bee, 12.4 ng/bee and 28.0 ng/bee respectively. These findings are generally in line with the findings of Arena and Sgolastra, with *B. terrestris* less sensitive than *A. mellifera* at all time points and *O. bicornis* less sensitive at 240-h.

Sgolastra *et al.* (2016) calculated relative sensitivity to clothianidin to these same three species over a range of time periods from 24–96 hours. The highest LD_50_ values were obtained after 24 hours for *A. mellifera* and *B. terrestris* and after 72 hours for *O. bicornis*. At these time points, *O. bicornis* was the most sensitive of the three species, with LD_50_ measurements of 1.17 ng/bee and 9.47 ng/g, compared to 1.68 ng/bee and 19.08 ng/g for *A. mellifera* and 3.12 ng/bee and 11.90 ng/g for *B. terrestris*. These results are in line with the values calculated by Spurgeon *et al.* (except for the 240 hour values), with decreasing sensitivity in the order of *O. bicornis* > *A. mellifera* > *B. terrestris*. Together, these studies support the position that small bodied species show greater sensitivity to neonicotinoids.

Around 2000 bee species are known from Europe. The biology, behaviour and ecology of each of these species differ from those of honeybees. Consequently, extrapolating from the limited toxicological data available for 19 bee species to the effects of neonicotinoids on the wider European fauna is fraught with difficulties given the wide variation in relative sensitivity. Current data suggests that wild bees are equally to slightly less sensitive to neonicotinoids compared to honeybees when considering direct mortality. However, care must be taken when considering individual bee species, genera and families, as different taxonomic groups may show consistently different individual level sensitivity. Most European wild bees are smaller than honeybees and there is the potential for them to be more sensitive on a ng/bee basis. In general, continuing to use honeybee neonicotinoid sensitivity metrics is likely to be a reasonable proxy measure for the direct sensitivity of the wild bee community to neonicotinoids (Arena and Sgolastra 2014), but further work is needed in this area to cover the wide range of bee species present in agricultural environments.

#### 3.1.2 Sublethal effects of neonicotinoids on wild bees

In 2013 a number of studies looking at sublethal effects of neonicotinoids were available, predominantly using honeybees as a model organism in laboratory conditions. Blacquière *et al.* (2012) reviewed studies on neonicotinoid side effects on bees published between 1995 and 2011 with a specific focus on sublethal effects. The authors found that whilst many laboratory studies described lethal and sublethal effects of neonicotinoids on the foraging behaviour and learning and memory abilities of bees, no effects were observed in field studies at field-realistic dosages. Two major studies that substantially contributed towards the initiation and subsequent implementation of the European Union neonicotinoid moratorium were published after this review in 2012.

Henry *et al.* (2012) gave honeybee workers an acute dose of 1.34 ng of thiamethoxam in a 20 μl sucrose solution, equivalent to 27% of the LD_50_ (see Section 3.1.1) then released them 1 km away from their nests and measured their return rate. Dosed bees were significantly less likely to return to the nest than control bees. Whitehorn *et al.* (2012) exposed *B. terrestris* colonies to two levels of neonicotinoid-treated pollen (6 and 12 ng/g plus control) and nectar (0.7 and 1.4 ng/g plus control) in the laboratory for two weeks before moving them outdoors to forage independently for six weeks, aiming to mimic a pulse exposure that would be expected for bees foraging on neonicotinoid-treated oilseed rape. Bees in the two neonicotinoid treatments grew significantly more slowly and had an 85% reduction in the number of new queens produced when compared to control colonies.

Both of these studies have been criticised for using neonicotinoid concentrations greater than those wild bees are likely to be exposed to in the field (see Godfray *et al.* 2014, Carreck and Ratnieks 2014) . The 1.34 ng of thiamethoxam in a 20 μl sucrose solution used by Henry *et al.* is a concentration of 67 ng/g. Taking maximum estimated concentrations of thiamethoxam in oilseed rape nectar of 2.72 ng/g (see Section 2.1.1), a honeybee would have to consume 0.49 g of nectar to receive this dose. Honeybees typically carry 25–40 mg of nectar per foraging trip, equivalent to 0.025–0.040 g, some 10% of the volume necessary to receive a dose as high as the one used by Henry *et al.* Moreover, as honeybee workers regurgitate this nectar at the hive, the total dose consumed is likely to be a fraction of the total amount carried. Consequently, it is extremely unlikely that the findings of Henry *et al.* are representative of a real world situation.

The pollen and nectar concentrations used by Whitehorn *et al.* are much closer to field-realistic levels with the lower treatment within maximum estimated concentrations of imidacloprid in oilseed rape pollen and nectar (see Section 2.1.1). However, the experimental set up, where bees had no choice but to consume treated pollen and nectar has been criticised as unrealistic, as in the real world alternative, uncontaminated forage sources would be available. Studies that have measured residues in both crop and wildflower pollen and have assessed the origin of bee-collected pollen (see Section 2.2.4) have recorded neonicotinoid concentrations of between 0.84–27.0 ng/g in wild bee-collected pollen where a substantial proportion of this pollen is collected from crop plants during their period of peak flowering. Pollen extracted from bumblebee nests contained neonicotinoid concentrations of 6.5 ng/g in urban areas and 21.2 ng/g in rural areas during the peak flowering period of oilseed rape, though the number of nests sampled (three and five) were low. However, other studies measuring levels in pollen taken directly from bumblebees found concentrations of <1 ng/g, so there is still a lack of clarity surrounding true levels of neonicotinoid exposure for wild bumblebees. On the basis of these described concentrations, the results of Whitehorn *et al.* are likely to be closer to real world conditions than the findings of Henry *et al.*

Post-April 2013, much work on sublethal effects of neonicotinoids on bees has been carried out on individual honeybees and honeybee colony fitness metrics, such as colony growth, overwintering success and the production of sexuals. This work is beyond the scope of this review, but important recent publications include Pilling *et al.* (2013), Cutler *et al.* (2014a), Rundlöf *et al.* (2015) and Dively *et al.* (2015) who all found limited to negligible impacts of neonicotinoids at the colony level. See also Cresswell (2011) for a meta-analysis of 13 laboratory and semi-field studies conducted before 2011. Various authors note that interpreting the findings of studies on honeybees to wild bees is fraught with difficulty, given the differing size of individual bees and the social behaviour of honeybees that gives rise to colonies containing many thousands of workers.

##### 3.1.2.1 Impact on colony growth and reproductive success

Several authors have investigated the effects of neonicotinoids on bumblebees using micro-colonies. These are small groups of worker bumblebees that are taken from a queenright colony and isolated in a new nest box. These workers, lacking a queen, will begin to rear their own male offspring. As such, micro-colonies are useful for generating a large sample size for investigating pesticide impacts on bee mortality and larval rearing behaviour and reproductive success.

Elston *et al.* (2013) fed micro-colonies of three *B. terrestris* workers a ‘field-realistic’ dose of 1 ng/g thiamethoxam and a ‘field-maximum’ dose of 10 ng/g in both pollen paste and sugar solution for a 28-day period. Micro-colonies from both thiamethoxam treatments consumed significantly less sugar solution than control colonies. There was no impact on worker mortality, but colonies fed 10 ng/g thiamethoxam had reduced nest-building activity and produced significantly fewer eggs and larvae, with the 10 ng/g thiamethoxam treatment the only one to produce no larvae over the 28-day experimental period.

Laycock *et al.* (2014) fed micro-colonies of four *B. terrestris* workers thiamethoxam-treated sugar solution at a range of concentrations up to 98 ng/g. Pollen was not treated with thiamethoxam. Sugar solution consumption was significantly reduced at the 39 and 98 ng/g treatments. Worker mortality was only increased at the highest dose of 98 ng/g. Worker oviposition failure was only significantly higher at the 39 and 98 ng/g treatments, with no significant differences seen between the lower concentration treatments between 0 and 16 ng/g.

The findings of these two studies are generally in line with pre-2013 knowledge. Mommaerts *et al.* (2010) exposed *B. terrestris* micro-colonies to sugar solution treated with thiamethoxam concentrations of up to 100 ng/g. Whilst the 100 ng/g level reduced brood production, the 10 ng/g treatment had no detectable effect. The difference between the findings of Elston *et al.* and Laycock *et al.* may partially be explained by the fact that Elston *et al.* treated pollen with thiamethoxam as well as sugar solution. Laycock *et al.* confirm that concentrations of 98 ng/g increase worker mortality, but as such concentrations are not usually encountered in the field this is of limited relevance.

Scholer and Krischik (2014) exposed greenhouse queenright colonies of *B. impatiens* to imidacloprid- and clothianidin-treated sugar syrup at concentrations of 0, 10, 20, 50 and 100 ng/g for 11 weeks. Queen mortality was significantly increased at six weeks for the 50 and 100 ng/g treatments, and at 11 weeks for the 20 ng/g treatment for both clothianidin and imidacloprid. Surprisingly, no significant impact was found on numbers of workers or new queens produced, though this was in part because very low numbers of new queens were produced across all treatments (average of four per colony). Colonies in treatments above 10 ng/g imidacloprid and 20 ng/g of clothianidin gained significantly less weight over the course of the study. Neonicotinoid concentrations of 20 ng/g and above are very high and are unlikely to be consistently encountered by bees for prolonged periods of times under real world conditions. As a result, queen mortality in the real world is unlikely to be significantly affected by currently observed neonicotinoid concentrations.

Several field studies have also been published since 2013 that investigate the impact of neonicotinoid-treated mass flowering crops on wild bee colony growth and reproductive success. Cutler and Scott-Dupree (2014) placed *B. impatiens* colonies adjacent to maize fields during pollen shed in Ontario, Canada. Four neonicotinoid-treated conventional and four untreated organic fields were used. Colonies were placed out adjacent to each field on the first day of major pollen shed. Colonies were left for 5–6 days and then transported to an area of semi-natural habitat for 30–35 days, after which they were frozen. Colonies placed next to treated maize produced significantly fewer workers than those placed next to organic farms. All other metrics (colony weight, honey and pollen pots, brood cells, worker weight, male and queen numbers and weights) were not significantly different. Bumblebees collected less than 1% of their pollen from maize (Section 2.2.4) and neonicotinoid residues in collected pollen were low, at 0.4 ng/g from bees foraging adjacent to treated fields and below the LOD for bees adjacent to organic fields. Given that it is well known that bumblebees collect very low volumes of maize pollen, the relevance of this study is unclear.

Rundlöf *et al.* (2015) conducted an extensive field trial of the effects of clothianidin-treated oilseed rape on wild bees. Sixteen oilseed rape fields separated by at least 4 km were selected across southern Sweden and were paired on the basis of similar landscape composition. In each pair, one of the fields was randomly selected to be sown with oilseed rape treated with 10 g clothianidin/kg of seed and the other field was sown without a neonicotinoid seed treatment. Twenty-seven cocoons of the solitary bee *O. bicornis* (15 male, 12 female) were placed out alongside each field a week before the oilseed rape began to flower, and six colonies of *B. terrestris* were placed alongside each field on the day the oilseed rape began to flower. The *O. bicornis* placed adjacent to treated oilseed rape showed no nesting behaviour and did not initiate brood cell construction. *O. bicornis* adjacent to untreated fields showed nesting behaviour in six of the eight fields studied. The reasons for these differences in nest initiation are unclear and it is difficult to draw firm conclusions with a small sample size. Bumblebees placed next to treated oilseed rape showed reduced colony growth and reproductive output. Bumblebee colonies were collected and frozen when new queens began to emerge, with this happening between the 7^th^ of July and 5^th^ of August depending on each colony. The number of queen and worker/male cocoons present was counted. At the point of freezing, colonies placed next to treated oilseed rape fields had significantly fewer queen and worker/male cocoons present.

Sterk *et al.* (2016) performed a similar field experiment to Rundlöf *et al.* Two 65 km^2^ areas in northern Germany were selected in which the only flowering crops comprised winter-sown oilseed rape. In one area the oilseed rape was treated with the same seed coating used by Rundlöf *et al.* of 10 g clothianidin/kg seed. The other area was an untreated control. In each area, ten *B. terrestris* colonies were placed at each of six localities. Colonies were left adjacent to oilseed rape between April and June, covering its main flowering period. After this the colonies were moved to a nature reserve. No differences were found in colony weight growth, number of workers produced or reproductive output as measured by the production of new queens.

That these two field studies using the same neonicotinoid seed dressing found markedly different results is interesting. The major difference is that whilst Rundlöf *et al.* used spring-sown oilseed rape, Sterk *et al.* used winter-sown oilseed rape. The length of time between sowing and peak flowering is much greater for winter-sown oilseed rape (mid-August to May) than for spring-sown oilseed rape (April/May to mid-June). As such, there is more time for neonicotinoids to leach into soil and water for winter-sown oilseed rape, reducing the amount of active ingredient available to be taken up by the crop. This may explain some of the order of magnitude differences in neonicotinoid concentrations in pollen collected from the two crops (Section 2.2.4) and the difference in reported colony growth and number of reproductives produced. An additional difference is that in the Sterk *et al.* study, colonies were moved to a nature reserve consisting of forests, lakes and heaths after the flowering period of oilseed rape ended. The quality of available forage at this nature reserve is likely to have been of both a higher quality and quantity than what was available in a conventional agricultural landscape and is not typical of the experience of a bumblebee colony located in such a landscape that will have to continue foraging there after crops such as oilseed rape cease flowering. In addition, a major problem with the experimental design of Sterk *et al.* is that only one treated and one control area were used, so there is no true site level replication, as opposed to Rundlöf *et al.* who used eight treated and eight control fields. These differences in experimental design should be taken into account when considering why the studies produced such different results.

One of the studies conducted in response to the results of Henry *et al.* (2012) and Whitehorn *et al.* (2013) was produced by FERA (2013). It consisted of a field trial with bumblebee colonies placed out adjacent to oilseed rape treated with either clothianidin, imidacloprid or an untreated control. Colonies were allowed to forage freely for 6–7 weeks whilst the oilseed rape flowered and then were moved to a non-agricultural area to continue developing. The initial aim was to measure colony growth and development across these three treatments and compare this with neonicotinoid concentrations collected from food stores within the nests, but the study was criticised for a number of methodological problems such as variable placement date and initial colony size, lack of site level replication and contamination of control colonies with neonicotinoid residues during the experiment. The study was ultimately not published in a peer reviewed journal but it came to the conclusion that there was no clear relationship between bumblebee colony success and neonicotinoid concentrations. Goulson (2015) reanalysed the FERA data using linear models and retaining two colonies excluded in the original study as outliers, but which do not meet the statistical definition of this term. This reanalysis found that the concentration of clothianidin in nectar and the concentration of thiamethoxam in pollen significantly negatively predicted both colony weight gain and production of new queens.

Only one study is available that looked at the impact of neonicotinoids on the reproductive success of a solitary bee in controlled conditions. Sandrock *et al.* (2014) established laboratory populations of *O. bicornis,* a solitary stem nesting bee. Bees were fed on sugar solution treated with 2.87 ng/g thiamethoxam and 0.45 ng/g clothianidin along with untreated pollen. There was no impact of neonicotinoids on adult female longevity or body weight. However, treated bees completed 22% fewer nests over the course of the experiment. Nests completed by treated bees contained 43.7% fewer total cells and relative offspring mortality was significantly higher, with mortality rates of 15% and 8.5% in the treated and untreated groups, respectively. Overall, chronic neonicotinoid exposure resulted in a significant reduction in offspring emergence per nest, with treated bees producing 47.7% fewer offspring. These results suggest that exposure to these low level, field-realistic doses of neonicotinoids (<3.5 ng/g) did not increase adult mortality but did have sublethal impacts on their ability to successfully build nests and provision offspring.

Overall, the studies produced since 2013 are generally in line with existing knowledge at this point but have advanced our knowledge in several key areas. Laboratory studies have continued to demonstrate negative effects of neonicotinoids on bumblebee reproductive output at generally high concentrations, with the lowest sublethal effects on reproductive output detected at 10 ng/g. Field studies using bumblebees demonstrate that exposure to neonicotinoid-treated flowering crops can have significant impacts on colony growth and reproductive output depending on the levels exposed to, with crop flowering date relative to sowing and availability of uncontaminated forage plants likely to explain variation in the detected residues between the available studies. Our understanding of the impact on solitary bees is much improved with the findings of Sandrock *et al.* (2014) suggesting substantial impacts on solitary bee reproductive output at field-realistic concentrations of 3.5 ng/g. Field studies demonstrating this under real-world conditions are limited with the work of Rundlöf *et al.* (2015) suffering from no nest-building activity at the neonicotinoid treatment sites.

##### 3.1.2.2 Impact on foraging efficiency

In 2013 a limited amount was known about how neonicotinoids affected the foraging behaviour of individual bees, and whether this affected colony level fitness. Gill *et al.* (2012) exposed *B. terrestris* colonies to 10 ng/g imidacloprid in sugar solution in the nest for a period of four weeks. Colonies were housed indoors but access tubes allowed them to forage freely outdoors. Imidacloprid exposed colonies grew more slowly but there were substantial effects on worker foraging behaviour. Compared to controls, imidacloprid treated colonies had more workers initiating foraging trips, workers brought back smaller volumes of pollen on each successful trip and successful pollen foraging trips were of a significantly longer duration. Treated workers also collected pollen less frequently, with 59% of foraging bouts collecting pollen versus 82% for control workers, a decline of 28%. The authors conclude that exposure to imidacloprid at these concentrations significantly reduced the ability of bumblebee workers to collect pollen in the field. The reduced ability to collect pollen resulted in imidacloprid treated colonies collecting less pollen than control colonies, subsequently resulting in reduced growth through pollen limitation. Since the publication of this paper, several new studies assessing neonicotinoid impacts on the foraging behaviour of bumblebees have been published.

Feltham *et al.* (2014) exposed *B. terrestris* colonies to sugar solution treated with 0.7 ng/g and pollen treated with 6 ng/g of imidacloprid for two weeks. These sugar solution concentrations were an order of magnitude lower than the 10 ng/g used by Gill *et al.* (2012). Colonies were then placed out in an urban area in Scotland. The foraging workers from each nest were then monitored for a further four weeks. There was no difference in the length of time spent collecting nectar or the volume of nectar collected between workers from treated and control colonies. However, treated workers collected significantly less pollen, bringing back 31% less pollen per time unit to their colonies. Treated workers also collected pollen less frequently, with 41% of foraging bouts collecting pollen versus 65% for control workers, a decline of 23%.

Gill and Raine (2014) performed a similar experiment to Gill *et al.* (2012) where *B. terrestris* colonies were exposed to sugar solution treated with 10 ng/g of imidacloprid whilst also having access to forage freely outside. Colonies and individual worker bumblebees were studied over a four week period. In common with their previous findings (Gill *et al.* 2012), imidacloprid treated workers initiated significantly more foraging trips across all four weeks of the experiment. The authors note that this is likely driven by an acute individual-level response in the first weeks (neonicotinoids acting as a neural partial agonist, increasing desire to forage) and by a chronic colony-level response in the latter part of the experiment, with treated colonies allocating a higher proportion of workers to pollen collection. Pollen foraging efficiency of treated workers decreased as the experiment progressed with the smallest collected pollen loads recorded in week four, suggesting a chronic effect of imidacloprid on pollen foraging ability. It is not clear whether this is as a result of individual performance deteriorating, or new emerging workers having been exposed for a greater period of time.

Stanley *et al.* (2015) exposed *B. terrestris* colonies to 2.4 or 10 ng/g thiamethoxam treated sugar solution for 13 days. Colonies were then moved to pollinator exclusion cages where they were allowed to forage freely on two varieties of apple blossom. Bees from colonies exposed to 10 ng/g spent longer foraging, visited fewer flowers and brought back pollen on a lower proportion of foraging trips compared to bees from control colonies. Stanley and Raine (2016) also exposed *B. terrestris* colonies to 10 ng/g thiamethoxam sugar solution for a nine to ten day period. At this point, colonies were moved to a flight arena provisioned with two common bird’s-foot trefoil *Lotus corniculatus* and one white clover *Trifolium repens* plants. Worker bees were individually released and their interaction with the flowers was recorded. Significantly more treated workers displayed pollen-foraging behaviour compared to control workers. However, control workers learnt to handle flowers efficiently after fewer learning visits.

Arce *et al.* (2016) placed *B. terrestris* nests out in an area of parkland for a five week period whilst also supplying them with sugar solution treated with 5 ng/g of clothianidin. The volume of sugar solution provided was estimated to be half that which colonies typically consume over the course of the experiment. No pollen was provided, so workers had to forage for this and to make up the shortfall in nectar resources. In contrast to the previous papers, only subtle changes to patterns of foraging activity and pollen collection were detected. There was no clear difference in colony weight gain between treatments or number of brood individuals. However, by the end of the experiment, treated colonies contained fewer workers, drones and gynes when compared with control colonies.

Switzer and Combes (2016) studied the impact of acute imidacloprid ingestion on sonicating behaviour of *B. impatiens.* Sonicating is a behaviour whereby a bumblebee lands on a flower and vibrates loudly to shake pollen loose from anthers. Bumblebee workers were fed a dose of 0, 0.0515, 0.515 or 5.15 ng of imidacloprid in 10 μL of sugar solution. These are equivalent to concentrations of 0, 5.15, 51.5 and 515 ng/g, with the highest volume consumed equivalent to 139% of the honeybee LD_50_, a moderate proxy for bumblebees, as bumblebees are generally less sensitive than honeybees (Section 3.1.1). Bees were then allowed to forage from tomato *Solanum lysopersicum* plants and sonicating behaviour was observed. At the lowest dose of 0.0515 ng of imidacloprid, no impact was found on wingbeat frequency, sonication frequency or sonication length. No analysis could be made for higher doses, as bees in these treatments rarely resumed foraging behaviour after ingesting imidacloprid. Given the neonicotinoid concentrations used in this study and sample size problems it is difficult to draw many conclusions other than that high levels of exposure impair bumblebee pollen foraging behaviour.

Overall, these studies suggest that exposure to neonicotinoids in nectar at concentrations of between 0.7–10 ng/g can have sublethal effects on the ability of bumblebees to collect pollen at both the individual and colony level. This shortfall in pollen and subsequent resource stress is a plausible mechanism to explain diminished colony growth and production of sexuals in the absence of increased direct worker mortality. Given that concentrations as high as 10 ng/g are at, but within, the upper limit of what bumblebees are likely to experience in the field (Section 2.1.1 and Section 2.2.4), it is likely that wild bumblebees exposed to neonicotinoids in contemporary agricultural environments suffer from a reduced ability to collect pollen, with a subsequent impact on their reproductive output.

##### 3.1.2.3 Impact on bee immune systems

Bee diseases (including both parasites and pathogens) have been implicated as the major factor affecting managed honeybee colony survival in recent years (vanEngelsdorp *et al.* 2010). Whilst most evidence for the negative effects of diseases comes from studies of honeybees, most diseases can affect a wide range of bee species. For example, the microsporidian parasite *Nosema ceranae* originates in Asia and has been spread around the world by the trade in honeybees. *N. ceranae* has now been detected in four different genera of wild bees (*Bombus*, *Osmia*, *Andrena*, *Heriades*) across Europe and the Americas (see Goulson *et al.* 2015). The spread of diseases between wild and managed bees can occur at shared flowering plants (Graystock *et al.* 2015).

Sánchez-Bayo *et al.* (2016) reviewed evidence that linked the use of neonicotinoids to the incidence and severity of bee diseases. Prior to 2013, several studies demonstrated a link between neonicotinoid exposure and increased susceptibility to diseases in honeybees (Vidau *et al.* 2011; Pettis *et al.* 2012). Exposure of honeybees infected with *N. ceranae* to imidacloprid reduced their ability to sterilise the brood, increasing the spread of *N. ceranae* within the colonies (Alaux *et al.* 2010). In addition, exposure to sublethal doses of imidacloprid or fipronil increased honeybee worker mortality due to a suppression of immunity-related genes (Aufauvre *et al.* 2014). Di Prisco *et al.* (2013) found that sublethal doses of clothianidin adversely affected honeybee antiviral defences. By enhancing the transcription of the gene encoding a protein that inhibits immune signalling activation, the neonicotinoid pesticides reduce immune defences and promote the replication of deformed wing virus in honeybees bearing covert viral infections. At the field level, a positive correlation is found between neonicotinoid treatment and Varroa mite infestation and viral load of honeybee colonies (Divley *et al.* 2015; Alburaki *et al.* 2015). No studies are available that measure the impact of neonicotinoids on the immune systems of wild bees or on the incidence of diseases in wild bees in conjunction with neonicotinoid usage. However, given that wild bees share a very similar nervous and immune system it is highly likely that neonicotinoids will have similar effects, increasing wild bee susceptibility to parasites and pathogens.

#### 3.1.3 Population level effects of neonicotinoids on wild bees

Nothing was known about the population level effects of neonicotinoids on wild bees in 2013. As a managed domesticated species, population trends are available for honeybees, but no such data are available for wild bees. One study has attempted to investigate the impact of neonicotinoids on wild bee population trends. Woodcock *et al.* (2016) used an incidence dataset of wild bee presence in 10 x 10 km grid squares across the United Kingdom. The dataset is comprised of bee sightings by amateur and professional entomologists and is probably the most complete national bee distribution database currently in existence. Sixty-two wild bee species were selected and their geographic distance and persistence over an 18 year period between 1994 and 2011 was calculated. Neonicotinoid seed-treated oilseed rape was first used in the UK in 2002, and so the authors calculated spatially and temporally explicit information describing the cover of oilseed rape and the area of this crop treated with neonicotinoids. The 62 species were split into two groups – species that foraged on oilseed rape (n=34) and species that did not (n=28). Species persistence across this time period was then compared with expected neonicotinoid exposure. Over the 18 year period, wild bee species persistence was significantly negatively correlated with neonicotinoid exposure for both the foraging and non-foraging group, with the effect size three times larger for the oilseed rape foraging group.

The characterisation of bees as foragers or non-foragers has one major problem. Many species of bees are obligately parasitic on other bees and do not forage for their own pollen. Some parasitic bees were included in the oilseed rape forager category (n=2), and some in the non-forager category (n=12) based on observed nectar visits from a previous study. Some of the parasitic bees in the non-forager group are parasitic on bees included in the forager group (n=10/28). Given that these species are highly dependent on their host’s abundance this classification does not make ecological sense. A decline due to a decline in their host or because of increased direct mortality cannot be separated, introducing an additional confounding issue into the analysis. In addition, given the presence of neonicotinoids in wild plants adjacent to agricultural areas (Section 2.2.4), the amount applied to oilseed rape is not necessarily a true measure of actual neonicotinoid exposure for wild bees.

Overall, the study suggests that bee species were more likely to disappear from areas with a high exposure to neonicotinoids as measured by the amounts applied as seed dressings to oilseed rape, and that this trend was more pronounced for species known to forage on oilseed rape. Whilst more work is needed, this is a major correlational study that suggests a link between levels of neonicotinoid exposure and bee community persistence at a national scale.

### 3.2 Sensitivity of butterflies and moths to neonicotinoids

Pisa *et al.* (2015) reviewed the existing literature on the impact of neonicotinoids on butterflies and moths (Lepidoptera). In contrast to bees, very few comparative toxicity tests have been conducted for butterflies. Most existing studies have compared butterfly abundance and diversity on organic versus conventional farms. Organic farms host a greater diversity of species, but the specific reasons for this cannot be isolated. For example, the relative importance of herbicide use that reduces the abundance of larval food and adult nectar plants versus direct mortality or sublethal stress from pesticides is unknown.

Most available toxicological studies looking at the sensitivity of Lepidoptera to neonicotinoids and fipronil have been conducted on 32 species of moths from nine families that are pests of crops (Pisa *et al.* 2015). There is considerable variation in reported sensitivities between species, with the susceptibility to acetamiprid of two cotton pests differing almost 3-fold (LC50=11,049 and 3,798 ppm). There is also variation between different stages of larval development, with first instar caterpillars more than 100 times as sensitive as fifth instar caterpillars with a LC_50_/LC_90_ of 0.84/1.83 and 114.78/462.11 ppm, respectively. Botías *et al.* (2016) listed LC_50_ values for three moth species that are agricultural crop pests, with 24 h LC_50_ values between 2400 and 186,000 ppb clothianidin. These levels are generally very high and there are multiple examples of neonicotinoid resistance in wild populations (see Pisa *et al.* 2015). Because many of the studied moths species are pests of major crops they have been exposed to multiple pesticides over many generations in recent decades, and their sensitivity to neonicotinoids many not necessarily be representative of non-pest wild Lepidoptera species.

Since 2013, few studies looking at the sensitivity of wild Lepidoptera to neonicotinoids are available. Pecenka and Lundgren (2015) assessed the lethality of clothianidin to caterpillars of monarch butterflies *Danaus plexippus*. First instar caterpillars were fed treated leaves for a 36 hour period. A LC_50_ of 15.63 ng/g was calculated. In addition, sublethal effects on growth were measured at 0.5 ng/g with first instar larvae taking longer to develop, having reduced body length and lower weight. These differences did not extend into the second instar. Yu *et al.* (2015) fed second instar silkworm *Bombyx mori* caterpillars leaves treated with imidacloprid and thiamethoxam for a 96 hour period. They calculated LC_50_ values of 1270 ng/g for imidacloprid and 2380 ng/g for thiamethoxam. This wide range of reported tolerances for a limited number of ecologically different species means that thorough assessment of butterfly and moth sensitivity to neonicotinoids is difficult. Much more research is required in this area.

Whilst there is a paucity of toxicological data on wild butterflies and moths, two recent studies have used long term butterfly population datasets to assess the relative impact of neonicotinoid usage in agricultural areas. Gilburn *et al.* (2015) used data from the UK butterfly monitoring scheme. The data consists of butterfly counts from a wide variety of habitats and the period studied was 1984–2012, a more extensive time period that than used for UK wild bees by Woodcock *et al.* 2016, Section 3.1.3) in order to have a ten year period before the introduction of neonicotinoids onto British farmland. Seventeen UK butterfly species were selected that are predominantly generalists and are found in a wide range of habitats including agricultural habitats. The area of the UK treated with neonicotinoids and a range of temperature and weather variables were included in the model, as local climatic conditions are a very important factor impacting butterfly populations. In line with expectations, summer temperature was significantly positively and spring rainfall significantly negatively correlated with the butterfly population indexes. Neonicotinoid usage was also significantly negatively associated with butterfly population indices after controlling for the effects of weather. The pattern of association varied between butterfly species, but most (14 out of 17) had a negative association. In the most recent time period between 2000–2009 when neonicotinoid usage was at its highest, 15 of the 17 studied species showed a negative population trend.

Forister *et al.* (2016) conducted a similar analysis on Californian lowland butterfly populations. Butterflies have been monitored continuously with biweekly walks at four sites in a region of northern California since 1972, 1975 and 1988 depending on the individual site. These sites are situated across a land gradient that includes arable, semi-natural and urban habitats. The data were used to examine the impact of annual neonicotinoid input and other factors such as summer temperature and land-use change.

A substantial decline in butterfly species richness was seen from 1997 onwards (Figure 10a, 1997 being the breakpoint identified by the statistical models). Neonicotinoid usage in the region began in 1995 and has increased since that point (Figure 10b). Neonicotinoid use was significantly negatively correlated with butterfly species richness (Figure 10c) and smaller bodied butterflies had the strongest negative response to neonicotinoids (Figure 10d).

**Figure 10.**
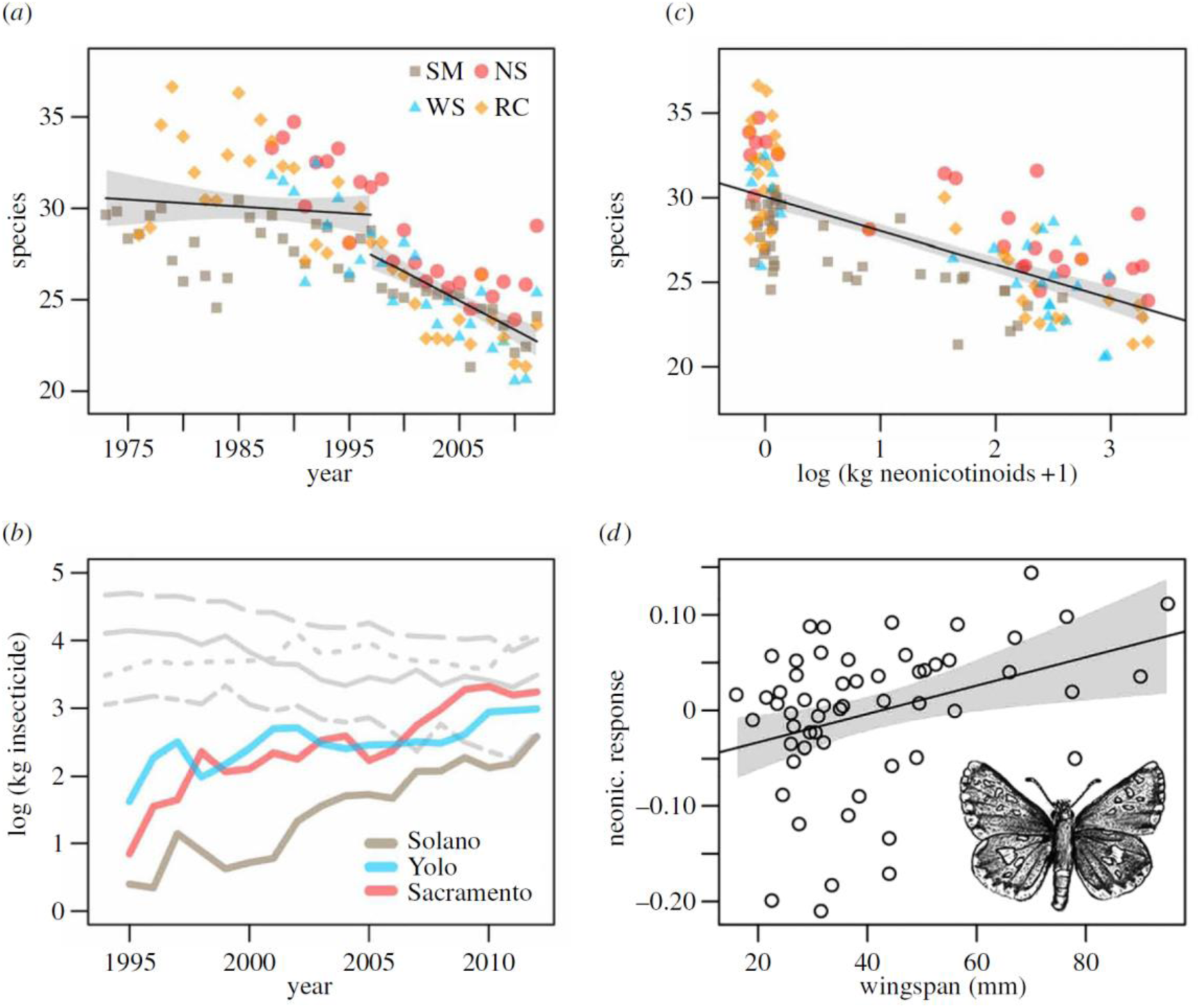
(a) The number of observed butterfly species at four sites. The response variable (in (a) and (c)) is the exponential of Shannon diversity, i.e. the effective number of species; the spline knot in (a) is 1997 (95% confidence interval: 1990–2001). (b) Pesticide application for neonicotinoids in focal counties (coloured lines), and for the four most commonly applied non-neonicotinoid classes (grey lines). The non-neonicotinoids are, in decreasing order of line elevation in 1995; organophosphates, carbamates, pyrethroids and organochlorines (lines are county averages). Note the different range of years in the first two panels, as (b) starts in the year in which neonicotinoids are first reported. (c) Relationship between number of butterfly species and neonicotinoids (values of the latter at zero jittered for visualization). (d) Response of individual species to neonicotinoids as predicted by wingspan; more negative values on the y-axis indicate species with more negative associations with neonicotinoids. Grey polygons in panels (a), (c), and (d) are 95% confidence intervals. Reproduced from Forister *et al.* 2016.

Both of these analyses are strictly correlational and neonicotinoid usage may simply be a proxy measurement for some other factor that is driving declines. Gilburn *et al.* note that if habitat deterioration and loss of food plants is the main cause of butterfly declines, and agricultural intensification is playing a key role in this habitat deterioration, then levels of neonicotinoid usage might be acting as a proxy for agricultural intensification and therefore habitat deterioration. Thus, neonicotinoid usage could be responsible for driving butterfly declines or alternatively it could provide the first useful quantifiable measure of agricultural intensification that strongly correlates with butterfly population trends. As most of the UK butterfly monitoring scheme survey areas are not directly on agricultural land, Gilburn *et al.* suspect that it is the transport of neonicotinoids into the wider environment (Section 2.2.4) and farmed areas acting as population sinks that is driving the declines of butterflies, rather than neonicotinoid use acting as a proxy for agricultural intensification. No data is available to assess this hypothesis.

Overall, recent studies have demonstrated that Lepidoptera show a wide range of tolerances to ingested neonicotinoids in their larval stages. No data is available on sensitivity to neonicotinoids ingested during the adult stage, for example from crop plant nectar. Two correlational studies using long term datasets show a strong association between neonicotinoid use and declines in butterfly abundance and species-richness, though more laboratory and field studies are required to establish the exact mechanism causing this decline.

### 3.3 Sensitivity of other terrestrial invertebrates to neonicotinoids

Most available studies that have assessed neonicotinoid sensitivity for insect species have focussed on pest species of economically important crops. Pisa *et al.* (2015) reviewed existing literature on the impacts of neonicotinoids on other terrestrial invertebrates and Botías *et al.* (2016) presented a summary on reported LC_50_s for 24 species of insects across four orders (Hymenoptera, Lepidoptera, Hemiptera and Coleoptera) from studies conducted between 1996 and 2015. Pisa *et al’s.* (2015) review found no post-2013 research on the effects of neonicotinoids on Neuroptera, Hemiptera and Syrphidae (hoverflies).

#### 3.3.1 Sensitivity of natural enemies of pest insects

Douglas *et al.* (2015) investigated the impact of thiamethoxam seed-treated soybean on the agricultural pest slug *Deroceras reticulatum* and one of their natural predators, the carabid beetle *Chlaenius tricolor*, using both laboratory assays and field studies. Slugs collected from the field that had been allowed to feed freely on developing soybean seedlings contained total neonicotinoid concentration as high as 500 ng/g with average levels over 100 ng/g after 12 days of feeding. In the laboratory, slugs consuming soybean seedlings incurred low mortality of between 6–15% depending on the strength of the seed treatment. Under laboratory conditions, 61.5% (n=16/26) of *C. tricolor* beetles that consumed slugs from the neonicotinoid treatment subsequently showed signs of impairment compared to none of those in the control treatment (n=0/28). Of the 16 that showed impairment, seven subsequently died. In the field, seed-treated soybean reduced potential slug predator activity-density by 31% and reduced predation by 33%, resulting in increased slug activity-density by 67%.

Douglas *et al.* argue that the introduction of neonicotinoids into soybean results in a trophic cascade, whereby the predators of slugs are more significantly affected than the slugs themselves, resulting in an increase in the slug population as predation pressure is relaxed. This trophic cascade argument may also explain the results of Szczepaniec et al. (2011) who found that the application of imidacloprid to elm trees caused an outbreak of spider mites *Tetranychus schoenei*. This increase was as a result of a reduction in the density of their predators which incurred increased mortality after ingesting imidacloprid-containing prey items. Many beneficial predatory invertebrates feed on pests of crops known to be treated with neonicotinoids, but to date no other studies have assessed whether neonicotinoids are transmitted to these predators through direct consumption of crop pests in agro-ecosystems.

Frewin *et al.* (2014) studied the impact of imidacloprid and thiamethoxam seed-treated soybean on the soybean aphid parasitoid wasp *Aphelinus certus*. Mated females were placed in petri dishes containing soybean leaves with soybean aphid *Aphis glycines* populations for 24 hours. Petri dishes were then monitored for eight days with the numbers of alive, dead and juvenile aphids recorded. The effects of pesticide treatment was significant on the proportion of aphids parasitised, with no difference between the two different neonicotinoid seed treatments (Figure 11). Frewin *et al.* hypothesise two potential reasons for this effect – firstly that exposure to neonicotinoid residues within aphid hosts may have increased mortality of the immature parasitoid or the parasitism combined with residues may have increased aphid mortality. Secondly, *A. certus* may avoid parasitising pesticide-poisoned aphids. *Aphelinus* species are known to use internal cues to determine host suitability, and it is possible that they may use stress- or immune-related aphid hormones to judge host suitability. Given that a key part of biological control of insect pests using parasitic wasps is to increase the parasitoid abundance early in the season, the reduction in the parasitism rate caused by neonicotinoid seed-treatment could potentially impair the ability of *A. certus* to control soybean aphid. It is not known if *A. certus* emerging from contaminated hosts will incur lethal or sublethal effects which may further impair this ability.

**Figure 11.**
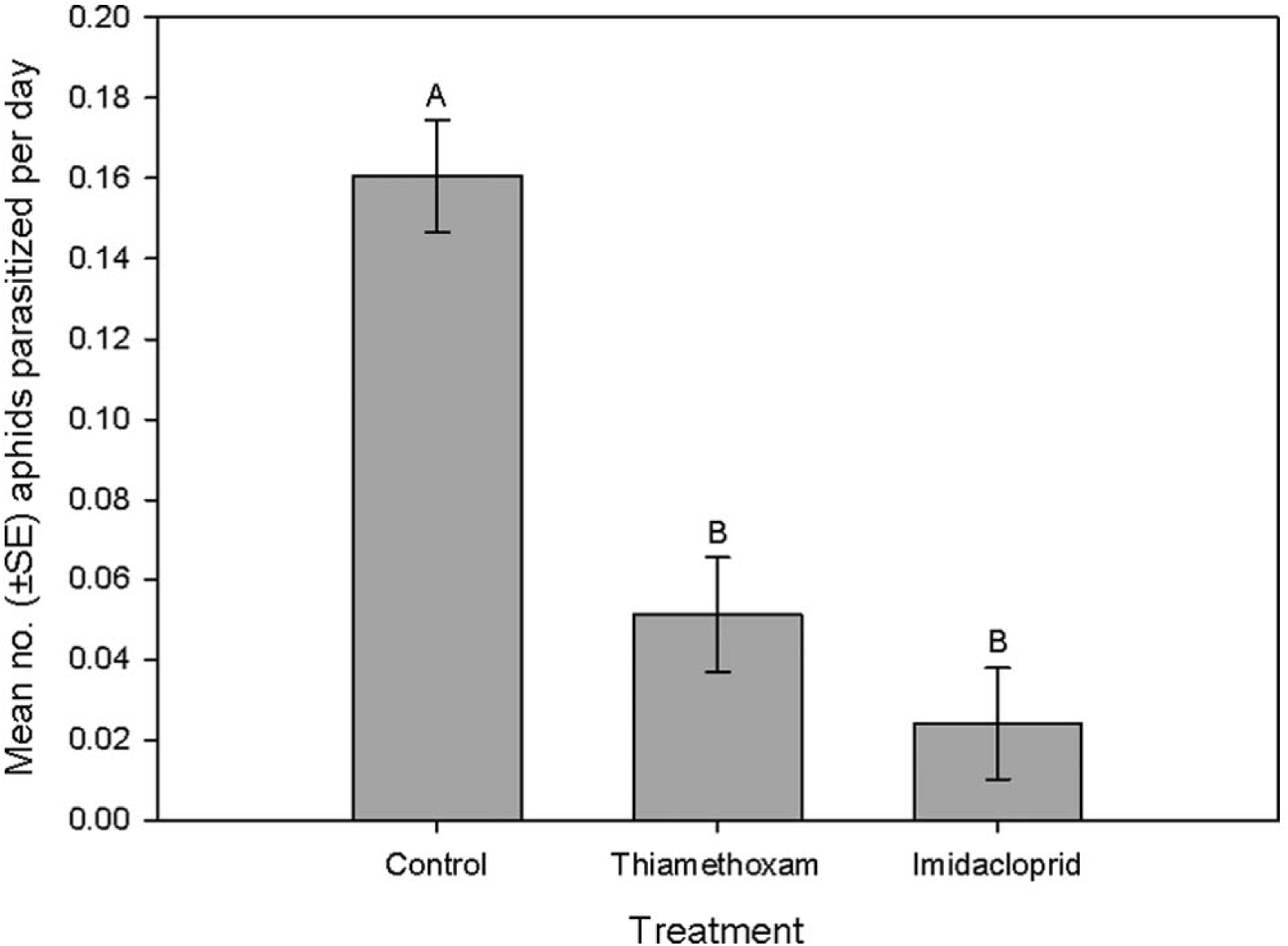
Parasitism rates (±SE) of *Aphelinus certus* on *Aphis glycines* feeding on soybean plants grown from seed not treated (control) with insecticidal seed treatment compared with those feeding on plants grown from seed treated with imidacloprid or thiamethoxam. Bars with the same letter are not significantly different (Tukey’s honestly significant difference, α = 0.05), n=35 for each treatment. Reproduced from Frewin *et al.* 2014.

Overall, where predatory species have a greater sensitivity to neonicotinoids than their prey species, such as insect predators of non-insect groups like molluscs and arachnids which have differing neuroreceptors that renders them less sensitive to neonicotinoids, there is the possibility of unintended negative effects on populations of beneficial natural enemies.

#### 3.3.2 Sensitivity of ants to neonicotinoids

Four studies are available that have looked at the impact of neonicotinoids on ants. Galvanho *et al.* (2013) treated *Acromyrmex subterraneus* leafcutter ants with imidacloprid to investigate impacts on grooming, an important behaviour for limiting the spread of fungal pathogens. Workers were treated with 10, 20 or 40 ng/insect imidacloprid. Only workers with a head capsule of 1.6–2.0 mm in width were selected. This is a large size relative to most species of ants in the world. At this size, individual ants would weigh around 10–20 mg, giving a concentration of 10–40 ng active ingredient per 0.015 g of ant, or 666.7–2666.7 ng/g. The lowest dose was sufficient to significantly decrease grooming behaviour. Mortality was not measured, but a previous study found that another species of leaf-cutter ant, *Atta sexdens*, had significantly increased mortality when exposed to a fungal pathogen and imidacloprid at the same concentration 10 ng/insect concentration compared to ants exposed only to the fungal pathogen (Santos *et al.* 2007).

Barbieri *et al.* (2013) exposed colonies of the Southern ant *Monomorium antarcticum* (native to New Zealand where the study was conducted) and the invasive Argentine ant *Linepithema humile* to imidacloprid in sugar water at a concentration of 1.0 μg/ml, equivalent to 1000 ng/g. Relative aggression was affected by neonicotinoid exposure, with native ants lowering their aggression to invasive ants, and conversely exposed invasive ants increasing their aggression, resulting in a lower survival probability. Brood production was not affected in the Southern ant, but exposure to neonicotinoids reduced Argentine ant brood production by 50% relative to non-exposed colonies. No effect of neonicotinoid exposure on foraging ability was detected.

Wang *et al.* (2015a) fed colonies of fire ants *Solenopsis invicta* sugar water at concentrations of 0.01, 0.05, 0.25, 0.50 and 1.00 μg/ml, equivalent to 10–1000 ng/g. The impact on feeding, digging and foraging were quantified. Ants exposed to the 10 ng/g concentration consumed significantly more sugar water and increased digging activity. Concentrations greater than or equal to 250 ng/g significantly supressed sugar water consumption, digging and foraging behaviour.

Wang *et al.* (2015b) fed *Solenopsis invicta* newly mated queens water containing imidacloprid concentrations of 10 or 250 ng/g. Neither concentration increased queen mortality but they did both significantly reduce queen’s brood tending ability and the length of time taken to respond to light, an indication of disturbance and colony threat. In *Solenopsis* species, eggs are groomed and coated with an adhesive substance that maintains moisture levels and allows for rapid transport of egg clumps. At the 250 ng/g concentration, the number of egg clumps was significantly increased (indicating low egg care and an increase in the effort needed to transport brood), suggesting that the queens had a reduced ability to groom eggs. Untended eggs become mouldy, reducing colony growth. Colonies exposed to 10 ng/g showed no difference in egg clump numbers compared to controls.

Across these ant studies, the neonicotinoid concentrations used are generally very high, in most cases far higher than expected exposure rates under field-realistic conditions (Section 2.1 and 2.2). Few sublethal effects were detected at 10 ng/g, the levels that might be reasonably expected to be encountered under field conditions. More laboratory and field work is required using lower concentrations to better understand the likely effects of neonicotinoids on ants.

#### 3.3.3 Sensitivity of earthworms to neonicotinoids

Pisa *et al.* (2015) reviewed existing literature on the impact of neonicotinoids on earthworms. Earthworms have similar neural pathways to insects, and earthworms are highly likely to be exposed to neonicotinoids through direct contact with soil, ingestion of organic material bound to neonicotinoids and consumption of contaminated plant material (Wang *et al.* 2012, Section 2.2.1) Reported neonicotinoid LC_50_s for earthworms from 13 studies range from 1,500 to 25,500 ppb, with a mean of 5,800 ppb and a median of 3,700 ppb (see Pisa *et al.* 2015). Fewer studies are available that measured sublethal effects on reproduction. Negative impacts on cocoon production were measured at between 300–7,000 ppb depending on earthworm species and neonicotinoid type. Very little data is available for realistic neonicotinoid exposure to earthworms under field conditions. Neonicotinoid concentrations in soils can range from 2–50 ng/g depending on organic matter composition, application rate and other factors, although they may be much higher in immediate proximity to dressed seeds (Section 2.2.1). Douglas *et al.* (2015) detected neonicotinoids in earthworms present in thiamethoxam-treated soybean fields. Two earthworms were casually collected during soil sample collection. The two samples were found to contain total neonicotinoid concentrations of 54 and 279 ppb corresponding to ~16 and ~126 ng per worm. In addition to thiamethoxam and its degradates, the two earthworm samples contained imidacloprid at 25 and 23 ppb. The fields from which they were taken had not been treated with imidacloprid for at least one year previously, adding further to the evidence that neonicotinoids can persist in soils for over one year (Section 2.2.1). Because only live earthworms were collected and the small sample size, it is not clear if these are representative of typical concentrations or are an underestimate. For example, if earthworms are exposed to higher levels that cause mortality, they cannot be subsequently sampled for residue analysis. More work is needed in this area.

Overall, these studies continue to increase our understanding of the negative effects of neonicotinoids on non-target organisms. In contrast to bees, most studied groups had lower sensitivity to neonicotinoids, in some cases by several orders of magnitude. The trophic level of the study organism may be important, with low trophic level insects better able to detoxify neonicotinoids due to their obligately herbivorous lifestyle that results in frequent contact with harmful plant metabolites. The most pronounced reported effects have been on predatory insects.

### 3.4 Sensitivity of aquatic invertebrates to neonicotinoids

The most comprehensive review of the acute and chronic effects of neonicotinoids on aquatic invertebrates was conducted by Morrissey *et al.* (2015). This followed on from and updated the reviews of Goulson (2013), Mineau and Palmer (2013) and Vijver and van den Brink (2014). Morrissey’s analysis covered 214 toxicity tests for acute and chronic exposure to imidacloprid, acetamiprid, clothianidin, dinotefuran, thiacloprid and thiamethoxam for 48 species of aquatic invertebrate species from 12 orders (Crustacea: Amphipoda (11.7% of tests), Cladocera (21.0%), Decapoda (1.9%), Isopoda (4.2%), Mysida (7.9%), Podocopida (12.6%), Insecta: Diptera (22.9%), Ephemeroptera (6.5%), Hemiptera (3.7%), Megaloptera (1.9%), Odonata (1.9%), Trichoptera (3.3%)) from peer reviewed and government studies. Both LC_50_ and ED_50_ values were included. Acute and chronic toxicity of neonicotinoids vary greatly across aquatic invertebrates with differences of six orders of magnitude observed (Figure 12). In general, insects were more sensitive than crustaceans, in particular the Ephemeroptera (mayflies), Trichoptera (caddisflies) and Diptera (flies, most specifically the midges, Chironomidae) were highly sensitive.

**Figure 12.**
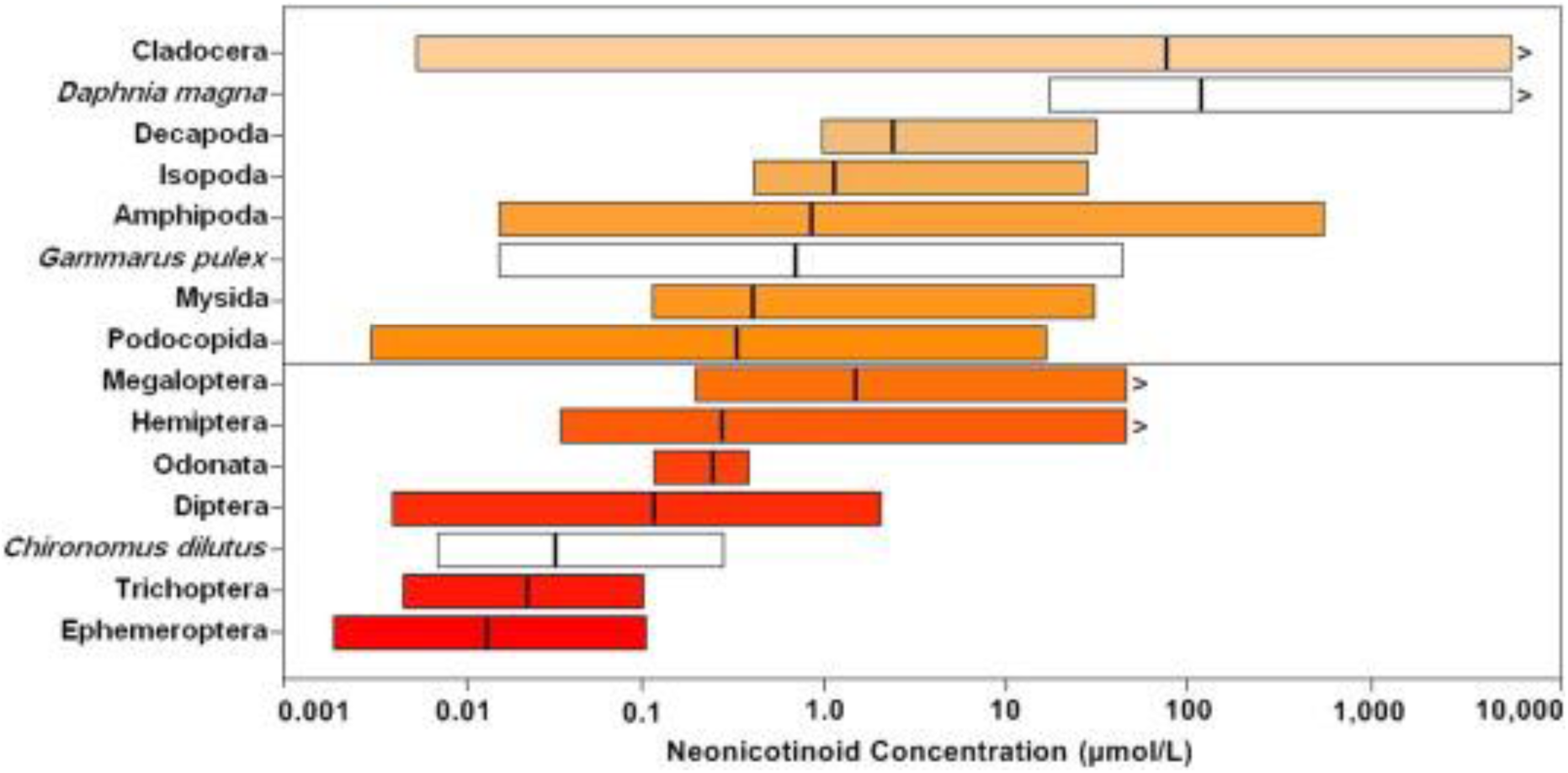
Range of neonicotinoid toxicity (L[E]C_50_: 24–96 h in μmol/L, both lethal and sublethal values included) among all tested aquatic invertebrate orders. For context, three of the most common test species (white bars) for the orders Cladocera (*Daphnia magna*), Amphipoda (*Gammarus pulex*) and Diptera (*Chironomus dilutus*) are shown to illustrate differences in sensitivity by species. Vertical lines within bars represent geometric means of test values. Concentrations are given as molar equivalents μmol/L to standardise for the variable molecular weights of the different neonicotinoids. Back conversions to concentrations in μg/L (ppb) can be obtained by multiplying the molar concentration by the molar weight of the neonicotinoid compound. Reproduced from Morrissey *et al.* 2015.

The Cladoceran water flea *D. magna* was the most commonly used model organism, represented in 34 of the 214 toxicity tests (16%). Its widespread use is because of its position as a global industry standard for the majority (82%) of commercial chemicals tested (Sánchez-Bayo 2006). It shows a wide variation in sensitivity to neonicotinoids but the mean short term L[E]C_50_ is at least two to three orders of magnitude greater than for all other tested invertebrate groups (Figure 12). This has been highlighted by several authors (e.g. Beketov and Liess 2008) who argue that given the low sensitivity of *D. magna* to neonicotinoids, a different model organism such as a Dipteran should be selected when conducting tests on this class of pesticide. This is illustrated by the most recent study to calculate LC_50_s for a range of aquatic invertebrates that was not included in Morrissey’s review. de Perre *et al.* (2015) found no sublethal or lethal effects of clothianidin on *D. magna* at concentrations of over 500 μg/L. In contrast, *C. dilutus* showed EC_50_ effects at 1.85 μg/L and LC_50_ effects at 2.32 μg/L, in line with previous findings (Figure 12).

Kunce *et al.* (2015) also investigated the impacts of neonicotinoids on the similar *C. riparius.* First instar midge larvae were exposed to thiacloprid and imidacloprid at 50% of the 96-h LC_50_s reported in the literature, corresponding to 2.3 μg/L for thiacloprid and 2.7 μg/L for imidacloprid. Three day old larvae were pulse exposed to these concentrations for 1 hour then transferred to clean water and allowed to develop normally. The one hour exposure to thiacloprid significantly decreased the proportion of larvae surviving to adulthood from 94% in the control to 68%. However, imidacloprid alone and thiacloprid and imidacloprid combined had no observable effect. No difference on adult egg production levels was detected.

These recent studies in conjunction with the review of Morrissey *et al.* strongly support the position that insect larvae are most sensitive to neonicotinoids in aquatic environments. Morrissey *et al.* conclude that chronic neonicotinoid concentrations of over 0.035 μg/L or acute concentrations of over 0.200 μg/L can affect the most sensitive aquatic invertebrate species. This finding is consistent with the value suggested by Vijver and van der Brink (2014) of 0.013–0.067 μg/L for imidacloprid. A number of water quality reference values have been published by governmental regulatory bodies and independent researchers in Europe and North America (Table 8). Most of these studies are based on assessments for imidacloprid only. Values for acceptable long term concentrations vary by three orders of magnitude from 0.0083 μg/L in the Netherlands (RIVM 2014; Smit *et al.* 2014) to 1.05 μg/L in the USA. There is considerable difference in the methodologies used to calculate these reference values, with the US EPA value likely to have been strongly based on results from *D. magna*, a species known to have relatively low sensitivity to neonicotinoids (Morrissey *et al.* 2015).

**Table 8.** Summary of published ecological quality reference values for neonicotinoids (imidacloprid except this review) in freshwater environments against which average (chronic or long-term) or maximum (acute or peak) exposure concentrations are to be compared. Reference values are placed in descending order. Reproduced from Morrissey *et al.* (2015)

| Source | Average concentration (µg/L) | Maximum concentration (µg/L) | Justification |
| --- | --- | --- | --- |
| EPA (2014) USA | 1.05 | 35.0 | Aquatic life benchmark – methodology uncertain |
| CCME (2007) Canada | 0.23 |  | EC<sub>15</sub> for the most sensitive of two freshwater species tested with assessment factor of 10 applied |
| EFSA (2008) European Union |  | 0.2 | No Observable Effect Concentration (NOEC) (0.6 µg/L) from a 21 d German microcosm study to which an assessment factor of 1–3 has been applied based on expert deliberations |
| RIVM (2008) Netherlands | 0.067 |  | Maximum permissible concentration (MPC) for long term exposure derived from the lowest NOEC value for chronic toxicity studies with assessment factor of 10 applied |
| Morrissey *et al.* (2015) | 0.035 | 0.2 | Lower confidence interval of HC<sub>5</sub> from SSDs generated using 137 acute (LC<sub>50</sub>) and 36 chronic (L[E]C<sub>50</sub>) toxicity tests considering all neonicotinoid compounds weighted and standardized to imidacloprid and all available test species |
| RIVM (2014) Netherlands (see Smit *et al.* 2014) | 0.0083 |  | Updated MPC for long-term exposure derived from chronic studies using species sensitivity distribution (SSD) approach and Hazard Concentration (HC<sub>5</sub>) applied to NOEC/LC<sub>10</sub>/EC<sub>10</sub> values with assessment factor of 3 applied |
| Mineau and Palmer (2013) | 0.0086 or 0.029 |  | The higher of two empirically-determined acute–chronic ratios applied to the most sensitive of 8 aquatic species tested to date; or HC<sub>5</sub> from SSD applied using NOECs from chronic studies of 7 single species and 1 species assemblage |

Current levels of neonicotinoids in aquatic habitats regularly exceed this threshold. Morrissey *et al.* reviewed 29 studies from nine countries and found geometric mean surface water concentrations of 0.130 μg/L (73.6%, 14/19 studies over 0.035 μg/L threshold) with geometric mean peak surface water concentration of 0.630 μg/L (81.4% 22/27 studies over 0.200 μg/L). Studies published since 2015 that are not included in Morrissey's review have also reported average neonicotinoid levels exceeding this threshold (see Section 2.2.2). Qi et al. (2015) and Sadaria *et al.* (2016) found levels of neonicotinoids above the threshold in influent and effluent wastewater at processing plants in the China and the USA. Benton *et al.* (2015) found average and peak imidacloprid levels above the thresholds in Appalachian streams in the USA. In contrast, low average levels of neonicotinoids were found in standing water and ditches on arable land in Ontario, Canada (Schaafsma *et al.* 2015) and in Iowan wetlands in the USA (Smalling *et al.* 2015). de Perre *et al.* (2015) found peak concentrations of 0.060 μg/L of clothianidin in groundwater below maize fields shortly after crop planting. In a nationwide study, Hladik and Kolpin (2016) found arithmetic mean neonicotinoid concentrations in streams across the USA to be just below the chronic threshold at 0.030 μg/L. However, peak concentration was 0.425 μg/L. Székács *et al.* (2015) also conducted a nationwide survey of Hungarian watercourses, finding clothianidin at concentrations of 0.017–0.040 μg/L and thiamethoxam at concentrations of 0.004–0.030 μg/L. The highest concentrations, of 10–41 μg/L, were only found in temporary shallow waterbodies after rain events in early summer.

Combining these recent studies with those included in Morrissey’s 2015 review a total of 65.3% of studies (17/26) report average neonicotinoid concentrations of over the 0.035 μg/L chronic threshold and 73.5% of studies (25/34) report peak concentrations over the 0.200 μg/L acute threshold. The number of countries that have been studied and their widespread distribution (Australia, Brazil, Canada, China, Hungary, Japan, the Netherlands, Sweden, Switzerland, the United States and Vietnam) indicates the widespread contamination of watercourses of all kinds with levels of neonicotinoids known to be harmful to sensitive aquatic invertebrates. This is now a chronic global problem, likely to be impacting significantly on aquatic insect abundance and on food availability for their predators, including fish, birds and amphibians.

### 3.5 Sensitivity of birds and bats to neonicotinoids

Gibbons *et al.* (2015) reviewed the direct and indirect effects of neonicotinoids and fipronil on vertebrate wildlife including mammals, fish, birds, amphibians and reptiles. LD_50_ values for imidacloprid, clothianidin and fipronil are available for 11 species of bird (Table 9). There is considerable variation in the lethality of these compounds to birds, both between bird species and pesticide type. Using US EPA (2012) classifications for toxicity (see legend for Table 9), imidacloprid ranged from moderately toxic to highly toxic, clothianidin from practically non-toxic to moderately toxic and fipronil from practically non-toxic to highly toxic.

**Table 9.** Single (acute) dose LD_50_ for bird species (mg/kg, equivalent to ppm) for imidacloprid, clothianidin and fipronil. Toxicity classification follows US EPA (2012): *PNT* practically non-toxic, *ST* slightly toxic, *MT* moderately toxic, *HT* highly toxic, *VHT* very highly toxic. For birds: *PNT* >2,000, *ST* 501–2,000, *MT* 51–500, *HT* 10–50, *VHT* <10. Reproduced from Gibbons *et al.* (2015)

| Species | Pesticide | LD<sub>50</sub> | Reference |
| --- | --- | --- | --- |
| Mallard, *Anas platyrhynchos* | Imidacloprid | 283 (MT) | Fossen (2006) |
| Grey partridge, *Perdix perdix* | Imidacloprid | 13.9 (HT) | Anon (2012) |
| Northern bobwhite quail, *Colinus virginianus* | Imidacloprid | 152 (MT) | SERA (2005) |
| Japanese quail, *Coturnix japonica* | Imidacloprid | 31 (HT) | SERA (2005) |
| Feral pigeon, *Columba livia* | Imidacloprid | 25-50 (HT) | SERA (2005) |
| House sparrow, *Passer domesticus* | Imidacloprid | 41 (HT) | SERA (2005) |
| Canary, *Serinus canaria* | Imidacloprid | 25-50 (HT) | SERA (2005) |
| Mallard, *Anas platyrhynchos* | Clothianidin | >752 (ST) | European Commission (2005) |
| Northern bobwhite quail, *Colinus virginianus* | Clothianidin | >2,000 (PNT) | Mineau and Palmer (2013) |
| Japanese quail, *Coturnix japonica* | Clothianidin | 423 (MT) | Mineau and Palmer (2013) |
| Mallard, *Anas platyrhynchos* | Fipronil | 2,150 (PNT) | Tingle *et al.* (2003) |
| Ring-necked pheasant, *Phasianus colchicus* | Fipronil | 31 (HT) | Tingle *et al.* (2003) |
| Red-legged partridge, *Alectoris rufa* | Fipronil | 34 (HT) | Tingle *et al.* (2003) |
| Northern bobwhite quail, *Colinus virginianus* | Fipronil | 11.3 (HT) | Tingle *et al.* (2003) |
| Feral pigeon, *Columba livia* | Fipronil | >2,000 (PNT) | Tingle *et al.* (2003) |
| Field sparrow, *Spizella pusilla* | Fipronil | 1,120 (ST) | Tingle *et al.* (2003) |
| Zebra finch, *Taeniopygia guttata* | Fipronil | 310 (MT) | Kitulagodage *et al.* (2008) |

Many of these studied species are granivorous and can be expected to feed on sown seeds shortly after the sowing period. Depending on crop species and consequent seed size, neonicotinoid-treated seeds can contain between 0.2–1 mg of active ingredient per seed. Goulson (2013) calculated that a granivorous grey partridge weighing 390 g would need to consume around five maize seeds, six sugar beet seeds or 32 oilseed rape seeds to receive a nominal LD_50_. Based on US Environmental Protection Agency estimates that around 1% of sown seed is accessible to foraging vertebrates at recommended sowing densities, Goulson calculated that sufficient accessible treated seed would be present to deliver a LD_50_ to ~100 partridges per hectare sown with maize or oilseed rape. Given that grey partridges typically consume around 25 g of seed a day there is the clear potential for ingestion of neonicotinoids by granivorous birds. However, no studies are available that demonstrate consumption of treated seed by farmland birds under field conditions or quantify relative consumption of treated versus untreated seed. More work is needed in this area to better understand total neonicotinoid exposure via this route.

In addition to lethal effects, several studies have identified sublethal effects of neonicotinoid ingestion on birds (Table 10). House sparrows can become uncoordinated and unable to fly, and studies of Japanese quail and red-legged partridges have reported DNA breakages and a reduced immune response, respectively. Many of these sublethal effects occur at lower concentrations than the lethal dose. A single oral dose of 41 mg/kg of imidacloprid will cause mortality in house sparrows, a substantially lower dose (6 mg/kg) can induce uncoordinated behaviour and an inability to fly (Cox 2001). While imidacloprid is highly toxic to Japanese quail, with an LD_50_ of 31 mg/kg, chronic daily doses of only 1 mg/kg/day can lead to testicular anomalies, DNA damage in males, and reductions in embryo size when those males are mated with control females (Tokumoto *et al.* 2013).

**Table 10.** Other studies of the direct effects of imidacloprid, clothianidin and fipronil on birds. Exposure could either be acute or chronic, the latter shown as /day (per day). All studies demonstrated deleterious effects at the given dosage, except those marked NE (no effect). Reproduced from Gibbons *et al.* (2015)

| Species | Effect on: | Imidacloprid | Clothianidin | Fipronil | Source and detailed effect |
| --- | --- | --- | --- | --- | --- |
| Mallard, *Anas platyrhynchos* | Reproduction | 16 mg/kg/day | >35 mg/kg/day (NE) |  | Adapted from figures in Mineau and Palmer (2013); various effects on reproduction |
| Chicken, *Gallus gallus domesticus* | Growth and development |  |  | 37.5 mg/kg | Kitulagodage *et al.* (2011a); reduced feeding and body mass, and developmental abnormalities of chicks |
| Chicken, *Gallus gallus domesticus* | Neurobehavioural |  |  | 37.5 mg/kg | Kitulagodage *et al.* (2011a); behavioural abnormalities of chicks |
| Red-legged partridge, *Alectoris rufa* | Survival | 31.9-53.4 mg/kg/day |  |  | Lopez-Antia *et al.* (2013); reduced chick survival at low dose, and reduced adult survival at high dose |
| Red-legged partridge, *Alectoris rufa* | Reproduction | 31.9 mg/kg/day |  |  | Lopez-Antia *et al.* (2013); reduced fertilisation rate and chick survival |
| Red-legged partridge, *Alectoris rufa* | Immunotoxic | 53.4 mg/kg/day |  |  | Lopez-Antia *et al.* (2013); reduced immune response |
| Northern bobwhite quail, *Colinus virginianus* | Reproduction |  | >52 mg/kg/day |  | Adapted from figures in Mineau and Palmer (2013); various effects on reproduction |
| Northern bobwhite quail, *Colinus virginianus* | Growth and development | 24 mg/kg/day<sup>a</sup> |  | 11 mg/kg<sup>b</sup> | <sup>a</sup>Adapted from figures in Mineau and Palmer (2013); various effects on weight<br><sup>b</sup>Kitulagodage *et al.* (2011b); birds stopped feeding so lost weight |
| Japanese quail, *Coturnix japonica* | Reproduction | 1 mg/kg/day |  |  | Tokumoto *et al.* (2013); testicular anomalies; reductions in embryo length when those males mated with un-dosed females |
| Japanese quail, *Coturnix japonica* | Genotoxic | 1 mg/kg/day |  |  | Tokumoto *et al.* (2013); increased breakage of DNA in males |
| House sparrow, *Passer domesticus* | Neurobehavioural | 6 mg/kg |  |  | Cox (2001); in-coordination, inability to fly |
| Zebra finch, *Taeniopygia guttata* | Reproduction |  |  | >1 mg/kg | Kitulagodage *et al.* (2011a); reduced hatching success |

In addition to the studies reviewed by Gibbons *et al.,* one additional study is available that assessed the impact of neonicotinoid ingestion on birds. Lopez-Anita *et al.* (2015) fed red-legged partridge ***Alectoris rufa*** imidacloprid-treated wheat seeds for a period of 25 days in the autumn and an additional period of 10 days in the spring, matching the pattern of cereal cropping in Spain. One treatment contained seeds treated at the recommended dosage rate and the second at 20% of the recommended rate, to mimic a diet comprised 20% of treated seeds. Treated seeds contained concentrations of imidacloprid of 0.14–0.7 mg/g at the two dose rates. As the 400 g partridges used in this study consume around 25 g of seeds a day, a daily ingestion of 8.8 and 44 mg/kg/day was expected, above the LD_50_ for Japanese quail (Table 9, SERA 2005).

Imidacloprid at the highest dose killed all adult partridges in 21 days, with first deaths occurring on day three. Mortality in the low dose and control groups was significantly lower at 18.7% and 15.6% respectively. As all partridges in the high dose died, effects on reproductive output were only measured in the low dose treatment. Compared to controls, low dose females laid significantly smaller clutches, and the time to first egg laying was also significantly increased. There was no difference in egg size, shell thickness, fertile egg rate and hatching rate. There was no detectable impact on chick survival, chick growth or sex ratio between these two groups. These results are in line with previous findings for lethal (Table 9) and sublethal (Table 10) effects of neonicotinoid consumption by birds. Whilst LD_50_s vary across two orders of magnitude from 11.3->2,000 mg/kg, sublethal effects are seen across a more consistent range of doses over one order of magnitude between 1–53 mg/kg. The greatest outstanding issue is that no data exist that quantify the actual exposure rate to granivorous birds from neonicotinoid-treated seeds. As such, it is difficult to judge whether these clearly demonstrated lethal and sublethal effects are manifested in wild bird populations in the field.

In addition to sublethal and lethal effects potentially caused by the ingestion of neonicotinoids from treated seeds, bird populations may also be affected by a reduction in invertebrate prey. Hallmann *et al.* (2014) used bird population data from the Dutch Common Breeding Bird Monitoring Scheme, a standardised recording scheme that has been running in the Netherlands since 1984. Surface water quality measurements are also regularly collected across the Netherlands, including data on imidacloprid levels. Hallmann *et al.* compared surface water imidacloprid levels between 2003–2009 with bird population trends for 15 farmland bird species that are insectivorous at least during the breeding season to assess the hypothesis that neonicotinoids may cause bird population declines through a reduction in invertebrate food availability. The average intrinsic rate of increase in local farmland bird populations was significantly negatively affected by the concentration of imidacloprid. At the individual level, 14 of the 15 bird species showed a negative response to imidacloprid concentrations, with 6 out of 15 showing a significant negative response. As previously discussed in Section 3.2, it is difficult to disentangle the effects of neonicotinoids from the effects of general agricultural intensification. Hallmann *et al.* attempt to control for proxy measures of intensification including changes in land use area, areas of cropped land and fertiliser input, but imidacloprid levels remained a significant negative predictor.

The only available study that has quantified changes in invertebrate prey availability after neonicotinoid treatment and concurrent changes in the bird community was conducted in the USA. Falcone and DeWald (2010) measured invertebrates in eastern hemlock *Tsuga canadensis* forests in Tennessee after trees has been treated with imidacloprid to control hemlock woolly adelgid *Adelges tsugae.* The imidacloprid treatment had a significantly negative effect on non-target Hemiptera and larval Lepidoptera. However, there was no corresponding decline in insectivorous bird density between treatments. Direct comparison between this study and the findings of Hallmann *et al.* 2014 are difficult due to the very different ecological conditions. It is likely sufficient untreated areas existed in hemlock forests for insectivorous birds to find sufficient forage. In the Netherlands, one of the most agriculturally intensified regions in the world, unaffected semi-natural habitat is scarce and a reduction in prey availability caused by neonicotinoid application would have a more severe impact.

No studies are available that measure the effect of neonicotinoids on bats and bat populations. A link between neonicotinoid use and declining farmland butterfly populations has been suggested (Gilburn *et al.* 2015; Forister *et al.* 2016) and given the ecological similarity between butterflies and moths a similar trend may be ongoing, though this has not yet been investigated. Many bat species feed on moths, so a reduction in the moth population is likely to impact bat populations through a reduction in food availability. Mason *et al.* (2014) link neonicotinoid use with an increase in the frequency of bat diseases such as White Nose Syndrome (caused by the fungus *Geomyces destructans)* in both the US and Europe. They hypothesise that consumption of neonicotinoid residues in insect prey weakens the immune system of bats. However, no evidence is presented demonstrating the presence of neonicotinoid residues in moths or bats, passage across these trophic levels or that exposure to neonicotinoids weaken the immune system of bats, resulting in increased rates of fungal infection. The position of Mason *et al.* must currently be considered unsupported.

### 3.6 Synergistic effects of additional pesticides with neonicotinoids

The EFSA (2013a; 2013b; 2013c) risk assessments for clothianidin, imidacloprid and thiamethoxam considered these pesticides and their impacts on honeybees individually. In the field, multiple neonicotinoids, other insecticides and other pesticides such as herbicides and fungicides are commonly applied to a single crop. Bees are frequently exposed to complex mixtures of pesticides, with 19 detected in trap caught bees from an agricultural region of Colorado (Hladik *et al.* 2016). It is possible that combinations of neonicotinoids and other pesticides may have antagonistic (become less effective), additive (equivalent to adding together existing effectiveness) or synergistic (multiplicative) effects. Morrissey *et al.* (2015) briefly listed known examples of synergistic effects between neonicotinoids and other pesticides. Several examples have been demonstrated by pesticide companies themselves. For example, Bayer demonstrated that the combination of clothianidin and the fungicide trifloxystrobin resulted in a 150-fold increase in kill rate to *Phaedon* leaf beetle larvae over clothianidin alone (Wachendorff-Neumann *et al.* 2012). Bayer scientists also demonstrated that treatments of 8,000 ppb of thiacloprid and 8,000 ppb of clothianidin resulted in aphid population kill rates of 25% and 0% after 6 days. Combining the two increased the kill rate to 98% (Andersch *et al.* 2010). Specifically for honeybees, Iwasa *et al.* (2004) demonstrated that the combination of thiacloprid with the fungicide propiconazole increased the toxicity of the mixture several hundred fold. Whilst synergies have been demonstrated, few environmental risk assessments have been made for neonicotinoids in combination with other pesticides.

Since 2013, a number of studies have investigated possible synergistic effects in neonicotinoids. Several have focussed on the interaction between neonicotinoids and ergosterol biosynthesis inhibitor (EBI) fungicides (which include propiconazole) and their impact on bees. Biddinger *et al.* (2013) studied the interaction between the contact toxicity of acetamiprid, imidacloprid and the fungicide fenbuconazole, a substance virtually non-toxic to bees (except at extremely high concentrations), using *A. mellifera* and Japanese orchard bees *Osmia cornifrons.* These pesticides are commonly found together in formulated products used in orchards. The doses ranged from 1.38–60 μg/bee 1:1 acetamiprid plus fenbuconazole mixture and 0.86–983 μg/bee 2:1 imidacloprid plus fenbuconazole mixture. At the LD_50_, the acetamiprid and fenbuconazole mixture was ~5 times more toxic than acetamiprid alone for *A. mellifera* and ~2 times more toxic than acetamiprid for *O. cornifrons.* However, these doses are exceptionally high, for example the 0.86 μg/bee imidacloprid:fenbuconazole mixture is equivalent to 567.6 ng/bee, with the *A. mellifera* contact toxicity to imidacloprid LD_50_ calculated as 81 ng/bee (Section 3.1). Unsurprisingly, this dose killed 85% of honeybee in this treatment. At unrealistically high concentrations it is not clear how informative these results are.

Thompson *et al.* (2014) investigated synergies between several EBI fungicides (flusilazole, propiconazole, myclobutanil and tebuconazole) and a range of neonicotinoids (clothianidin, thiacloprid, imidacloprid and thiamethoxam) on *A. mellifera.* Individual pesticides and mixtures of one neonicotinoid and one fungicide were administered through both contact and ingestion at a range of concentrations sufficient to increase mortality and bees were observed for a 96 hour period. LD_50_s were calculated after 48 hours as mortality did not significantly increase after this point. Single neonicotinoid and fungicide doses showed similar toxicity to previous published results, with no individual fungicide causing toxic effects even at concentrations of 22.4 μg/bee.

For neonicotinoid/fungicide mixtures, neonicotinoids were applied at calculated LD_50_s, in the region of 0.035–0.124 μg/bee for clothianidin, imidacloprid and thiamethoxam and 122.4 μg/bee for thiacloprid (cyano-substituted neonicotinoids having lower toxicity to bees, Section 3.1.1). Fungicides were applied at doses of between 0.161 and 0.447 μg/bee depending on the particular compound. These values of were calculated as realistic worst-case exposures based on approved application rates for UK crops. For these mixtures, a synergy ratio was calculated where the LD_50_ of the neonicotinoid was divided by the LD_50_ of the neonicotinoid plus fungicide mixture. Consequently, a value of over one indicates the mixture was more toxic and a value under one indicates the mixture was less toxic. Combinations of fungicides with thiacloprid and clothianidin showed negligible synergy for contact toxicity, with an average synergism ratio of 0.30 and 1.07 respectively. Imidacloprid and thiamethoxam were higher at 1.53 and 2.02. For oral toxicity, thiacloprid and imidacloprid showed low synergy at 0.60 and 0.48 whereas clothianidin and thiamethoxam were higher at 1.52 and 1.31 respectively. Only two combinations showed significant synergy, for a contact dose of tebuconazole and thiamethoxam with a synergy of 2.59 and for an oral dose of clothianidin and tebuconazole at a synergy of 1.90.

Sgolastra *et al.* (2016) investigated the interaction between clothianidin and the fungicide propiconazole in three bee species, *A. mellifera, B. terrestris* and *O. bicornis*. Each species was administered a LD_10_ dose of clothianidin (0.86, 1.87 and 0.66 ng/bee respectively, see Section 3.1.1 for more detail), a non-lethal dose of propiconazole (7 μg/bee) and a combination of the two treatments. Bees were then observed for a 96 hour period and mortality quantified. Some synergistic effects were seen. In *A. mellifera*, mortality was significantly higher for the combined dose in the first two time periods (4 and 24 hours). Mortality in *B. terrestris* for the combined dose was only significantly higher in the first time period, after 4 hours. However, in *O. bicornis*, exposure to the combination of clothianidin and propiconazole resulted in significantly higher mortality at all time points (Figure 13).

**Figure 13.**
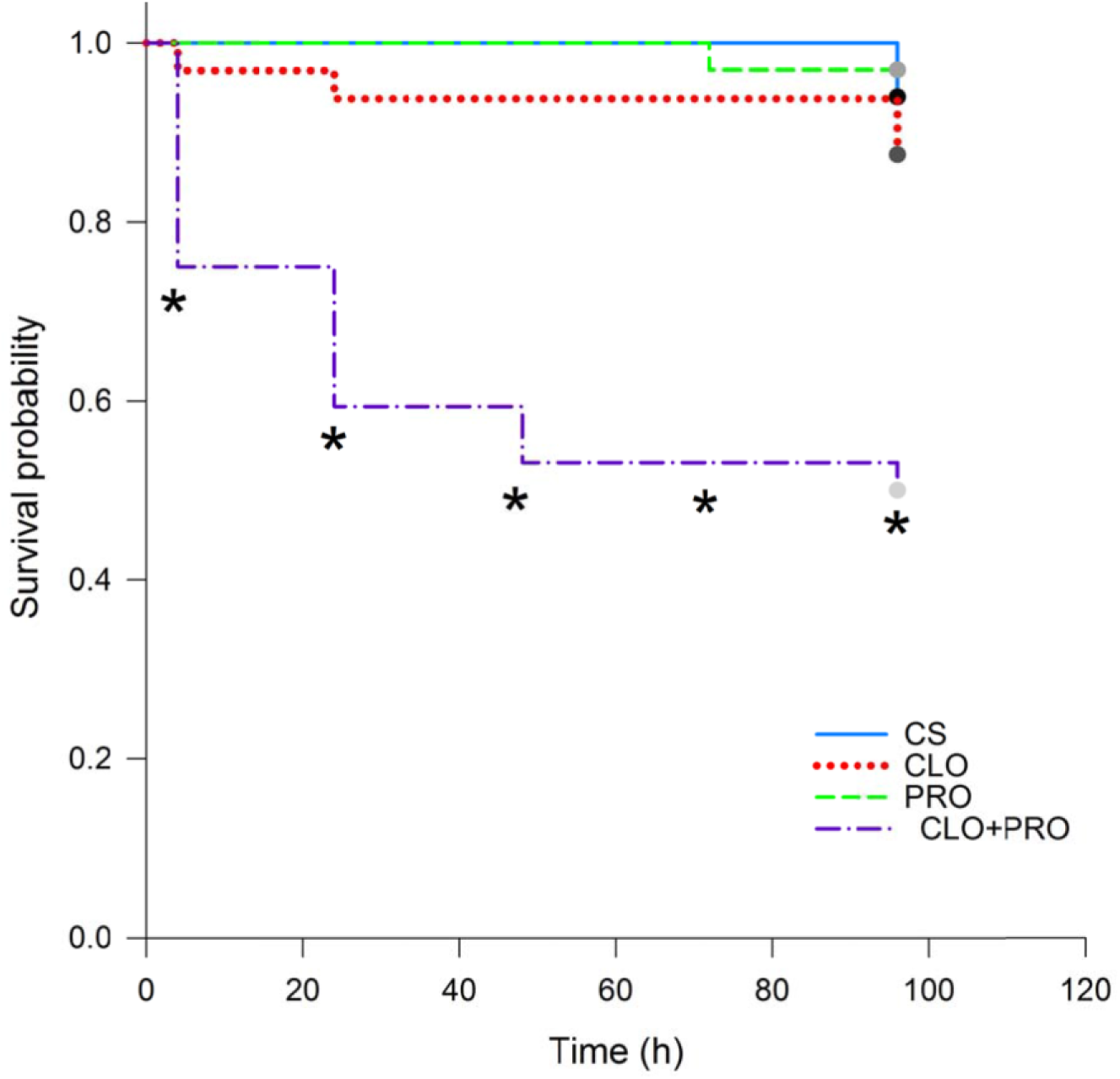
Cumulative proportion of surviving *Osmia bicornis* females exposed to a control solution (CS – sugar water solution with 3% acetone), clothianidin (CLO – 0.63 ng/bee) propiconazole (PRO – 7 μg/bee), and clothianidin plus propiconazole (CLO+PRO – 0.63 ng/bee plus 7 μg/bee). Statistically significant synergistic effects at the various assessment times (4, 24, 48, 72, 96 h) are marked with an asterisk.

Spurgeon *et al.* (2016) conducted similar experiments to Sgolastra *et al.,* investigating the effect of a combination of clothianidin and propiconazole on *A. mellifera, B. terrestris* and *O. bicornis*. In order to calculate an LD_50_, clothianidin concentrations were varied and propiconazole concentrations were held at zero, a low dose and a high dose. The low dose was taken from the EFSA Panel on Plant Protection Products (2012) reported environmental concentrations, and the high dose was 10 times the low dose to represent a plausible worst case scenario, but it is not clear what these values actually are. Mortality was quantified over 48, 96 and 240 hours. For *A. mellifera*, clothianidin LC_50_s with and without propiconazole were always within a factor of 2, with no clear negative trend at higher propiconazole concentrations. For *B. terrestris*, clothianidin LC_50_s with propiconazole were between 1.5 to 2 fold lower. For *O. bicornis*, clothianidin LC_50_s with propiconazole was up to 2 fold lower with a negative trend as propiconazole concentrations increased. Spurgeon *et al.* concluded that the clothianidin and propiconazole combination had no to slight synergy for *A. mellifera* and slight to moderate synergy for *B. terrestris* and *O. bicornis*.

In an additional trial, Thompson *et al.* (2014) demonstrated that the dose of fungicide applied is a key factor determining neonicotinoid toxicity using propiconazole and thiamethoxam mixtures (Table 11). The authors argue that their low rates of significant synergies between neonicotinoids and fungicides was because of their lower, more field-realistic fungicide doses of 0.161–0.447 μg/bee compared to 10 μg/bee used by Iwasa *et al.* (2004), an early study demonstrating this interaction. The values of 0.161–0.447 μg/bee were calculated as realistic worst-case exposures based on approved application rates for UK crops. However, data are lacking demonstrating true field-realistic exposure rates to fungicides for free flying bees. Whilst studies such as Sgolastra *et al.* (2016) show a clear synergistic effect between fungicides and neonicotinoids on *O. bicornis,* the dose of fungicide used is more than an order of magnitude greater than that used by Thompson *et al.* Bees are consistently exposed to fungicides with 40 types found in honeybee pollen, wax and nectar (Sánchez-Bayo and Goka 2014). Pollen collected by bumblebees and stored in their nests has also been found to contain fungicides at average concentrations between 0.15–25 ppb (EBI fungicides 0.15–17 ppb, David *et al.* 2016). However, almost nothing is known about how concentrations present in bee-collected material translate into acute or chronic exposure to bees. It is currently not possible to comment on what fungicide doses represent a realistic situation that bees are likely to encounter in the wild.

**Table 11.** Comparison of the ratio of propiconazole to the doses of thiamethoxam and the resultant LD_50_ in the contact and oral studies. Synergy ratios marked with an * were significantly different. Reproduced from Thompson *et al.* (2014).

| Contact dose<br>propiconazole<br>µg/bee | Ratio fungicide:<br>thiamethoxam<br>contact LD<sub>50</sub> | Contact LD<sub>50</sub><br>thiamethoxam<br>µg/bee | Synergy<br>ratio | Ratio fungicide:<br>thiamethoxam<br>oral LD<sub>50</sub> | Oral LD<sub>50</sub><br>thiamethoxam<br>µg/bee | Synergy<br>ratio |
| --- | --- | --- | --- | --- | --- | --- |
| 0 | - | 0.0373 | - | - | 0.0641 | - |
| 0.0224 | 0.6 | 0.0288 | 1.3 | 0.349 | 0.0268 | 2.4 |
| 0.224 | 6 | 0.0247 | 1.5 | 3.49 | 0.0277 | 2.3 |
| 2.24 | 60 | 0.0134 | 2.8* | 34.9 | 0.0265 | 2.4 |
| 22.4 | 600 | 0.0104 | 3.6* | 349 | 0.00776 | 8.3* |

In addition to work on bees, Kunce *et al.* (2015) investigated the impact of one hour pulse exposure of imidacloprid and thiamethoxam and two pyrethroids, deltamethrin and esfenvalerate in single, pairwise and combined doses on the development of the aquatic midge *C. riparius* (see Section 3.4 for more methodological and concentration details). Most pesticide treatments reduced the survival of the larvae, but the deleterious effects did not appear to be synergistically amplified by a combination of pesticides. Kunce *et al.* conclude that at the low doses and period of exposure used, the risk of synergistic or additive effects is very low. Much more work on the potential synergistic effects of pesticides in aquatic ecosystems is required.

Overall, these studies support the position that neonicotinoids can act synergistically with fungicides, increasing their lethality to bees. However, the dose rate of both neonicotinoids and fungicides, time of exposure, neonicotinoid and fungicide chemical class and length of time after exposure are all important explanatory factors affecting this relationship. The concentration of fungicide used in laboratory studies appears to be the most important factor determining synergistic lethality. Fungicides are regularly sprayed during the period when flowering crops are in bloom under the assumption that these compounds are safe for bees. Further work is needed in this area to establish realistic levels of fungicide exposure for free flying bees in order to assess the likely impact of neonicotinoid/fungicide synergies on bee populations.

Studies to date have only examined pairwise interactions between pesticides. It is clear that bees and other non-target organisms inhabiting farmland are routinely exposed to far more complex cocktails of pesticides than any experimental protocol has yet attempted to examine. For example, honeybee and bumblebee food stores commonly contain 10 or more pesticides (e.g. David *et al.* 2016). A major challenge for scientists and regulators is to attempt to understand how chronic exposure to complex mixtures of neonicotinoids and other chemicals affects wildlife.

## 4. CONCLUDING REMARKS

### 4.1 Advances in scientific understanding and comparison with the 2013 knowledge base

The EFSA reports into clothianidin, imidacloprid and thiamethoxam are naturally narrow in scope, focusing specifically on the risks that these neonicotinoids pose to bees, with almost all data consisting of and referring to the honeybee *Apis mellifera*. Because the scope of this review is much wider, focusing on neonicotinoid persistence in the wider environment and possible impacts on many non-target organisms, a simple comparison with the EFSA reports is not possible as there is no well-defined baseline of existing knowledge prior to 2013 for most topic areas. However, it is possible to comment on the change in the scientific evidence since 2013 compared to the EFSA reports. This process is not meant to be a formal assessment of the risk posed by neonicotinoids in the manner of that conducted by EFSA. Instead it aims to summarise how the new evidence has changed our understanding of the likely risks to bees; is it lower, similar or greater than the risk perceived in 2013. With reference to the EFSA risk assessments baseline, advances in each considered area and their impact on the original assessment can be briefly summarised thus:

- *Risk of exposure from pollen and nectar of treated flowering crops*. The EFSA reports calculated typical exposure from flowering crops treated with neonicotinoids as seed dressings. Considerably more data are now available in this area, with new studies broadly supporting the calculated exposure values. For bees, flowering crops pose a **Risk Unchanged** to that reported by EFSA 2013a.
- *Risk from non-flowering crops and cropping stages prior to flowering*. Non-flowering crops were considered to pose no risk to bees. No new studies have demonstrated that these nonflowering crops pose a direct risk to bees. They remain a **Risk Unchanged**.
- *Risk of exposure from the drilling of treated seed and subsequent dust drift*. Despite modification in seed drilling technology, available studies suggest that dust drift continues to occur, and that dust drift still represents a source of acute exposure and so is best considered a **Risk Unchanged**.
- *Risk of exposure from guttation fluid*. Based on available evidence this was considered a low-risk exposure path by EFSA 2013a. New data have not changed this position and so it remains a **Risk Unchanged**.
- *Risk of exposure from and uptake of neonicotinoids in non-crop plants*. Uptake of neonicotinoids by non-target plants was considered likely to be negligible, though a data gap was identified. Many studies have since been published demonstrating extensive uptake of neonicotinoids and their presence in the pollen, nectar and foliage of wild plants, and this source of exposure may be much more prolonged than the flowering period of the crop. Bees collecting pollen from neonicotinoid-treated crops can generally be expected to be exposed to the highest neonicotinoid concentrations, but non-trivial quantities of neonicotinoids are also present in pollen and nectar collected from wild plants. Exposure from non-target plants clearly represents a **Greater Risk**.
- *Risk of exposure from succeeding crops*. A data gap was identified for this issue. Few studies have explicitly investigated this, but this area does represent some level of risk as neonicotinoids and now known to have the potential to persist for years in the soil, and can be detected in crops multiple years after the last known application. However, as few data exist this is currently considered a **Risk Unchanged**.
- *Direct lethality of neonicotinoids to adult bees*. Additional studies on toxicity to honeybees have supported the values calculated by EFSA. More data have been produced on neonicotinoid toxicity for wild bee species and meta-analyses suggest a broadly similar response. Reference to individual species is important but neonicotinoid lethality should be broadly considered a **Risk Unchanged**.
- *Sublethal effects of neonicotinoids on wild bees*. Consideration of sublethal effects by EFSA was limited as there is no agreed testing methodology for the assessment of such effects. A data gap was identified. Exposure to neonicotinoid-treated flowering crops has been shown to have significant negative effects on free flying wild bees under field conditions and some laboratory studies continue to demonstrate negative effects on bee foraging ability and fitness using field-realistic neonicotinoid concentrations. **Greater Risk**.

Within this context, research produced since 2013 suggest that neonicotinoids pose a similar to greater risk to wild and managed bees, compared to the state of play in 2013. Given that the initial 2013 risk assessment was sufficient to impose a moratorium on the use of neonicotinoids on flowering crops, and given that new evidence either confirms or enhances evidence of risk to bees, it is logical to conclude that the current scientific evidence supports the extension of the moratorium.

In addition to the use of neonicotinoids on flowering crops, research since 2013 has demonstrated neonicotinoid migration into and persistence in agricultural soils, waterways and constituent parts of non-crop vegetation. Where assessments have been made of concentrations likely to significantly negatively affect non-target organisms, levels have been demonstrated to be above these thresholds in numerous non-crop agricultural habitats.

The strongest evidence for this is found in waterbodies surrounding agricultural areas, both temporary and permanent. The impact of neonicotinoids on aquatic organisms appears to be the easiest to quantify, as field-realistic concentrations can be easily obtained through sample collection and once neonicotinoids are present in waterbodies, aquatic organisms cannot limit their exposure to them. In contrast, assessing the field-realistic exposure of bees to neonicotinoids is much harder, as it will depend on numerous factors including but not limited to: the type of flowering crop, its relative attractiveness compared to existing available forage, the crop type and levels of neonicotinoid loss into the wider environment through seed dust and leaching, soil type and organic content and consequent retention of neonicotinoid active ingredient, uptake of neonicotinoids by surrounding vegetation and relative collection of pollen and nectar from various wild plants containing variable levels of neonicotinoids at different parts of the year. In addition, wild and managed bees have traits such as flight period, floral choice preferences and social structure that vary radically between different bee species, as can be clearly seen in the three most commonly used bee model organisms *A. mellifera*, *B. terrestris* and *O. bicornis*. As such, it is much more difficult to gain a completely accurate and consistent measure of neonicotinoid exposure for taxa such as these.

However, whilst these aforementioned factors are all important, it is still possible to comment on likely outcomes based on average exposure levels across a range of studies. This is as true for other taxa as it is for bees. Given these caveats, it is clear that since 2013, new research has substantially advanced our understanding of the effect of neonicotinoids on non-target organisms in the following areas:

- Non-flowering crops treated with neonicotinoids can pose a risk to non-target organisms through increasing mortality in beneficial predator populations.
- Neonicotinoids can persist in agricultural soils for several years, leading to chronic contamination and, in some instances, accumulation over time.
- Neonicotinoids continue to be found in a wide range of different waterways including ditches, puddles, ponds, mountain streams, rivers, temporary wetlands, snowmelt, groundwater and in outflow from water processing plants.
- Reviews of the sensitivity of aquatic organisms to neonicotinoids show that many aquatic insect species are several orders of magnitude more sensitive to these compounds than the traditional model organisms used in regulatory assessments for pesticide use.
- Neonicotinoids have been shown to be present in the pollen, nectar and foliage of non-crop plants adjacent to agricultural fields. This ranges from herbaceous annual weeds to perennial woody vegetation. We would thus expect non-target herbivorous insects and non-bee pollinators inhabiting field margins and hedgerows to be exposed to neonicotinoids. Of particular concern, this includes some plants sown adjacent to agricultural fields specifically for the purposes of pollinator conservation.
- Correlational studies have suggested a link between neonicotinoid usage in agricultural areas and population metrics for butterflies, bees and insectivorous birds in three different countries.

### 4.2 Existing knowledge gaps and future research

Whilst much research has been conducted on neonicotinoid pesticides and their impact on non-target organisms since 2013, a number of key knowledge gaps exist. As stated by Godfray *et al.* (2015) in their update on the existing scientific literature concerning neonicotinoids and insect pollinators, it is important to remember that major gaps in our understanding occur and different policy conclusions can be drawn depending on the weight given to important (but not definitive) scientific findings and the economic and other interests of different stakeholders. This review is not intended as a risk assessment, simply as a review of advances in our scientific understanding of the environmental risks that neonicotinoids pose.

From the perspective of better understanding the impacts of neonicotinoids on non-target organisms, further research is needed in the following areas:

- Whilst the impact of neonicotinoids on bees have been relatively well studied, few data exist for most taxa. The sensitivity of non-pest herbivorous taxa and important natural enemies of crop pests to neonicotinoids are particularly poorly understood.
- Continue to improve our understanding of realistic neonicotinoid and other pesticide exposure in agricultural and non-agricultural areas for understudied taxa. The implications of laboratory studies assessing the lethal and sublethal impacts of neonicotinoids are unclear without a realistic baseline for comparison with real world conditions. Data are most lacking for herbivorous, soil dwelling, parasitic and predatory invertebrates and granivorous and insectivorous terrestrial vertebrates.
- In addition to sensitivity and exposure, the movement of neonicotinoids through trophic levels is poorly understood with the exception of a few field studies which demonstrate the principle. Some authors have linked direct neonicotinoid exposure with declines in higher trophic level organisms, but little to no data exist regarding these claims.
- Long-term datasets exist that have demonstrated recent population declines across various taxa, with the most pronounced declines correlating with neonicotinoid use. Whilst these studies are suggestive in their own right, the effects of general agricultural intensification relative to the effects of neonicotinoid pesticides must be teased apart if long term declines in taxa are to be better understood and reversed.
- Possible synergistic and additive effects of neonicotinoids with other pesticides are still poorly understood for bees, and almost nothing is known about their effects on other non-target taxa. This problem is compounded by a lack of understanding of field-realistic exposures to the various constituent active ingredients, with different taxa likely to be receiving different doses depending on their interaction with agricultural environments.

### 4.3 Closing statement

Recent work on neonicotinoids continues to improve our understanding of how these compounds move through and persist in the wider environment. These water soluble compounds are not restricted to agricultural crops, instead permeating most parts of the agricultural environments in which they are used and in some cases reaching further afield via waterways and runoff water. Field-realistic laboratory experiments and field trials continue to demonstrate that residual neonicotinoid traces can have a mixture of lethal and sublethal effects on a wide range of taxa. Relative to the risk assessments produced in 2013 for clothianidin, imidacloprid and thiamethoxam which focussed on their effects on bees, new research strengthens arguments for the imposition of a moratorium on their use, in particular because it has become evident that they pose significant risks to many non-target organisms, not just bees. Given the improvement in scientific knowledge of how neonicotinoids move into the wider environment from all crop types, a discussion on the risks posed by their use on non-flowering crops and in non-agricultural areas is urgently needed.

